# Rapid and independent evolution of ancestral and novel defenses in a genus of toxic plants (*Erysimum*, Brassicaceae)

**DOI:** 10.1101/761569

**Authors:** Tobias Züst, Susan R. Strickler, Adrian F. Powell, Makenzie E. Mabry, Hong An, Mahdieh Mirzaei, Thomas York, Cynthia K. Holland, Pavan Kumar, Matthias Erb, Georg Petschenka, José María Goméz, Francisco Perfectti, Caroline Müller, J. Chris Pires, Lukas A. Mueller, Georg Jander

## Abstract

Phytochemical diversity is thought to result from coevolutionary cycles as specialization in herbivores imposes diversifying selection on plant chemical defenses. Plants in the speciose genus *Erysimum* (Brassicaceae) produce both ancestral glucosinolates and evolutionarily novel cardenolides as defenses. Here we test macroevolutionary hypotheses on co-expression, co-regulation, and diversification of these potentially redundant defenses across this genus. We sequenced and assembled the genome of *E. cheiranthoides* and foliar transcriptomes of 47 additional *Erysimum* species to construct a highly resolved phylogeny, revealing that cardenolide diversity increased rapidly rather than gradually over evolutionary time. Concentrations, inducibility, and diversity of the two defenses varied independently among species, with no evidence for trade-offs. Closely related species shared similar cardenolide traits, but not glucosinolate traits, likely as a result of specific selective pressures acting on distinct molecular diversification mechanisms. Ancestral and novel chemical defenses in *Erysimum* thus appear to provide complementary rather than redundant functions.

## Introduction

Plant chemical defenses play a central role in the coevolutionary arms race with herbivorous insects. In response to diverse environmental challenges, plants have evolved a plethora of structurally diverse organic compounds with repellent, antinutritive, or toxic properties (Fraenkel 1959, Mithöfer and Boland 2012). Chemical defenses can impose barriers to consumption by herbivores, but in parallel may favor the evolution of specialized herbivores that can tolerate or disable these defenses (Cornell and Hawkins 2003). Chemical diversity is likely evolving in response to a multitude of plant-herbivore interactions (Salazar et al. 2018), and community-level phytochemical diversity may be a key driver of niche segregation and insect community dynamics (Richards et al. 2015, Sedio et al. 2017).

For individual plants, the production of diverse mixtures of chemicals is often considered advantageous (Romeo et al. 1996, Firn and Jones 2003, Gershenzon et al. 2012, Forbey et al. 2013, Richards et al. 2016). For example, different chemicals may target distinct herbivores (Iason et al. 2011, Richards et al. 2015), or may act synergistically to increase overall toxicity of a plant (Steppuhn and Baldwin 2007). However, metabolic constraints can limit the extent of phytochemical diversity within individual plants (Firn and Jones 2003). Most defensive metabolites originate from a small group of precursor compounds and conserved biosynthetic pathways, which are modified in a hierarchical process into diverse, species-specific end products (Moore et al. 2014). As constraints are likely strongest for the early stages of these pathways, related plant species commonly share the same functional ‘classes’ of defensive chemicals (Wink 2003), but vary considerably in the number of compounds within each class (Fahey et al. 2001, Rasmann and Agrawal 2011).

Functional conservatism in defensive chemicals among related plants should facilitate host expansion and the evolution of tolerance in herbivores (Cornell and Hawkins 2003), as specific adaptations to deactivate or detoxify one compound are more likely to be effective against structurally similar than structurally dissimilar compounds. This may result in a seemingly paradoxical scenario, wherein well-defended plants are nonetheless attacked by a diverse community of specialized herbivores (Agrawal 2005, Bidart-Bouzat and Kliebenstein 2008). For example, most plants in the Brassicaceae produce glucosinolates as their primary defense, which upon activation by myrosinase (thioglucoside glucohydrolase) enzymes at leaf damage become potent repellents of many herbivores (Fahey et al. 2001). However, despite the potency of this defense system and the large diversity of glucosinolates produced by the Brassicaceae, several specialized herbivores have evolved strategies to overcome this defense, enabling them to consume most Brassicaceae and even to sequester glucosinolates for their own defense against predators (Müller 2009, Winde and Wittstock 2011).

Plants may occasionally overcome the constraints on functional diversification and gain the ability to produce new classes of defensive chemicals as a ‘second line of defense’ (Feeny 1977). Although this phenomenon is likely widespread across the plant kingdom, it has most commonly been reported from the well-studied Brassicaceae. In addition to producing evolutionarily ancestral glucosinolates, plants in this family have gained the ability to produce saponins in *Barbarea vulgaris* (Shinoda et al. 2002), alkaloids in *Cochlearia officinalis* (Brock et al. 2006), cucurbitacins in *Iberis* spp. (Nielsen 1978b), alliarinoside in *Alliaria petiolata* (Frisch and Møller 2012), and cardenolides in the genus *Erysimum* (Makarevich et al. 1994). These recently-evolved chemical defenses with modes of action distinct from glucosinolates have likely allowed the plants to escape attack from specialized, glucosinolate-adapted herbivores (Nielsen 1978b, Dimock et al. 1991, Haribal and Renwick 2001, Shinoda et al. 2002). Gains of novel defenses are expected to result in a release from selective pressures imposed by specialized antagonists, and thus may represent key steps in herbivore-plant coevolution that lead to rapid phylogenetic diversification (Weber and Agrawal 2014).

The production of cardenolides by species in the genus *Erysimum* is one of the longest- and best-studied examples of an evolutionarily recent gain of a novel chemical defense (Jaretzky and Wilcke 1932, Nagata et al. 1957, Singh and Rastogi 1970, Makarevich et al. 1994). Cardenolides are a type of cardiac glycoside, which act as allosteric inhibitors of Na^+^/K^+^-ATPase, an essential membrane ion transporter that is expressed ubiquitously in animal cells (Agrawal et al. 2012). Cardiac glycosides are produced by plants in approximately sixty genera belonging to twelve plant families, and several cardiac glycoside-producing plants are known for their toxicity or medicinal uses (Agrawal et al. 2012, Züst et al. 2018). *Erysimum* is a species-rich genus consisting of diploid and polyploid species with diverse morphologies, growth habits, and ecological niches (Al-Shehbaz 1988, Polatschek and Snogerup 2002, Al-Shehbaz 2010, Gómez et al. 2015). Of the *Erysimum* species evaluated to date, all produced some of the novel cardenolide defenses (Makarevich et al. 1994). Previous phylogenetic studies suggest a recent and rapid diversification of the genus, with estimates of the onset of radiation ranging between 0.5 and 2 million years ago (Gómez et al. 2014, Moazzeni et al. 2014), and of 150 to 350 extant species (Polatschek and Snogerup 2002, Al-Shehbaz 2010). The large uncertainty in species number reflects taxonomic challenges in this genus, which includes many species that readily hybridize, as well as cryptic species with near-identical morphology (Abdelaziz et al. 2011).

In most *Erysimum* species, cardenolides appear to have enabled an escape from at least some glucosinolate-adapted specialist herbivores. Cardenolides in *Erysimum* act as oviposition and feeding deterrents for different pierid butterflies (Chew 1975, 1977, Wiklund and Åhrberg 1978, Renwick et al. 1989, Dimock et al. 1991), and several glucosinolate-adapted beetles (*Phaedon* spp. and *Phyllotreta* spp.) were deterred from feeding by dietary cardenolides at levels commonly found in *Erysimum* (Nielsen 1978a, b). Nonetheless, *Erysimum* plants are still attacked by a range of herbivores and seed predators, including some mammals and several glucosinolate-adapted aphids, true bugs, and lepidopteran larvae (Gómez 2005, Züst et al. 2018). Despite their potency, cardenolides thus do not provide a universal defense.

The gain of a novel chemical defense makes the genus *Erysimum* an excellent model system to study the causes and consequences of phytochemical diversification (Züst et al. 2018). While an increasing number of studies are beginning to describe taxon-wide patterns of chemical diversity in plants (e.g., Richards et al. 2015, Sedio et al. 2017, Salazar et al. 2018), the *Erysimum* system is unique in combining two classes of plant metabolites with primarily defensive function – although a broader role of glucosinolates is increasingly recognized (e.g., Katz et al. 2015). The system thus is ideally suited to evaluate the evolutionary consequences of co-expressing two functionally distinct but potentially redundant defenses. Here, we present a high-quality genome sequence assembly and annotation for the short-lived annual *E. cheiranthoides* as an important resource for future molecular studies in this system. Furthermore, we present a highly resolved phylogeny for 47 additional species constructed from transcriptome sequences, corresponding to 10-30% of species in the genus *Erysimum*. We combine this phylogeny with a characterization of the full diversity of glucosinolates and cardenolides in leaves to evaluate macroevolutionary patterns in the evolution of phytochemical diversity across the genus. We complemented the characterization of defensive phenotypes by quantifying glucosinolate-activating myrosinase activity, inhibition of animal Na^+^/K^+^-ATPase by leaf extracts, and defense inducibility in response to exogenous application of jasmonic acid (JA). By assessing co-variation of diversity, abundance and inducibility of ancestral and novel defenses, we provide evidence that the two defense metabolite classes evolved in response to different selective pressures and appear to serve specific, non-redundant roles.

## Materials and Methods

### Plant material and growth conditions

The genus *Erysimum* is distributed across the northern hemisphere, with the center of diversity stretching from the Mediterranean Basin into Central Asia, and a smaller number of species centered in western North America (Moazzeni et al. 2014). Seeds of *Erysimum* species spanning a range of distributions in Europe and Western North America were collected in their native habitats or obtained from botanical gardens and commercial seed suppliers (Figure 1, Table S1). Ploidy levels of species were inferred from literature reports to test for the effect of ploidy on chemical diversity (Table S1). For seeds obtained from botanical gardens, we mostly used species names as provided by the supplier. As an exception, seeds of *E. collinum* (COL) had originally been designated as *E. passgalense*, but these species names are now considered as synonymous (German 2014). Furthermore, plants of four seed batches did not exhibit the expected phenotypes and likely were the result of seed mislabeling by the suppliers; we nonetheless included these plants for transcriptome sequencing, but refer to them as accessions ER1, ER2, ER3, and ER4 (Table S1). For genome sequencing of *E. cheiranthoides*, seeds that were collected from a natural population in the Elbe River floodplain (Germany, Figure 1) were planted in a greenhouse in one-liter pots in Cornell mix (by weight 56% peat moss, 35% vermiculite, 4% lime, 4% Osmocote slow-release fertilizer [Scotts, Marysville, OH], and 1% Unimix [Scotts]). This lineage, which we have designated “*Elbtalaue*”, was propagated by self-pollination and single-seed descent for six generations prior to further experiments.

**Figure. 1.**
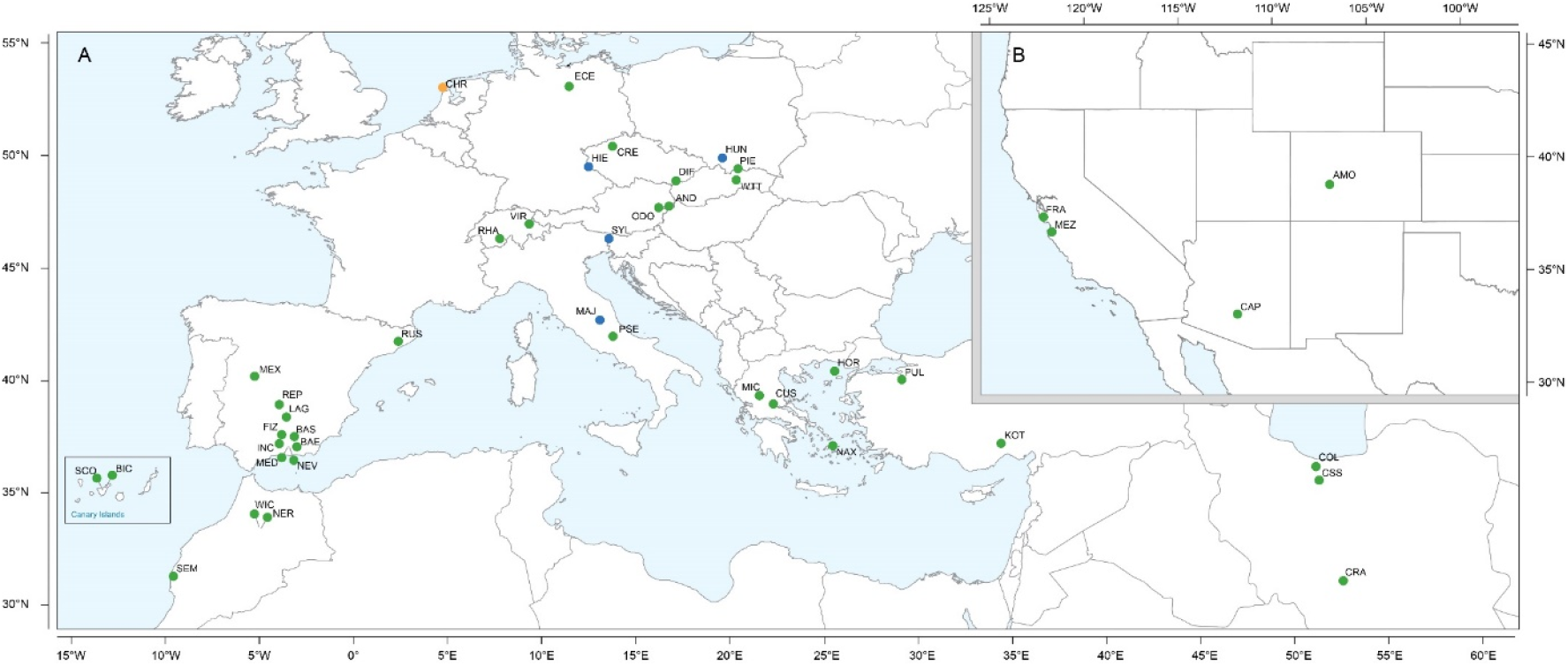
Geographic location of *Erysimum* spp. source populations in Europe (A) and North America (B). Inset: The Canary Islands (28°N, 16°W) are located further westward and southward than drawn in this map. Green symbols are exact collection locations, while blue symbols indicate approximate locations based on species distributions. Seeds of the originally Mediterranean species *E. cheiri* (CHR, orange symbol) were collected from a naturalized population in the Netherlands. Five species/accessions (ALI, ER1, ER2, ER3, ER4) could not be placed on the map due to uncertain species identity.

For transcriptome sequencing and metabolomic analyses of *Erysimum* species, subsets of the full species pool were grown in three separate experiments in 2016 and 2017. While some species were included in all three experiments, others could only be grown once due to limited seed availability or germination. To maximize germination success, seeds were placed on water agar (1%) in Petri dishes and cold-stratified for two weeks. After stratification, Petri dishes were moved to a growth chamber set to 24 °C day / 22 °C night at a 16:8 h photoperiod. Viable seeds germinated within 3-10 days of placement in the growth chamber. As soon as cotyledons had fully extended, we transplanted the seedlings into 10 x 10 cm plastic pots filled with a mixture of peat-based germination soil (Seedlingsubstrat, Klasmann-Deilmann GmbH, Geeste, Germany), field soil, sand, and vermiculite at a ratio of 6:3:1:5. Plants were moved to a climate-controlled greenhouse set to 24 °C day / 16 °C night and 60 % RH with natural light and supplemented artificial light set to a 14:10 h photoperiod. Plants were watered as needed throughout the experiments, and fertilized with a single application of 0.1 L of fertilizer solution (N:P:K 8:8:6, 160 ppm N) three weeks after transplanting.

### Erysimum cheiranthoides genome and transcriptome sequencing

DNA sequencing for genome assembly and RNA sequencing for annotation were conducted with samples prepared from sixth-generation inbred *E. cheiranthoides* var. *Elbtalaue*. High molecular weight genomic DNA was extracted from the leaves of a single *E. cheiranthoides* plant using Wizard® Genomic DNA Purification Kit (Promega, Madison WI, USA). The quantity and quality of genomic DNA was assessed using a Qubit 3 fluorometer (Thermo Fisher, Waltham, MA, USA) and a Bioanalyzer DNA12000 kit (Agilent, Santa Clara, CA, USA). Twelve µg of non-sheared DNA were used to prepare the SMRTbell library, and the size-selection of 15-50 kb was performed on Sage BluePippin (Sage Science, Beverly, MA, USA) following manufacturer’s instructions (Pacific Biosciences, Menlo Park, CA, USA) and as described previously (Chen et al. 2019). PacBio sequencing was performed by the Sequencing and Genomic Technologies Core of the Duke Center for Genomic and Computational Biology (Durham, NC, USA). For genome polishing, one DNA library was prepared using the PCR-free TruSeq DNA sample preparation kit following the manufacturer’s instructions (Illumina, San Diego, CA), and sequenced on an Illumina MiSeq instrument (paired-end 2×250bp) at the Cornell University Biotechnology Resource Center (Ithaca, NY).

The transcriptome of sixth-generation inbred *E. cheiranthoides* var. *Elbtalaue* plants was sequenced using both PacBio (Iso-Seq) and Illumina sequencing methods. Total RNA was isolated from stems, flowers, buds, pods, young and mature leaves of five plants (siblings of the plant used for genome sequencing) using the SV Total RNA Isolation Kit with on-column DNase I treatment (Promega, Madison, WI, USA). The RNA quantity and quality were assessed by RIN (RNA Integrity Number) using a 2100 Bioanalyzer (Agilent Technologies, Santa Clara, CA). The samples with a RIN value of >7 were pooled across all six tissue types. One µg of the pooled total RNA was used for the Iso-Seq following the manufacturer’s instructions (Iso-Seq^TM^). The library preparation and sequencing were performed by Sequencing and Genomic Technologies Core of the Duke Center for Genomic and Computational Biology (Durham, NC, USA). For Illumina sequencing, 2 μg of purified pooled total RNA from three replicates was used for the preparation of strand-specific RNAseq libraries with 14 cycles of final amplification (Zhong et al 2012). The purified libraries were multiplexed and sequenced with 101 bp paired-end read length in two-lanes on an Illumina HiSeq2500 instrument (Illumina, San Diego, CA) at the Cornell University Biotechnology Resource Center (Ithaca, NY). For Hi-C scaffolding, 500 mg of *E. cheiranthoides* leaf tissue was flash-frozen and sent to Phase Genomics (Phase Genomics Inc. Seattle, WA, USA).

### E. cheiranthoides genome assembly and gene annotation

PacBio sequences from the genome of *E. cheiranthoides* were assembled using Falcon (Chin et al. 2016). The assembly was polished using Arrow from SMRT Analysis v2.3.0 (https://www.pacb.com/products-and-services/analytical-software/smrt-analysis/) with PacBio reads, and then assembled into chromosome-scale scaffolds using Hi-C methods by Phase Genomics (Seattle, WA, USA). Scaffolding gaps were filled with PBJelly v13.10 (English et al. 2012) using PacBio reads followed by three rounds of Pilon v1.23 correction (Walker et al. 2014) with 9 Gbp of Illumina paired-end 2 x 150 reads. BUSCO v3 (Waterhouse et al. 2018) metrics were used to assess the quality of the genome assemblies.

For gene model prediction, *de novo* repeats were predicted using RepeatModeler v1.0.11 (Smit, AFA, Hubley, R. *RepeatModeler Open-1.0*. 2008-2015 http://www.repeatmasker.org), known protein domains were removed from this set based on identity to UniProt (Boutet et al. 2007) with the ProtExcluder.pl script from the ProtExcluder v1.2 package (Campbell et al. 2014), and the output was then used with RepeatMasker v4-0-8 (Smit, AFA, Hubley, R & Green, P. *RepeatMasker Open-4.0*. 2013-2015 http://www.repeatmasker.org) in conjunction with the Repbase library. For gene prediction, RNA-seq reads were mapped to the genome with hisat2 v2.1.0 (Kim et al. 2015). Portcullis v1.1.2 (Mapleson et al. 2018) and Mikado v1.2.2 (Venturini et al. 2018) were used to filter the resulting bam files and make first-pass gene predictions. PacBio IsoSeq data were corrected using the Iso-Seq classify + cluster pipeline (Gordon et al. 2015). Augustus v3.2 (Stanke et al. 2008) and Snap v2.37.4ubuntu0.1 (Korf 2004) were trained and then implemented through the Maker pipeline v2.31.10 (Cantarel et al. 2008) with Iso-Seq, proteins from Swiss-Prot, and processed RNA-seq added as evidence. Functional annotation was performed with BLAST v2.7.1+ (Altschul et al. 1990) and InterProScan v.5.36-75.0 (Jones et al. 2014).

### Repeat analysis

The genome of *E. cheiranthoides* was analyzed for LTR retrotransposons using LTRharvest (Ellinghaus et al. 2008), included in GenomeTools v1.5.10, with the parameters “-seqids yes - minlenltr 100 -maxlenltr 5000 -mindistltr 1000 -motif TGCA -motifmis 1 -maxdistltr 15000 -similar 85 -mintsd 4 -maxtsd 6 -vic 10 -seed 20 -overlaps best”. The genome was also analyzed using LTR_FINDER v1.07 (Xu and Wang 2007) with parameters “-D 15000 -d 1000 -L 5000 -l 100 -p 20 - C -M 0.85 -w 0”. The results from LTRharvest and LTR_FINDER were then passed as inputs to LTR_retriever v2.0 (Ou and Jiang 2018) using default parameters, including a neutral mutation rate set at 1.3×10^-8^.

Using the LTR_retriever repeat library, the genome was masked with RepeatMasker v4.0.7, and additional repetitive elements were identified *de novo* in the genome using RepeatModeler. These repeats were used with blastx v2.7.1+ (Altschul et al. 1990) against the Uniprot and Dfam libraries and protein-coding sequences were excluded using the ProtExcluder.pl script from the ProtExcluder v1.2 package (Campbell et al. 2014). The masked genome was then re-masked with RepeatMasker, with the repeat library obtained from RepeatModeler. Coverage percentages for repeat types were obtained using the fam_coverage.pl and fam_summary.pl scripts, which are included with LTR_retriever. All percentages were calculated based on the total length of the assembly.

### Genome-wide plot of genic sequence and repeats

A circular representation of the *E. cheiranthoides* genome was made with Circos v0.69-6 (Krzywinski et al. 2009). Gene and repeat densities were calculated by generating 1Mb windows and by calculating percent coverage for the features using bedtools coverage v2.26.0 (Quinlan and Hall 2010). The coverage values from the repeat library and the genome annotation were calculated independently of each other. Similarly, the total percentage of genic sequence for the genome was also calculated using bedtools genomecov v2.26.0. The gene and repeat percentages for the 1Mb windows were then plotted as histogram tracks in Circos.

For analysis of synteny between *E. cheiranthoides* and *Arabidopsis thaliana* (Arabidopsis; TAIR10; www.arabidopsis.org), the genome sequences were aligned using NUCmer, from MUMmer v3.23 (Kurtz et al. 2004) with the parameters --maxgap=500 --mincluster=100. The alignments were filtered with delta-filter -r -q -i 90 -l 1000 and coordinates of aligned segments were extracted with show-coords. The extracted coordinates were then used as the links input for Circos.

### Glucosinolate and myrosinase gene annotation in E. cheiranthoides

Known glucosinolate biosynthetic genes were annotated in *E. cheiranthoides* based on homology to Arabidopsis pathway genes in AraCyc (Rhee et al. 2006). Coding sequences for the Arabidopsis glucosinolate and myrosinase biosynthetic genes were obtained from NCBI and used in a BLASTn query against the *E. cheiranthoides* coding sequence dataset. A threshold percent identity of 80% and 70% or higher was set for glucosinolate and myrosinase genes, respectively. Protein sequences of myrosinase gene homologs in Arabidopsis and *E. cheiranthoides* were aligned using MUSCLE with UPGMA clustering and a phylogenetic tree was generated in MEGA X v10.0.4(Kumar et al. 2018) using the neighbor-joining method by sampling 1000 bootstrap replicates.

### Transcriptome sequencing of Erysimum species

To generate a high number of gene sequences required for a well-resolved phylogeny, we sequenced the foliar transcriptomes of 48 *Erysimum* species or accessions, including a first-generation inbred *E. cheiranthoides* var. *Elbtalaue*. Transcriptomes were generated from pooled leaf material of several individuals collected in the same experiment; five species were sequenced from plants in experiment 2016, 18 species from experiment 2017-1, and 25 species from experiment 2017-2 (Table S1). Leaf material was harvested 5-7 weeks after plants were transplanted into soil. To average environmental and individual effects on RNA expression, we pooled leaf material from 2-5 individual plants from one or two time points (separated by 1-2 weeks) to create a single pooled RNA sample per species (see Table S1 for details). For large-leaved species, we collected approximately 50 mg of fresh plant material from each harvested plant using a heat-sterilized hole punch (0.5 cm diameter). For smaller-leaved species, we collected an equivalent amount of material by harvesting multiple whole leaves. All leaf tissue was immediately snap frozen in liquid nitrogen and stored at −80 °C until further processing. For sample pooling, we combined leaf material of individual plants belonging to the same species in a mortar under liquid nitrogen and ground all material to a fine powder. We then weighed out 50-100 mg of frozen pooled powder for each species.

We extracted RNA from pooled leaf material using the RNeasy Plant Mini Kit (Qiagen AG, Hombrechtikon, Switzerland), including a step for on-column DNase digestion, and following the manufacturer’s instructions. The purified total RNA was dissolved in 50 µL RNase-free water, split into three aliquots, and stored at −80 °C until further processing. Assessment of RNA quality, library preparation, and sequencing were all performed by the Next Generation Sequencing Platform of the University of Bern (Bern, Switzerland). RNA quality was assessed in one aliquot per extract using a Fragment Analyzer (Model CE12, Agilent Technologies, Santa Clara, USA), and samples with low RIN scores (<7) were re-extracted and assessed again for quality. RNA libraries for TruSeq Stranded mRNA (Illumina, San Diego, USA) were assembled for each species and multiplexed in groups of eight, using unique index combinations (Illumina 2017). Groups of eight multiplexed libraries were run individually on single lanes (for a total of six lanes) of an Illumina HiSeq 3000 sequencer using 150 bp paired-end reads.

### De novo assembly of transcriptomes

RNA-seq data were cleaned with fastq-mcf v1.04.636 (https://github.com/ExpressionAnalysis/ea-utils/blob/wiki/FastqMcf.md) using the following parameters: quality = 20, minimum read length = 50. Filtered reads were assembled using Trinity v2.4.0 (Haas et al. 2013). The longest ORF was determined using TransDecoder v5.5.0 (https://github.com/TransDecoder). BUSCO v2 (Waterhouse et al. 2018) was run with lineage Embryophyta to assess gene representation and Orthofinder v2.3.1 (Emms and Kelly 2015) was used to cluster proteins from all 48 transcriptomes into orthogroups.

### Phylogenetic tree construction

We constructed phylogenetic trees using two alternative methods. In the first approach, we translated the assembled transcriptomes using TransDecoder v5.5.0. We then followed the Genome-Guided Phylo-Transcriptomics Pipeline (Washburn et al. 2017) to infer orthologous genes using synteny between genomes of *E. cheiranthoides* and Arabidopsis (TAIR10). Briefly, we obtained 26,830 orthologs between *E. cheiranthoides* and Arabidopsis through the syntenic blocks that were identified by SynMap from CoGe (https://genomevolution.org/coge/SynMap.pl). Sequences for each of the 48 *Erysimum* species and Arabidopsis (TAIR10_pep_20101214; www.arabidopsis.org) (total of 49 samples) were annotated using protein sequences of the orthologs using blastp v2.7.1 with an e-value < 10^-4^ and identity > 85%. After annotation, single copy genes and one copy of repetitive genes were kept if they were present in more than 39 (>80% of 49) species. In total, we recovered 11,890 genes, 9,868 of which had orthologs with Arabidopsis. Each of these 9,868 genes was aligned using MAFFT v7.394 (Katoh et al. 2002), and cleaned using Phyutility v2.2.6 (Smith and Dunn 2008) with the parameter -*clean 0.3*. Maximum-likelihood tree estimation for each gene was constructed using RAxML v8.2.8 (Stamatakis 2014) using the PROTCATWAG model with 100 bootstrap replicates. Finally, coalescent species tree inference was performed with these 9,868 gene trees as input, using ASTRAL-III v5.6.3 (Zhang et al. 2018).

In the second approach, we used transcriptome sequences translated by TransDecoder to predict protein sequences, after which the longest predicted protein for each gene was retained. For *E. cheiranthoides*, the genome rather than the transcriptome sequence was used. Next, we constructed gene families by running OrthoFinder v1.1.10 (Emms and Kelly 2015) on a subset of 18 *Erysimum* species (‘E18’), seven other Brassicaceae species with published genomes (*A. thaliana*, *A. lyrata*, *Boechera stricta*, *Capsella rubella*, *Eutrema salsugineum*, *Brassica rapa*, and *Schrenkiella parvula*), and three outgroup species (*Tarenaya hassleriana*, *Carica papaya*, and *Theobroma cacao*). The *Tarenaya* and *Schrenkiella* genomes were obtained from Plaza v4 Dicots (Van Bel et al. 2017), and the remaining genomes from Phytozome v12.1 (Goodstein et al. 2011). We constructed gene trees for each family using MAFFT v7.407 and FastTree v2.1.8 (Price et al. 2010) rooted with the three outgroups, and retained 3,525 subtrees with single gene copies present in at least 17 of the E18 species and in at least 6 of the 7 other Brassicaceae species. The *Erysimum* protein sequences in the 3,525 subtrees were used to identify high quality matches against the full protein sequences of the remaining 30 *Erysimum* species in the second set (‘E30’) by BLAST. High quality matches were defined as matching at least 15 of the E18 species sequences in the subtree. We retained subtrees having high quality matches in at least 24 of the E30 species, resulting in 3,098 subtrees. Finally, we identified mutual best matches between both sets of *Erysimum* species by matching the E30 protein sequences in the 3,098 subtrees against the full set of E18 protein sequences. For each subtree, we required the matches in the second set to be mutual best matches to all of the E18 proteins in the subtree, and that there be at least 24 of the E30 species in the second set. This resulted in a final set of 2,306 subtrees, from which we constructed protein sequence alignments for all *Erysimum* species and Arabidopsis using GUIDANCE2 v2.0.2 (Sela et al. 2015) and MAFFT. We then eliminated all alignment columns identified by GUIDANCE2 as low quality (column score < 0.93) and transformed protein sequences to codons. From the codon alignments, we constructed trees using RAxML v8.2.8 with the GTRGAMMA model and treating the 1^st^, 2^nd^, and 3^rd^ codon positions as three separate partitions. We concatenated all 2,306 gene family alignments, inserting gaps where a species was missing from an alignment. Additionally, we performed coalescent species tree inference with the 2,306 gene families as input, after deleting sequences with fewer than 100 non-ambiguous characters in a gene family alignment.

### Metabolite profiling of Erysimum leaves

We harvested leaf material for targeted metabolomic analysis of defense compounds from the same plants as used for transcriptome sequencing, one week after leaves for RNA extraction had been harvested. In each of the three experiments, we collected several leaves from 1-5 plants per species, and immediately snap froze the harvested leaves in liquid nitrogen. While most plant samples were screened for constitutive levels of chemical defenses only, we quantified inducibility of chemical defenses in a subset of 30 species with sufficient replication (eight or more plants) in the third experiment (2017-2). For these species, half of all plants were randomly assigned to the induction treatment and given a foliar spray of JA one week prior to harvest. Plants were sprayed with 2-3 mL of a 0.5 mM JA solution (Cayman Chemical, MI, USA) in 2% ethanol until all leaves were evenly covered in droplets on both sides. Control plants were sprayed with an equivalent amount of 2% ethanol solution. Harvested frozen plant material was lyophilized to dryness and ground to a fine powder. We weighed out 10 mg leaf powder per sample into a separate tube and added 1 mL of 70% MeOH extraction solvent. Samples were extracted by adding three 3 mm ceramic beads to each tube and shaking tubes on a Retsch MM400 ball mill three times for 3 min at 30 Hz. We centrifuged samples at 18,000 x g and transferred 0.9 mL of the supernatant to a new tube. Samples were centrifuged again, and 0.8 mL of the final supernatant was transferred to an HPLC vial for analysis by high-resolution mass spectrometry.

We analyzed extracts of individual plants (experiments 2016, 2017-1) or of multiple pooled individuals per species and induction treatment (experiment 2017-2) on an Acquity UHPLC system coupled to a Xevo G2-XS QTOF mass spectrometer with electrospray ionization (Waters, Milford MA, USA). Due to large differences in the physiochemical properties between glucosinolates and cardenolides, each plant extract was analyzed in two different modes to optimize detection of each compound class. For glucosinolates, extracts were separated on a Waters Acquity charged surface hybrid (CSH) C18 100 × 2.1 mm column with 1.7 µm pore size, fitted with a CSH guard column. The column was maintained at 40 °C and injections of 1 µl were eluted at a constant flow rate of 0.4 mL/min with a gradient of 0.1% formic acid in water (A) and 0.1% formic acid in acetonitrile (B) as follows: 0-6 min from 2% to 45 % B, 6-6.5 min from 45% to 100% B, followed by a 2 min wash phase at 100% B, and 2 min reconditioning at 2% B. For cardenolides, extracts were separated on a Waters Cortecs C18 150 × 2.1 mm column with 2.7 µm pore size, fitted with a Cortecs C18 guard column. The column was maintained at 40 °C and injections were eluted at a constant flow rate of 0.4 mL/min with a gradient of 0.1% formic acid in water (A) and 0.1% formic acid in acetonitrile (B) as follows: 0-10 min from 5% to 40 % B, 10-15 min from 40% to 100% B, followed by a 2.5 min wash phase at 100% B, and 2.5 min reconditioning at 5% B.

Compounds were ionized in negative mode for glucosinolate analysis and in positive mode for cardenolide analysis. In both modes, ion data were acquired over an m/z range of 50 to 1200 Da in MS^E^ mode using alternating scans of 0.15 s at low collision energy of 6 eV and 0.15 s at high collision energy ramped from 10 to 40 eV. For both positive and negative modes, the electrospray capillary voltage was set to 2 kV and the cone voltage was set to 20 V. The source temperature was maintained at 140 °C and the desolvation gas temperature at 400 °C. The desolvation gas flow was set to 1000 L/h, and argon was used as a collision gas. The mobile phase was diverted to waste during the wash and reconditioning phase at the end of each gradient. Accurate mass measurements were obtained by infusing a solution of leucine-enkephalin at 200 ng/mL at a flow rate of 10 µL/min through the LockSpray probe.

### Identification and quantification of defense compounds

Glucosinolates consist of a β-D-glucopyranose residue linked via a sulfur atom to a (Z)-N-hydroximinosulfate ester and a variable R group (Halkier and Gershenzon 2006). We identified candidate glucosinolate compounds from negative scan data by the exact molecular mass of glucosinolates known to occur in *Erysimum* and related species (Huang et al. 1993, Fahey et al. 2001). In addition, we screened all negative scan data for characteristic glucosinolate mass fragments to identify additional candidate compounds (Cataldi et al. 2010). For mass features with multiple possible identifications, we inferred the most likely compound identity from relative HPLC retention times and the presence of biosynthetically related compounds in the same sample. We confirmed our identifications using commercial standards for glucoiberin (3-methylsulfinylpropyl glucosinolate, Phytolab GmbH, Germany), glucocheirolin (3-methylsulfonylpropyl glucosinolate, Phytolab GmbH), and sinigrin (2-propenyl glucosinolate, Sigma-Aldrich), as well as by comparison to extracts of Arabidopsis accessions with known glucosinolate profiles. Compound abundances of all glucosinolates were quantified by integrating ion intensities of the [M-H]^-^ adducts using QuanLynx in the MassLynx software (v4.1, Waters).

All cardenolides share a highly conserved structure consisting of a steroid core (5β,14β-androstane-3β14-diol) linked to a five-membered lactone ring, which as a unit (the genin) mediates the specific binding of cardenolides to Na^+^/K^+^-ATPase (Dzimiri et al. 1987). While cardenolide genins are sufficient to inhibit Na^+^/K^+^-ATPase function, genins are commonly glycosylated or modified by hydroxylation on the steroid moiety to change the physiochemical properties and binding affinity of compounds (Dzimiri et al. 1987, Petschenka et al. 2018). We obtained commercial standards for the abundant *Erysimum* cardenolides erysimoside and helveticoside (Sigma-Aldrich), allowing us to identify these compounds through comparison of retention times and mass fragmentation patterns. Additional cardenolide compounds were tentatively identified from characteristic LC-MS fragmentation patterns. Sachdev-Gupta et al. (1990, 1993) reported fragmentation patterns for glycosides of strophanthidin, digitoxigenin, and cannogenol from *E. cheiranthoides*. Their results highlight the propensity of cardenolides to fragment at glycosidic bonds, with genin masses in particular being a prominent feature of cardenolide mass spectra. Additionally, cardenolide genins exhibit further fragmentation related to the loss of OH-groups from the steroidal core structure. We confirmed these rules of fragmentation for our mass spectrometry system using commercial standards of strophanthidin and digitoxigenin (Sigma-Aldrich). Importantly, while fragments were most abundant under high-energy conditions (MS^E^), they were still apparent under standard MS conditions, likely due to in-source fragmentation.

Characteristic fragmentation allowed us to identify candidate cardenolide compounds in a genin-guided approach, where the presence of characteristic genin fragments in a chromatographic peak indicated the likely presence of a cardenolide molecule. We then identified the parental mass of these chromatographic peaks from the presence of paired mass features separated by 21.98 m/z, corresponding to the [M+H]^+^ and [M+Na]^+^ adducts of the intact molecule. For di-glycosidic cardenolides, additional fragments corresponding to the loss of the outer sugar moiety allowed us to determine the mass and order of sugar moieties in the linear glycoside chain of the molecule. We screened our data for the presence of glycosides of strophanthidin, digitoxigenin, and cannogenol, and additional genins known to occur in *Erysimum* species (Makarevich et al. 1994). Multiple cardenolide genins can share the same molecular structure and may not be distinguished by mass spectrometry alone. Thus, all genin identifications are tentative and based on previous literature reports. We screened LC-MS data from all three experiments to generate a list of cardenolide compounds. Compounds had to be consistently detectable in at least one *Erysimum* species in at least two out of three experiments to be included in the final list. Relative compound abundances were quantified by integrating the ion intensities of the [M+H]^+^ or the [M+Na]^+^ adduct, whichever was more abundant for a given compound across all samples. In the third experiment, we added hydrocortisone (Sigma-Aldrich) to each sample as an internal standard, but between-sample variation (technical noise) was negligible compared to between-species variation.

For glucosinolate and cardenolide data separately, raw ion counts for each compound were averaged across experiments to yield robust chemotype data. Raw ion counts were standardized by the dry sample weight, possible dilution of samples, and internal standard concentrations (where available). For pooled samples, ion counts were standardized by the average dry weight calculated from all samples that contributed to a pool. The full set of standardized compound ion counts was then analyzed using linear mixed effects models (package *nlme* v3.1-137 in R v3.5.3). Because standardized ion counts still had a heavily skewed distribution, we applied a log(+0.1) transformation to all values. Log-transformed ion counts were modelled treating experiment as a fixed effect, and a species-by-compound identifier as the main random effect. Nested within the main random effect, we fitted a species-by-compound-by-experiment identifier as a second random effect to account for the difference of pooled or individual samples among experiments. The fixed effect of this model thus captures the overall differences in compound ion counts between experiments, while the main random effect captures the average deviation from an overall compound mean for each compound in each species. We extracted the overall compound mean and the main random effects from these models, providing us with average ion counts for each compound in each species on the log-scale. Negative values on the log-scale were set to zero as they would correspond to values below the limit of reliable detection of the LC-MS on the normal scale.

### Inhibition of mammal Na^+^/K^+^-ATPase by leaf extracts

Although all cardenolides target the same enzyme in animal cells, structural variation among different cardenolides can significantly influence binding affinity and thus affect toxicity (Dzimiri et al. 1987, Petschenka et al. 2018). Cardenolide quantification from LC-MS mass signal intensity does not capture such differences in biological activity, and furthermore may be challenging due to compound-specific response factors and narrow ranges of signal linearity. To evaluate whether total ion counts are an appropriate and biologically relevant measure for between-species comparisons of defense levels, we therefore quantified cardenolide concentrations by a separate method (Züst et al. 2019). For the subset of plants in the 2016 experiment, we measured the biological activity of leaf extracts on the Na^+^/K^+^-ATPase from the cerebral cortex of pigs (*Sus scrofa*, Sigma-Aldrich, MO, USA) using an *in vitro* assay introduced by Klauck and Luckner (1995) and adapted by Petschenka et al. (2013). This colorimetric assay measures Na^+^/K^+^-ATPase activity from phosphate released during ATP consumption, and can be used to quantify relative enzymatic inhibition by cardenolide-containing plant extracts. Briefly, we tested the inhibitory effect of each plant extract at four concentrations to estimate the sigmoid enzyme inhibition function from which we could determine the cardenolide content of the extract relative to a standard curve for ouabain (Sigma Aldrich, MO, USA). We dried a 100 µL aliquot of each extract used for metabolomic analyses at 45 °C on a vacuum concentrator (SpeedVac, Labconco, MO, USA). Dried residues were dissolved in 200 µL 10% DMSO in water, and further diluted 1:5, 1:50, and 1:500 using 10% DMSO. To quantify potential non-specific enzymatic inhibition that could occur at high concentrations of plant extracts, we also included control extracts from *Sinapis arvensis* leaves (a non-cardenolide producing species of the Brassicaceae) in these assays.

Assays were carried out in 96-well microplate format. Reactions were started by adding 80 μL of a reaction mix containing 0.0015 units of porcine Na^+^/K^+^-ATPase to 20 μL of leaf extracts in 10% DMSO, to achieve final well concentrations (in 100 μL) of 100 mM NaCl, 20 mM KCl, 4 mM MgCl_2_, 50 mM imidazol, and 2.5 mM ATP at pH 7.4. To control for coloration of leaf extracts, we replicated each reaction on the same 96-well plate using a buffered background mix with identical composition as the reaction mix but lacking KCl, resulting in inactive Na^+^/K^+^-ATPases. Plates were incubated at 37 °C for 20 minutes, after which enzymatic reactions were stopped by addition of 100 μL sodium dodecyl sulfate (SDS, 10% plus 0.05% Antifoam A) to each well. Inorganic phosphate released from enzymatically hydrolyzed ATP was quantified photometrically at 700 nm following the method described by Taussky and Shorr (1953).

Absorbance values of reactions were corrected by their respective backgrounds, and sigmoid dose-response curves were fitted to corrected absorbances using a non-linear mixed effects model with a 4-parameter logistic function in the statistical software R (function *nlme* with *SSfpl* in package *nlme* v3.1-137).

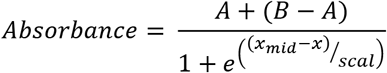

The absorbance values at four dilutions *x* are thus used to estimate the upper (*A*, fully active enzyme) and lower (*B*, fully inhibited enzyme) asymptotes, the dilution value *x_mid_* at which 50% inhibition is achieved, and a shape parameter *scal*. In order to estimate four parameters from four absorbance values per extract, the *scal* parameter was fixed for all extracts and changed iteratively to optimize overall model fit, judged by AIC. Individual plant extracts were treated as random effects to account for lack of independence within extract dilution series. For each extract we estimated *x_mid_* from the average model fit and the extract-specific random deviate. Using a calibration curve made with ouabain ranging from 10^-3^ to 10^-8^ M that was included on each 96-well plate, we then estimated the concentration of the undiluted sample in ouabain equivalents, i.e., the amount of ouabain required to achieve equivalent inhibition.

### Quantification of myrosinase activity

For the subset of plants in experiment 2017-2, we extracted the total amounts of soluble myrosinases from leaf tissue and quantified their activity as an important component of the glucosinolate defense system of these species. At time of harvest of metabolomic samples, we collected an additional set of leaf disks from each plant, corresponding to approximately 50 mg fresh weight. After determination of exact fresh weight, samples were flash frozen in liquid nitrogen and stored at −80 °C until enzyme activity measurements. Following the protocol of Travers-Martin et al. (2008), frozen leaf material was ground and extracted in Tris-EDTA buffer (200 mM Tris, 10 mM EDTA, pH 5.5) and internal glucosinolates were removed by rinsing the extracts over a DEAE Sephadex A25 column (Sigma-Aldrich). Myrosinase activities were determined by adding sinigrin to plant extracts and monitoring the enzymatic release of glucose from its activation. Control reactions with sinigrin-free buffer were used to correct for plant-derived glucose. All samples were measured in duplicate and mean values related to a glucose calibration curve, measured also in duplicate. Reactions were carried out in 96-well plates and concentrations of released glucose were measured by adding a mix of glucose oxidase, peroxidase, 4-aminoantipyrine and phenol as color reagent to each well and measuring the kinetics for 45 min at room temperature in a microplate photometer (Multiskan EX, Thermo Electron, China) at 492 nm.

### Similarity in defense profiles between Erysimum species

To quantify chemical similarity among species, we performed separate cluster analyses on the glucosinolate and cardenolide profile data averaged across the three experiments. For each species, the log-transformed average ion counts of all compounds were converted to proportions (all compounds produced by a species summing to 1). From this proportional data we then calculated pairwise Bray-Curtis dissimilarities for all species pairs using function *vegdist* in the R package *vegan* v2.5-4. We incorporated *vegdist* as a custom distance function for *pvclust* in the R package *pvclust* v2.0 (Suzuki and Shimodaira 2014), which performs multiscale bootstrap resampling for cluster analyses. We constructed dendrograms of glucosinolate and cardenolide profile similarities by fitting hierarchical clustering models (Ward’s D) and estimated support for individual species clusters from 10,000 permutations. To compare chemical similarity to phylogenetic relatedness we performed principal coordinate analyses (PCoA) on Bray-Curtis dissimilarity matrices of glucosinolate and cardenolide data using function *pcoa* in R package *ape* v5.0 (Paradis and Schliep 2019), and extracted the first two principal coordinates for each defense trait to test for phylogenetic signal.

### Relationship between plant traits and phylogenetic signal

We evaluated a prevalence of phylogenetic signal in chemical defense traits, myrosinase activity, and principal coordinates for both chemical similarity matrices using Blomberg’s *K* (Blomberg et al. 2003). *K* is close to zero for traits lacking phylogenetic signal; it approaches 1 if trait similarity among related species matches a Brownian motion model of evolution, and it can be >1 if similarity is even higher than expected under a Brownian motion model. We estimated *K* for all traits using function *phylosig* in the R package *phytools* v0.6-60 (Revell 2012). Additionally, we used the geographic coordinates for all species with known collection locations to construct pairwise geographic distances, calculated pairwise geographic dissimilarities, performed principal coordinate analyses on the geographic dissimilarity matrix, and estimated *K* for the first two components.

To test for directional effects in the evolution of compound number and abundance for both glucosinolates and cardenolides, we applied Pagel’s method (Pagel 1999). Specifically, we compared a Brownian motion model of trait evolution to a model in which additionally a directional trend is assessed by regressing the path length (i.e., molecular branch length from root to tip) against trait values. For this analysis we used the concatenated 2,306-gene tree for which branch lengths are an estimate of substitutions per site. Models were fit using function *fitContinuous* in R package *geiger* v2.0.6.2 (Harmon et al. 2008), where the default setting fits a Brownian motion model, whereas the additional argument ‘model=drift’ specifies a directional trend model. Support for directional trends in defense traits was evaluated using likelihood-ratio tests between the two models.

## Results

### E. cheiranthoides genome assembly

A total of 39.5 Gb of PacBio sequences with an average read length of 10,603 bp were assembled into 1,087 contigs with an N50 of 1.5 Mbp (Table 1). Hi-C scaffolding oriented 98.5 % of the assembly into eight large scaffolds representing pseudomolecules (Table 1, Figure S1), while 216 small contigs remained unanchored. The final assembly (v1.2) had a total length of 174.5 Mbp, representing 86% of the estimated genome size of *E. cheiranthoides* and capturing 99% of the BUSCO gene set (Table 1, Figure S2). Sequences were deposited under GenBank project ID PRJNA563696 and additionally are provided at www.erysimum.org. A total of 29,947 gene models were predicted and captured 98% of the BUSCO gene set (Figure S3). In the presumed centromere regions of each chromosome, genic sequences were less abundant, whereas repeat sequences were more common (Fig 2A). Repetitive sequences constituted approximately 29% of the genome (Table S2). Long terminal repeat retrotransposons (LTR-RT) made up the largest proportion of the repeats identified (Figure S4). Among these, repeats in the *Gypsy* superfamily constituted the largest fraction of the genome (Table S2). The majority of the LTR elements appeared to be relatively young, with most having estimated insertion times of less than 1 MYA (Figure S5). Synteny analysis showed evidence of several chromosomal fusions and fissions between the eight chromosomes of *E. cheiranthoides* and the five chromosomes of Arabidopsis (Figure 2B).

**Table 1.** Assembly metrics for the *E. cheiranthoides* genome: v0.9 = Falcon +Arrow assembly results, v1.2 = genome assembly after Hi-C scaffolding and Pilon correction.

|  | v0.9 | v1.2 pseudomolecules<br>and contigs | v1.2 pseudomolecules<br>only |
| --- | --- | --- | --- |
| total length (Mbp) | 177.4 | 177.2 | 174.5 |
| expected size (Mbp) | 205 | 205 | 205 |
| number of contigs | 1087 | 224 | 8 |
| N50 (Mbp) | 1.5 | 22.4 | 22.4 |
| complete BUSCOs (out of 1,375) | 1359 | 1346 | 1356 |
| complete and single copy BUSCOs (out of 1,375) | 1271 | 1300 | 1306 |
| complete and duplicated BUSCOs (out of 1,375) | 88 | 46 | 50 |
| fragmented BUSCOs (out of 1,375) | 5 | 8 | 6 |
| missing BUSCOs (out of 1,375) | 11 | 21 | 13 |

**Figure 2.**
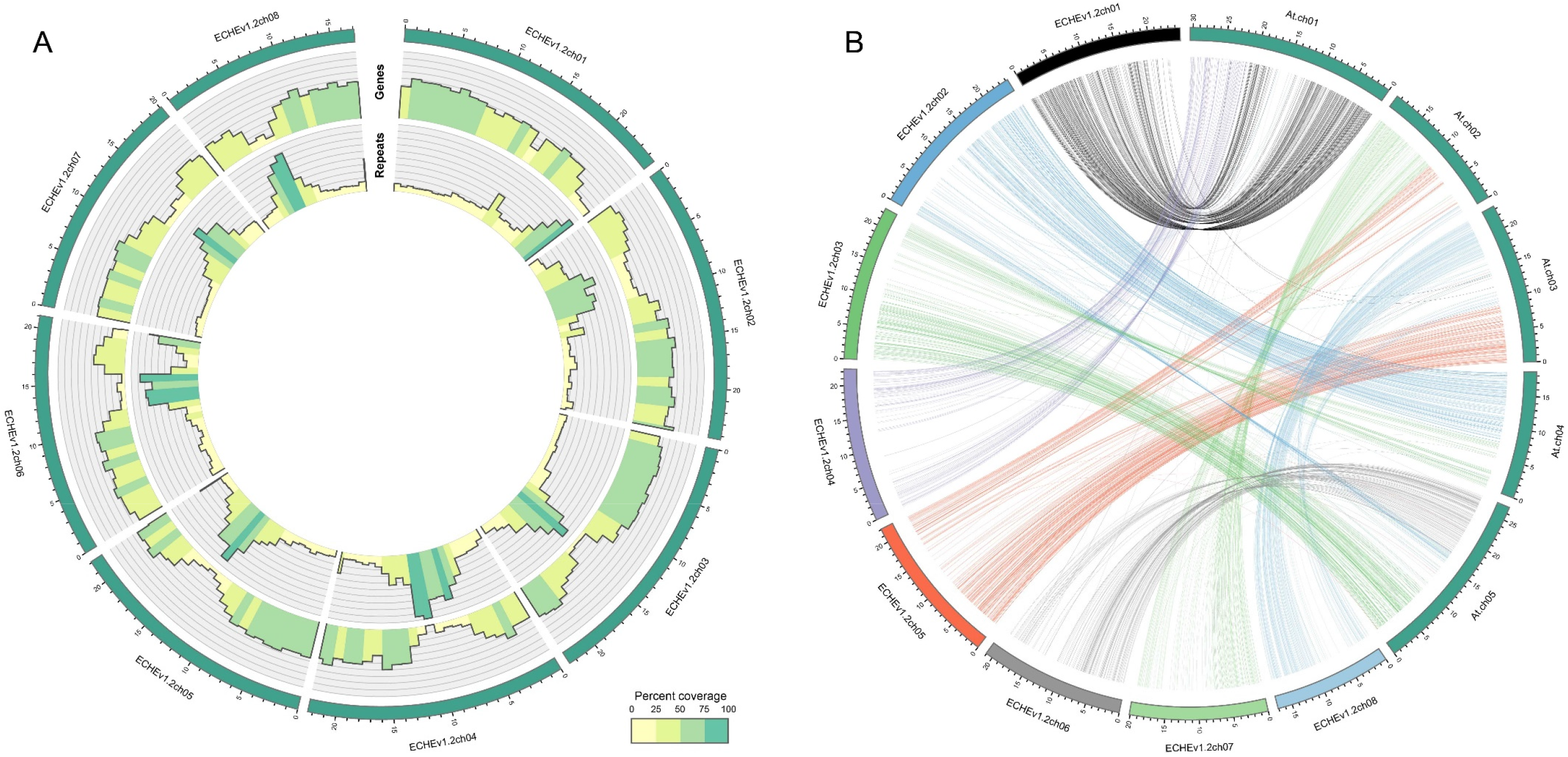
(A) Circos plot of the *E. cheiranthoides* genome with gene densities (outer circle) and repeat densities (inner circle) shown as histogram tracks. Densities are calculated as percentages for 1 Mb windows. (B) Synteny plot of *E. cheiranthoides* and *A. thaliana*. Lines between chromosomes connect aligned sequences between the two genomes.

### Glucosinolate and myrosinase genes in the E. cheiranthoides genome

Three aliphatic glucosinolates – glucoiberverin (3-methylthiopropyl glucosinolate), glucoiberin, and glucocheirolin – have been reported as the main glucosinolates in *E. cheiranthoides* (Cole 1976, Huang et al. 1993). We confirmed their dominance in *E. cheiranthoides* var. *Elbtalaue*, but also identified additional aliphatic and indole glucosinolates at lower concentrations. By making use of the glucosinolate biosynthetic pathway for Arabidopsis (Halkier and Gershenzon 2006) and comparing nucleotide coding sequences of Arabidopsis and *E. cheiranthoides*, we identified homologs of genes encoding both indole (Figure 3) and aliphatic (Figure 4) glucosinolate biosynthetic enzymes. Homologs of all genes of the complete biosynthetic pathway for glucobrassicin (indol-3-ylmethyl glucosinolate) and its 4-hydroxy and 4-methoxy derivatives were present in *E. cheiranthoides* (Figure 3). Consistent with the absence of neoglucobrassicin (1-methoxy-indol-3-ylmethyl glucosinolate) in *E. cheiranthoides* var. *Elbtalaue*, we did not find homologs of the Arabidopsis genes encoding the biosynthesis of this compound.

**Figure 3.**
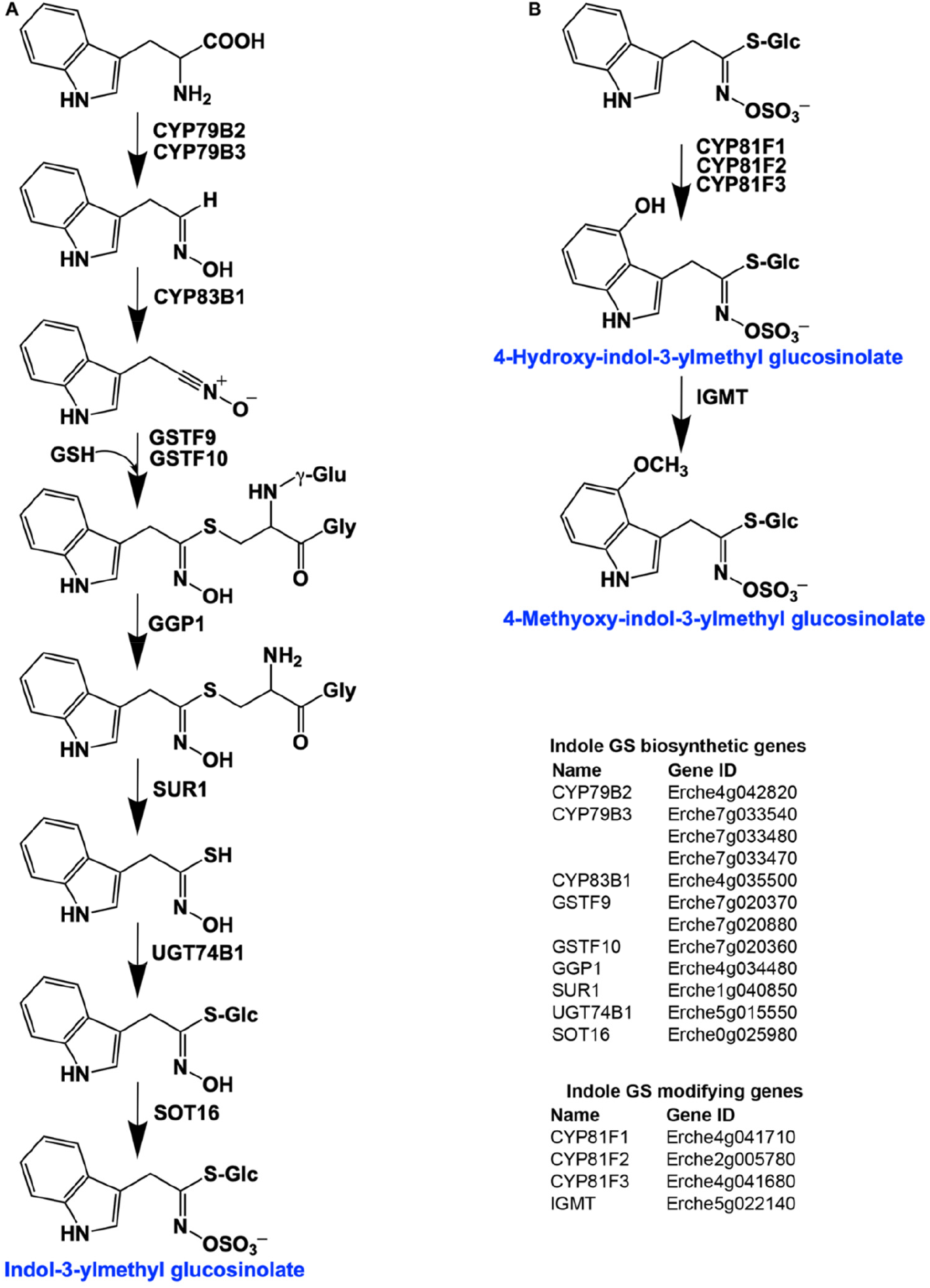
Identification of indole glucosinolate biosynthetic genes and glucosinolate-modifying genes in *Erysimum cheiranthoides*. (A) Starting with tryptophan, indole glucosinolates are synthesized using some enzymes that also function in aliphatic glucosinolates biosynthesis (GGP1; SUR1; UGT74B1) while also using indole glucosinolate-specific enzymes. (B) Indole glucosinolates can be modified by hydroxylation and subsequent methylation. Glucosinolates with names highlighted in blue were identified in *Erysimum cheiranthoides* var. *Elbtalaue*. Abbreviations: cytochrome P450 monooxygenase (CYP); glutathione S-transferase F (GSTF); glutathione (GSH); γ-glutamyl peptidase 1 (GGP1); SUPERROOT 1 C-S lyase (SUR1); UDP-dependent glycosyltransferase (UGT); sulfotransferase (SOT); glucosinolate (GS); indole glucosinolates methyltransferase (IGMT).

**Figure 4.**
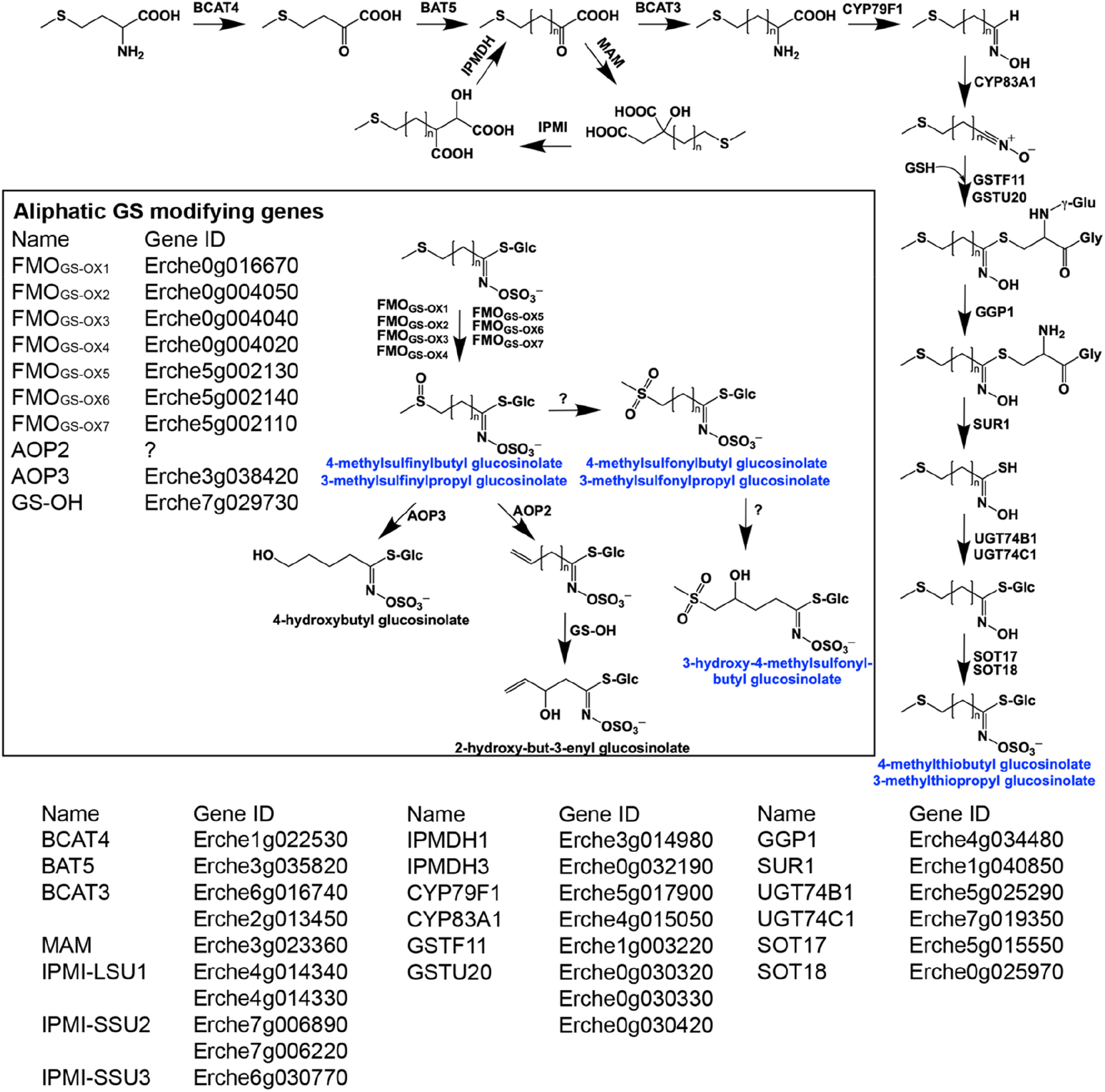
Identification of aliphatic glucosinolate biosynthetic genes in *Erysimum cheiranthoides* starting from methionine and modifications of aliphatic glucosinolates (black box). Glucosinolates with names highlighted in blue were identified in *Erysimum cheiranthoides* var. *Elbtalaue.* Abbreviations: branched-chain aminotransferase (BCAT); bile acid transporter (BAT); methylthioalkylmalate synthase (MAM); isopropylmalate isomerase (IPMI); large subunit (LSU); small subunit (SSU); isopropylmalate dehydrogenase(IPMDH); cytochrome P450 monooxygenase (CYP); glutathione S-transferase F (GSTF); glutathione S-transferase Tau (GSTU); glutathione (GSH); γ-glutamyl peptidase 1 (GGP1); SUPERROOT 1 C-S lyase (SUR1); UDP-dependent glycosyltransferase (UGT); sulfotransferase (SOT); flavin monooxygenase (FMO); glucosinolate oxoglutarate-dependent dioxygenase (AOP); 3-butenyl glucosinolate 2-hydroxylase (GS-OH).

Genes encoding the complete biosynthetic pathway of the *E. cheiranthoides* aliphatic glucosinolates glucoiberverin, glucoiberin, glucoerucin (4-methylthiobutyl glucosinolate), and glucoraphanin (4-methylsulfinylbutyl glucosinolate) were present in the genome (Figure 4). Because the *E. cheiranthoides* methylsulfonyl glucosinolates glucocheirolin, 4-methylsulfonylbutyl glucosinolate, and 3-hydroxy-4-methylsulfonylbutyl glucosinolate are not present in Arabidopsis, genes encoding their biosynthesis could not be identified. Consistent with the absence of an Arabidopsis AOP2 homolog, we did not find alkenyl glucosinolates in *E. cheiranthoides*.

In response to insect feeding or pathogen infection, glucosinolates are activated by myrosinase enzymes (Halkier and Gershenzon 2006). Between-gene phylogenetic comparisons revealed that homologs of known Arabidopsis myrosinases, the main foliar myrosinases *TGG1* and *TGG2* (Barth and Jander 2006), root-expressed *TGG4* and *TGG5* (Andersson et al. 2009), and likely pseudogenes *TGG3* and *TGG6* (Rask et al. 2000, Zhang et al. 2002), were also present in the *E. cheiranthoides* genome (Figure S6). Additionally, we found homologs of the more distantly related Arabidopsis myrosinases *PEN2* (Bednarek et al. 2009, Clay et al. 2009) and *PYK10* (Sherameti et al. 2008, Nakano et al. 2017). Thus, the pathway of glucosinolate activation appears to be largely conserved between Arabidopsis and *E. cheiranthoides*.

### Phylogenetic relationship of 48 Erysimum species

Assemblies of transcriptomes from 48 *Erysimum* species (including *E. cheiranthoides*) had N50 values ranging from 574 - 2,160 bp (Table S3). Transcriptome assemblies contained completed genes from 54% - 94% of the BUSCO set and coding sequence lengths were generally shorter on average than the *E. cheiranthoides* coding sequence lengths (Table S3). Transcriptome sequences were deposited under GenBank project ID PRJNA563696 and at www.erysimum.org. The large number of orthologous gene sequences identified among the *E. cheiranthoides* genome and the 48 transcriptomes allowed us to infer phylogenetic relatedness with high confidence. We assume that the coalescent phylogeny generated by the first approach of tree construction using 9,868 syntenic gene sequences represents our best estimate of species relationships (Figure 5). However, both phylogenies generated by the second approach using 2,306 orthologous genes had highly similar tree topologies (Figures S7, S8), suggesting overall high reliability of our results regardless of the phylogenetic inference method.

**Figure. 5.**
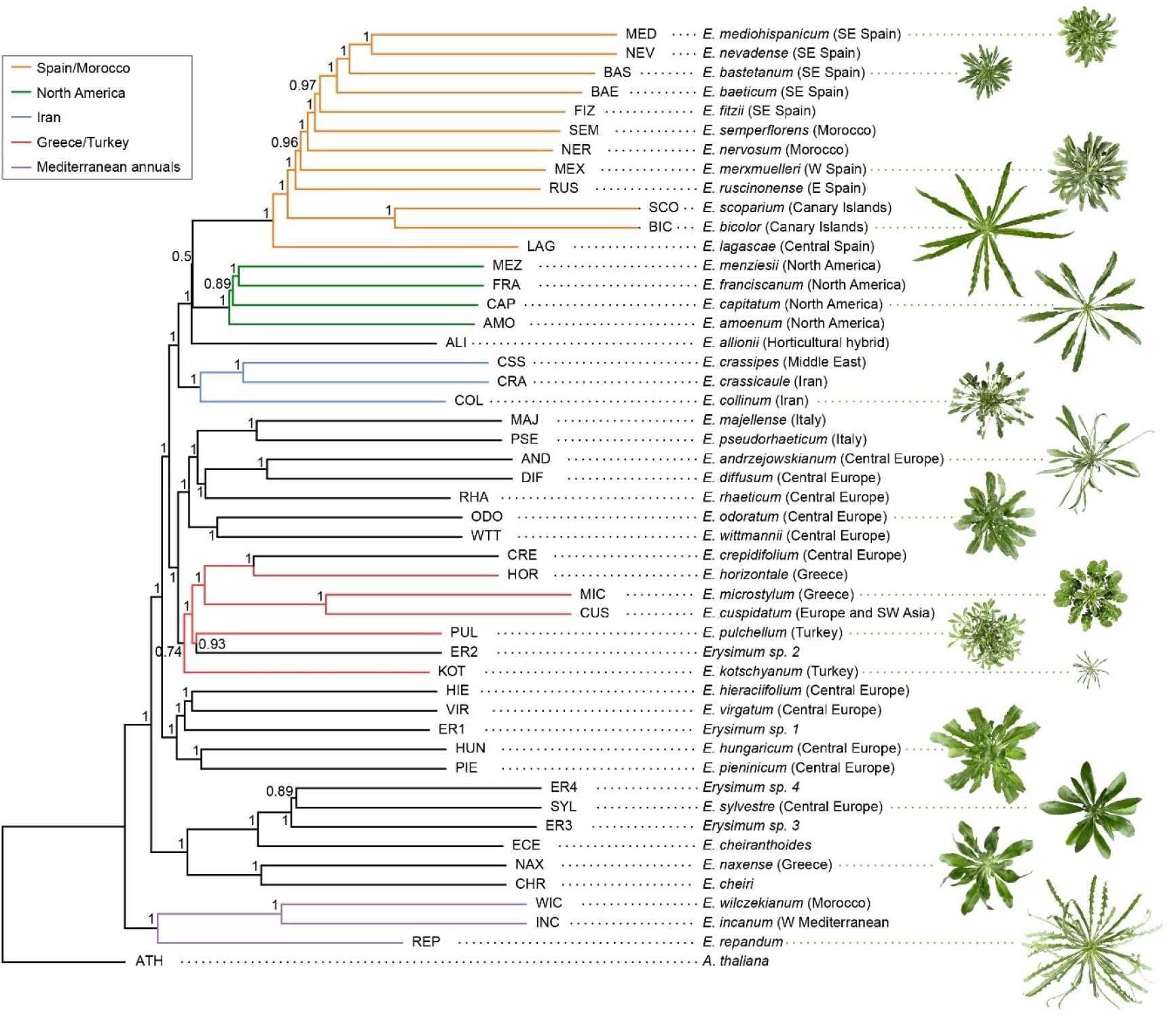
Genome-guided coalescent species tree of 48 *Erysimum* species. Phylogenetic relationships were inferred from 9,868 orthologous genes using ASTRAL-III. Nodes are labelled with local posterior probability, indicating level of support. Geographic range of species is provided in parentheses. The horticultural species *E. cheiri* and the weedy species *E. cheiranthoides* and *E. repandum* are of European origin but are now widespread across the Northern Hemisphere. Clades of species from shared geographic origins are highlighted in different colors. On the right, pictures of rosettes of a representative subset of species is provided to highlight the morphological diversity within this genus. Plants are of same age and relative size differences are conserved in the pictures.

Virtually all phylogenetic nodes of the coalescent trees had high statistical support (all but one node with local posterior probability > 0.7; Figure 5). The three Mediterranean annual species *E. incanum* (INC), *E. repandum* (REP), and *E. wilczekianum* (WIC) formed a monophyletic sister clade to all other sequenced species. The only other annual in the set of sampled species, *E. cheiranthoides* (ECE), was part of a second early-diverging clade, together with several perennial species from Greece and central Europe, including the widespread ornamental *E. cheiri* (CHR). Several other geographic clades were apparent in the phylogeny, with species from the Iberian peninsula/Morocco, North America, Iran, and Greece forming additional distinct geographic clades. The clear geographic structure of the phylogeny was confirmed by a very strong phylogenetic signal for the first two components of the geographic principal coordinate analysis (Table 2).

**Table 2.** Measure of phylogenetic signal for total defensive traits and principal coordinates of the cardenolide and glucosinolate similarity matrices (PCO) using Blomberg’s *K*. Significant values are highlighted in bold.

| Plant trait | K statistics | *P* (10'000 simulations) |
| --- | --- | --- |
| Glucosinolate PCO1 (18.8%) | 0.89 | 0.176 |
| Glucosinolate PCO2 (13.6%) | 0.85 | 0.375 |
| Total glucosinolate concentrations | 0.88 | 0.216 |
| Number of glucosinolate compounds | 0.94 | 0.060 |
| Myrosinase activity | 1.02 | **0.019** |
| Cardenolide PCO1 (16.5%) | 1.55 | **<0.001** |
| Cardenolide PCO2 (12.2%) | 1.18 | **<0.001** |
| Total cardenolide concentrations | 1.08 | **0.016** |
| Number of cardenolide compounds | 1.16 | **0.001** |
| Geographical PCO1 (15.8%)<sup>1</sup> | 2.33 | **<0.001** |
| Geographical PCO2 (12.8%)<sup>1</sup> | 1.37 | **<0.001** |

<sup>1</sup> results are for 43 species with reliable geographic information

### Glucosinolate diversity and myrosinase activity

Across the 48 *Erysimum* species, we identified 25 candidate glucosinolate compounds with distinct molecular masses and HPLC retention times (Table S4). Of these, 24 compounds could be assigned to known glucosinolate structures with high certainty. The remaining compound appeared to be an unknown isomer of glucocheirolin. Individual *Erysimum* species produced between 5 and 18 glucosinolates (Figure 6A), and total glucosinolate concentrations were highly variable among species (Figure 6B). The ploidy level of species explained a significant fraction of total variation in the number of glucosinolates produced (F_4,38_ = 4.63, p = 0.004), with hexaploid species producing the highest number of compounds (Figure S9). However, neither the numbers of distinct glucosinolates nor the total concentrations exhibited a phylogenetic signal (Table 2). Similarly, there was no evidence for a directional trend in either glucosinolate trait (Table 3), suggesting that glucosinolate defenses varied among species independently of phylogenetic history.

**Table 3.** Maximum-likelihood estimation of directional trends (β, root-to-tip regression) in cardenolide and glucosinolate evolution. Directional trends are assessed for gradual models of evolution using the concatenated 2306-gene tree in which branch lengths are proportional to estimated substitutions per site. Each directional model is assessed against a random walk model without a trend.

| Plant traits | $\beta$ | Likelihood ratio | p-value |
| --- | --- | --- | --- |
| Total glucosinolate concentrations | -37,160.0 | 0.001 | 0.974 |
| Number of glucosinolate compounds | 40.7 | 0.049 | 0.825 |
| Total cardenolide concentrations | 707,093.6 | 1.759 | 0.185 |
| Number of cardenolide compounds | -218.4 | 0.173 | 0.677 |

**Figure 6.**
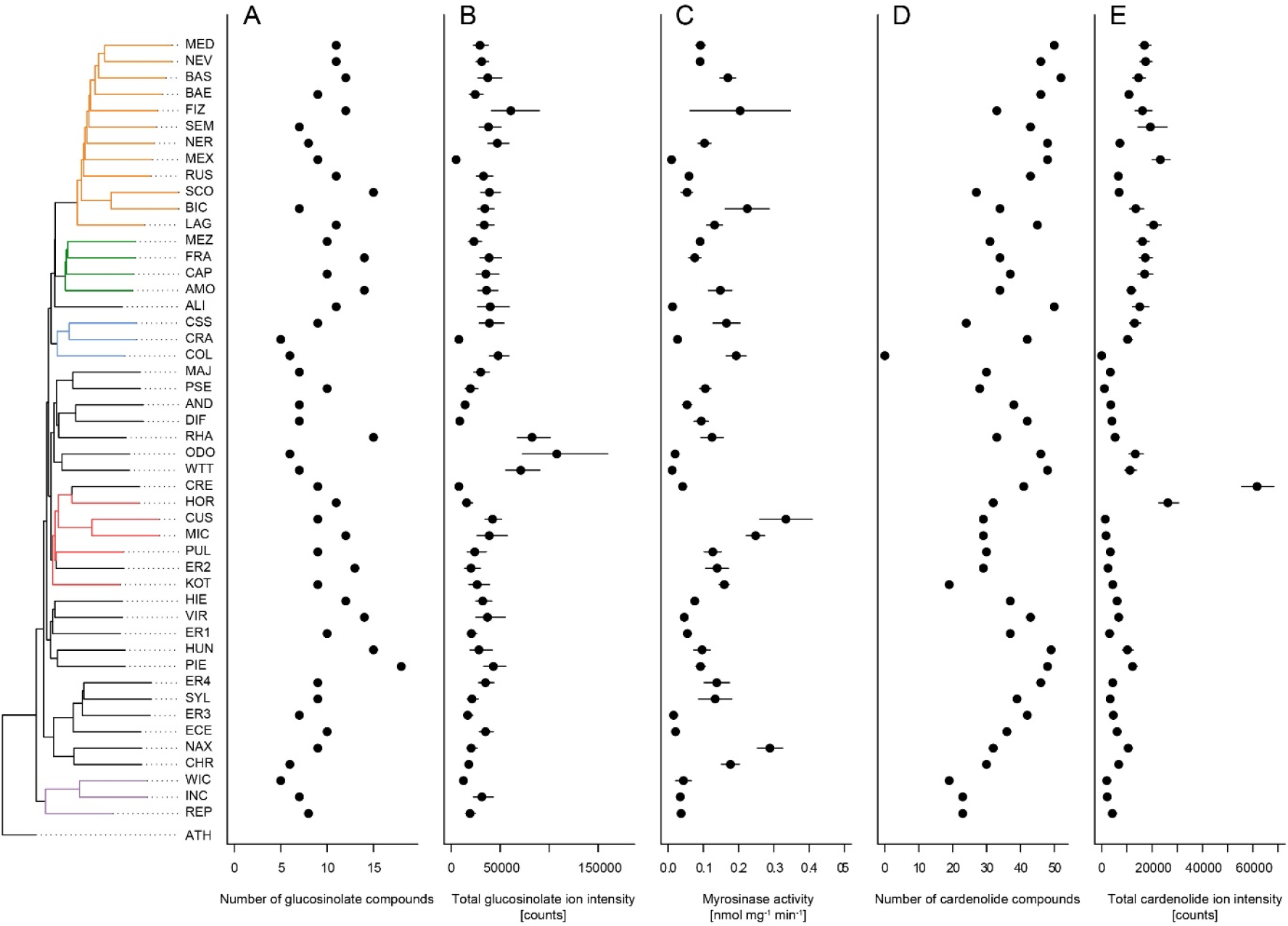
Mean defense traits of 48 *Erysimum* species, grouped by phylogenetic relatedness. Not all traits could be quantified for all species. (A) Total number of glucosinolate compounds detected in each species. (B) Total glucosinolate concentration found in each species, quantified by total ion intensity in mass spectrometry analyses. Values are means ± 1SE. (C) Quantification of glucosinolate-activating myrosinase activity. Enzyme kinetics were quantified against the standard glucosinolate sinigrin and are expressed per unit fresh plant tissue. Values are means ± 1SE. (D) Total number of cardenolide compounds detected in each species. (E) Total cardenolide concentrations found in each species, quantified by total ion intensity in mass spectrometry analyses. Values are means ± 1SE.

Clustering species according to similarity in glucosinolate profiles mostly resulted in chemotype groups corresponding to known underlying biosynthetic genes, although support for individual species clusters was variable (Figure 7). The majority of species produced glucoiberin as the primary glucosinolate. Of these, approximately half also produced sinigrin as a second dominant glucosinolate compound. Further chemotypic subdivision, related to the production of glucocheirolin and 2-hydroxypropyl glucosinolate, appeared to be present but only had relatively weak statistical support. However, eight species clearly differed from these general patterns. The species *E. allionii* (ALI), *E. rhaeticum* (RHA), and *E. scoparium* (SCO) mostly lacked glucosinolates with 3-carbon side-chains, but instead accumulated glucosinolates with 4-, 5- and 6-carbon side-chains. The two closely-related species *E. odoratum* (ODO) and *E. wittmannii* (WIT) predominantly accumulated indole glucosinolates, while *E. collinum* (COL), *E. pulchellum* (PUL), and accession ER2 predominantly produced glucoerypestrin (3-methoxycarbonylpropyl glucosinolate), a glucosinolate that is exclusively found within *Erysimum* (Fahey et al. 2001). As with total glucosinolate concentrations, similarity in glucosinolate profiles of the species was again unrelated to phylogenetic relatedness (Table 2).

**Figure 7.**
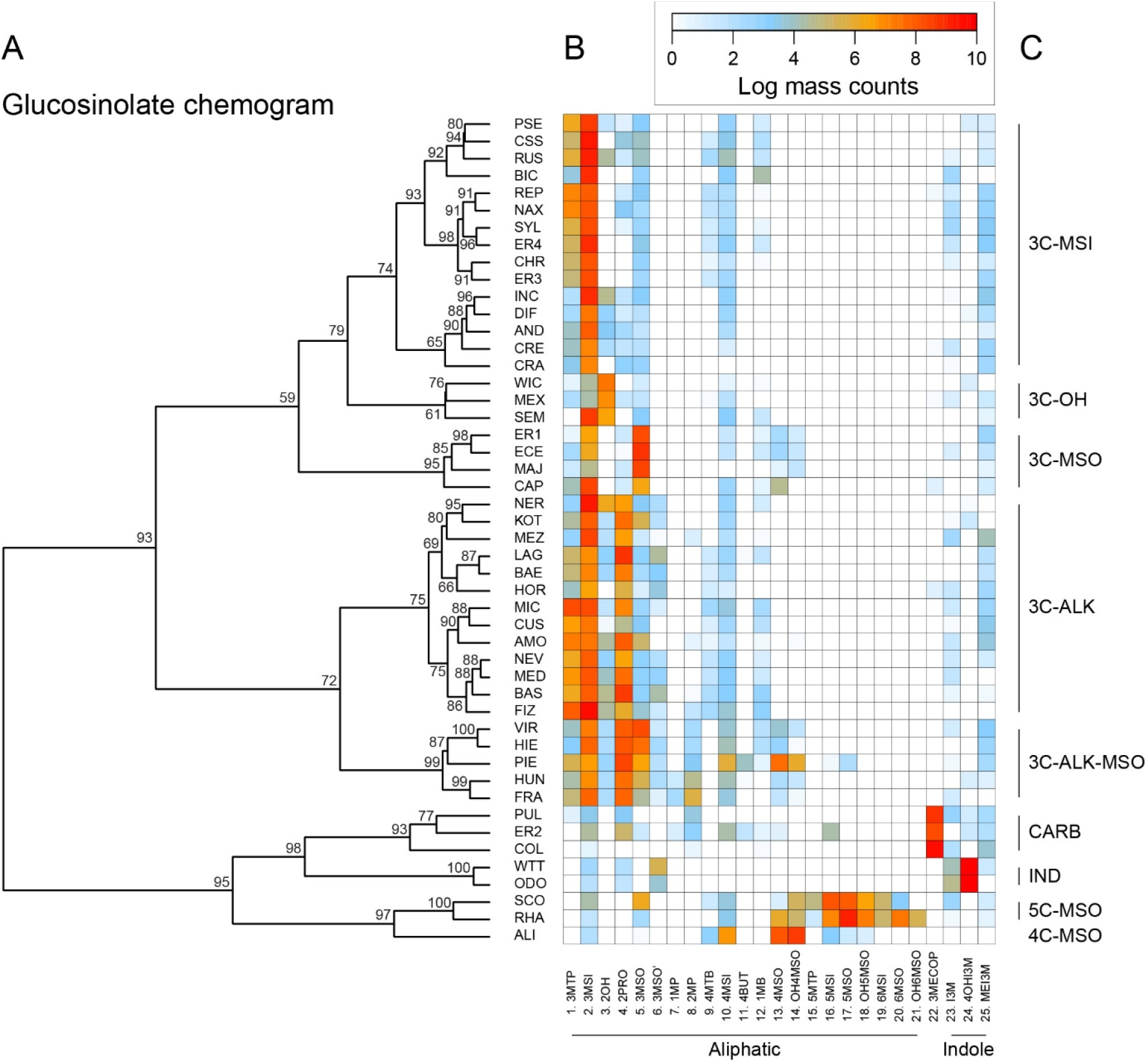
(A) Glucosinolate chemogram clustering species according to overlap in glucosinolate profiles. Values at nodes are confidence estimates (approximately unbiased p-value, function *pvclust* in R) based on 10,000 iterations of multiscale bootstrap resampling. (B) Heatmap of glucosinolate profiles expressed by the 48 *Erysimum* species. Color intensity corresponds to log-transformed integrated ion counts recorded at the exact parental mass ([M-H]^-^) for each compound, averaged across samples from multiple independent experiments. Compounds are grouped by major biosynthetic classes and labelled using systematic short names. See Table S4 for full glucosinolate names and additional compound information. (C) Classification of species chemotype based on predominant glucosinolate compounds. 3C/4C/5C = length of carbon side chain, MSI = methylsulfinyl glucosinolate, MSO = methylsulfonyl glucosinolate, OH = side chain with hydroxy group, ALK = side chain with alkenyl group, CARB = carboxylic glucosinolate, IND = indole glucosinolate.

As glucosinolates require activation by myrosinase enzymes upon tissue damage by herbivores, myrosinase activity in leaf tissue determines the rate at which toxins are released. We quantified myrosinase activity of *Erysimum* leaf extracts and found it to be highly variable among species (Figure 6C). After grouping species into nine chemotypes defined by chemical similarity and the production of characteristic glucosinolate compounds (Figure 7C), we found that myrosinase activity significantly differed among these chemotypes (Figure 8, F_8,33_ = 7.06, p < 0.001). Chemotypes that predominantly accumulated methylsulfonyl glucosinolates, hydroxy glucosinolates, or indole glucosinolates had low to negligible activity against the assayed glucosinolate sinigrin. It is important to note that sinigrin is an alkenyl glucosinolate and activity with other, structurally dissimilar glucosinolates may differ. After chemotype differences were accounted for, myrosinase activity was related positively to total glucosinolate concentrations (F_1,33_ = 5.92, p = 0.021). Surprisingly, uncorrected myrosinase activity was the only glucosinolate-related trait that showed a significant phylogenetic signal (Table 2).

**Figure 8.**
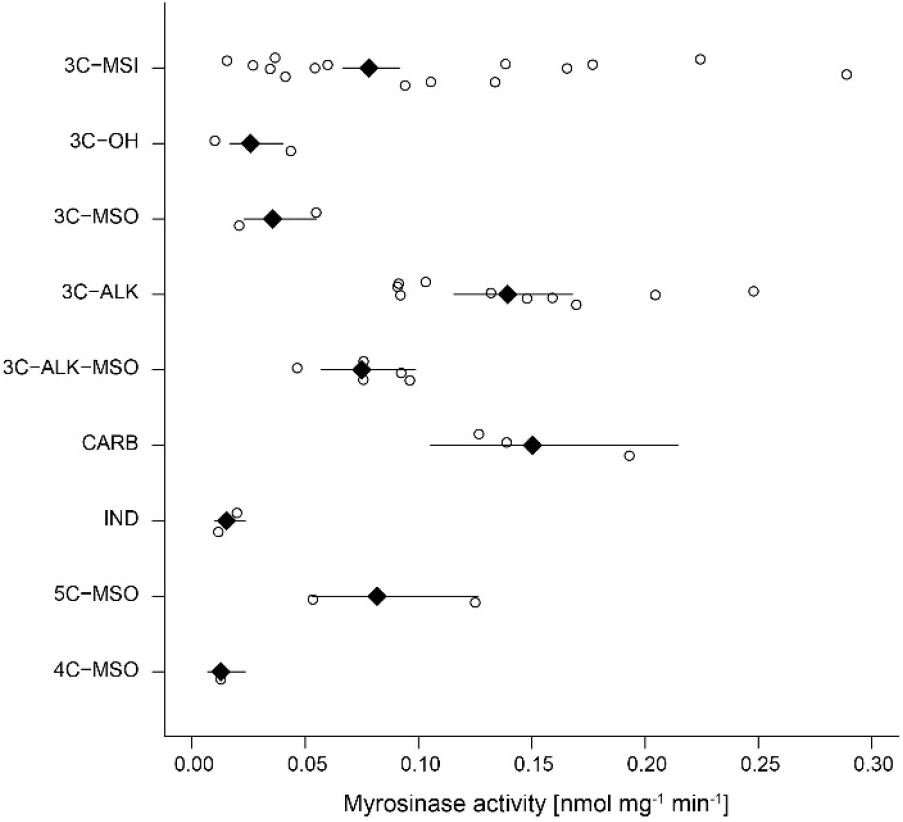
Myrosinase activity of *Erysimum* leaf extracts grouped by glucosinolate chemotype. Open circles are species means and black diamonds are chemotype means ± 1 SE. See also Figure 7 for chemotype information. 3C/4C/5C = length of carbon side chain, MSI = methylsulfinyl glucosinolate, MSO = methylsulfonyl glucosinolate, OH = side chain with hydroxy group, ALK = side chain with alkenyl group, CARB = carboxylic glucosinolate, IND = indole glucosinolate.

### Cardenolide diversity

With the exception of *E. collinum* (COL), which only contained trace amounts of cardenolides in leaves, all *Erysimum* species contained diverse mixtures of cardenolide compounds and accumulated considerable amounts of cardenolides (Figure 6D-E). The ploidy level of species again explained a significant fraction of the total variation in the number of cardenolides (F_4,38_ = 3.47, p = 0.016), with hexaploid species producing the highest average number of compounds (Figure S9). To obtain an estimate of biological activity and evaluate quantification from total MS ion counts, we used an established assay that quantifies cardenolide concentrations from specific inhibition of animal Na^+^/K^+^-ATPase by crude *Erysimum* leaf extracts. We found generally strong enzymatic inhibition, with leaves of *Erysimum* species containing an equivalent of 5.72 ± 0.12 µg mg^-1^ (± 1 SE) of the reference cardenolide ouabain on average. Despite only producing trace amounts of cardenolides, *E. collinum* (COL) extracts caused significantly stronger inhibition than the Brassicaceae control, *S. arvensis* (Figure S10). Overall, quantification of cardenolide concentrations by Na^+^/K^+^-ATPase inhibition was highly correlated with the total MS ion count (Fig S8, r = 0.95, p < 0.001). Thus, the use of ion count data for cross-species comparisons was appropriate for this purpose. Both the total numbers of compounds and the total abundances exhibited a strong phylogenetic signal (Table 2), indicating that closely-related species were more similar in their cardenolide traits than expected by chance. However, there was again no evidence for a directional trend in the evolution of either number or abundance of cardenolides (Table 3), suggesting a rapid rather than a gradual gain of cardenolide diversity, which is also evident from the considerable number of cardenolide compounds present in the earliest-diverging species (Figure 6D).

Cardenolide diversity was considerably higher than that of glucosinolates, with a total of 95 distinguishable candidate cardenolide compounds identified across the 48 *Erysimum* species (Table S5). Of these, 46 compounds had distinct molecular masses and mass fragments, while the remaining compounds likely were isomers, sharing a molecular mass with another compound but having a distinct HPLC retention time. The 95 putative cardenolides comprised nine distinct genins (Figures 9, S11), the majority of which were glycosylated with digitoxose, deoxy hexoses, xylose, or glucose moieties. Only digitoxigenin and cannogenol accumulated as free genins, while all other compounds occurred as either mono- or di-glycosides. A major source of isomeric cardenolide compounds was thus likely the incorporation of different deoxy hexoses of equivalent mass, such as rhamnose, fucose, or gulomethylose. A subset of compounds had molecular masses that were heavier by 42.011 m/z than known mono- or di-glycoside cardenolides. Such a gain in mass corresponds to the gain of an acetyl-group, and mass fragmentation patterns indicated that these compounds were acetylated on the first sugar moiety (Table S5). Out of the nine detected genins, six had previously been described from *Erysimum* species (Makarevich et al. 1994). In addition, we identified three previously undescribed mass features with fragmentation patterns characteristic of cardenolide genins (Figure S11). Of these three, one matched an acetylated digitoxigenin (also known as oleandrigenin), a common cardenolide in *Nerium oleander*. The other two matched molecular structures of digitoxigenin-formate (also known as gitaloxigenin) and strophanthidin-formate. Formate adducts can sometimes be formed during LC-MS due to the addition of formic acid to solvents, although this is less common with positive ionization. To exclude to possibility that these were technical artefacts, we analyzed a subset of extracts by LC-MS without the addition of formic acid and found both genin-formates at comparable concentrations (Figure S12). We therefore assume that all three novel structures are natural variants of cardenolides produced by *Erysimum* plants, even though we currently lack final structural elucidation.

**Figure 9.**
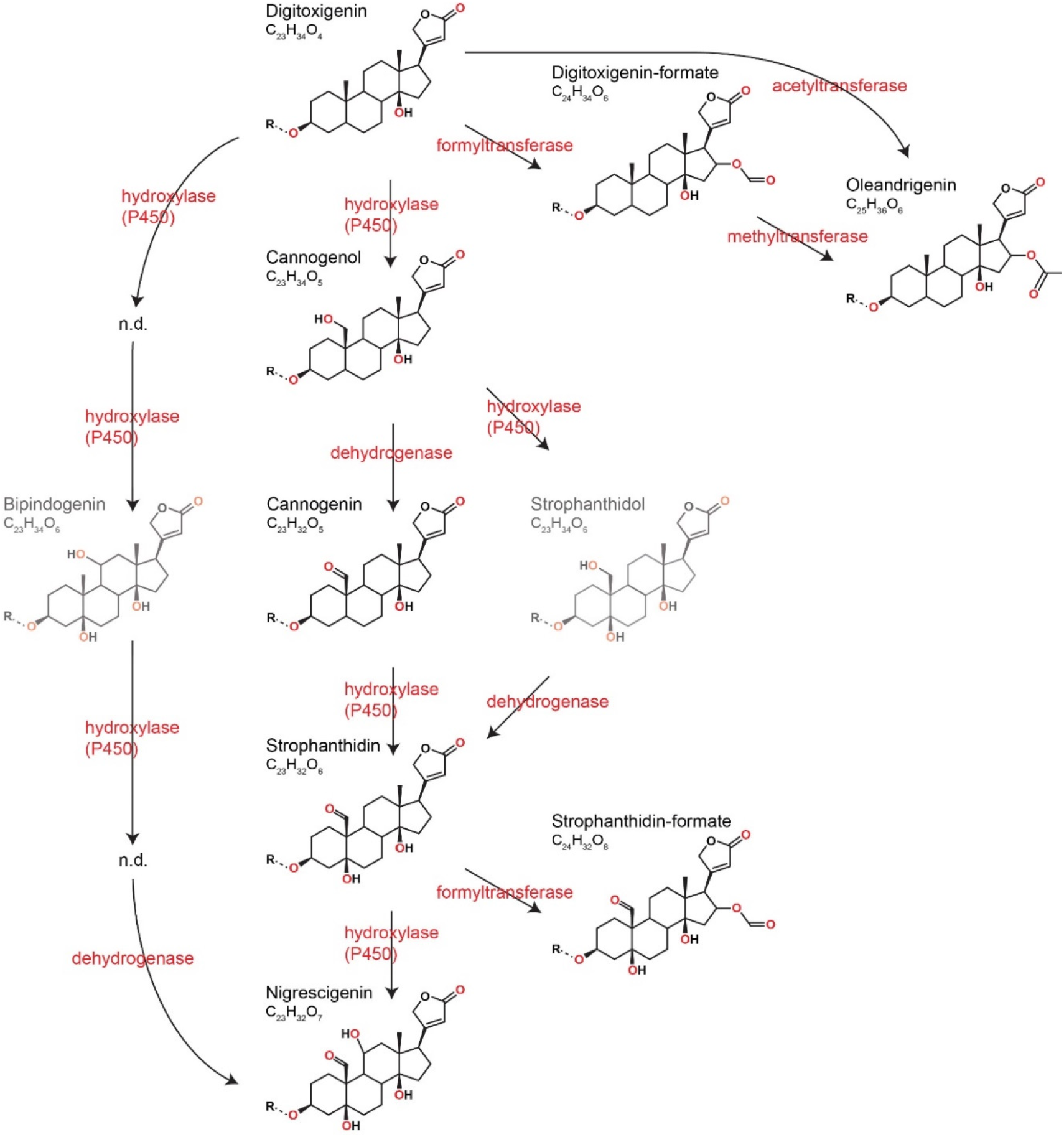
Predicted pathways of cardenolide genin modification in *Erysimum*. Genin diversity likely originates from digitoxigenin, which by hydroxylases (P450-like enzymes), dehydrogenases, and formyl-, methyl-, or acetyltransferases is transformed into structurally more complex cardenolides. Oleandrigenin could be derived from digitoxigenin or from digitoxigenin-formate, with the frequent co-occurrence of oleandrigenin and digitoxigenin-formate in leaf extracts suggesting the latter. According to their exact mass, frequently detected dihydroxy-digitoxigenin compounds (C_23_H_34_O_6_) could be either bipindogenin or strophanthidol. While bipindogenin cardenolides have commonly been reported for *Erysimum* species in the literature, their structure would require additional intermediate structures that were not detected here (n.d.). Thus, strophanthidol appears to be the more likely isomer to occur in *Erysimum*. All cardenolide genins are further modified by glycosylation at a conserved position in the molecule (R).

Clustering of species based on similarity in cardenolide profiles revealed fewer obvious species clusters than for glucosinolates, and particularly higher-level species clusters had only weak statistical support (Figure 10). A clear exception to this was a species cluster that included *E. cheiranthoides* (ECE) and *E. sylvestre* (SYL), which lacked several otherwise common cannogenol- and strophanthidin-glycosides, while accumulating unique digitoxigenin-glycosides. A second major cluster that was visually apparent – yet not statistically significant – separated groups of species that did or did not produce glycosides of the newly discovered putative strophanthidin-formate (Figure 10). Similarity in cardenolide profiles among species quantified as the first and second principal coordinate of the Bray-Curtis dissimilarity matrix exhibited a very strong phylogenetic signal (Table 2), suggesting that closely-related species not only were more similar in their total cardenolide concentrations, but also had more similar cardenolide profiles than expected by chance.

**Figure 10.**
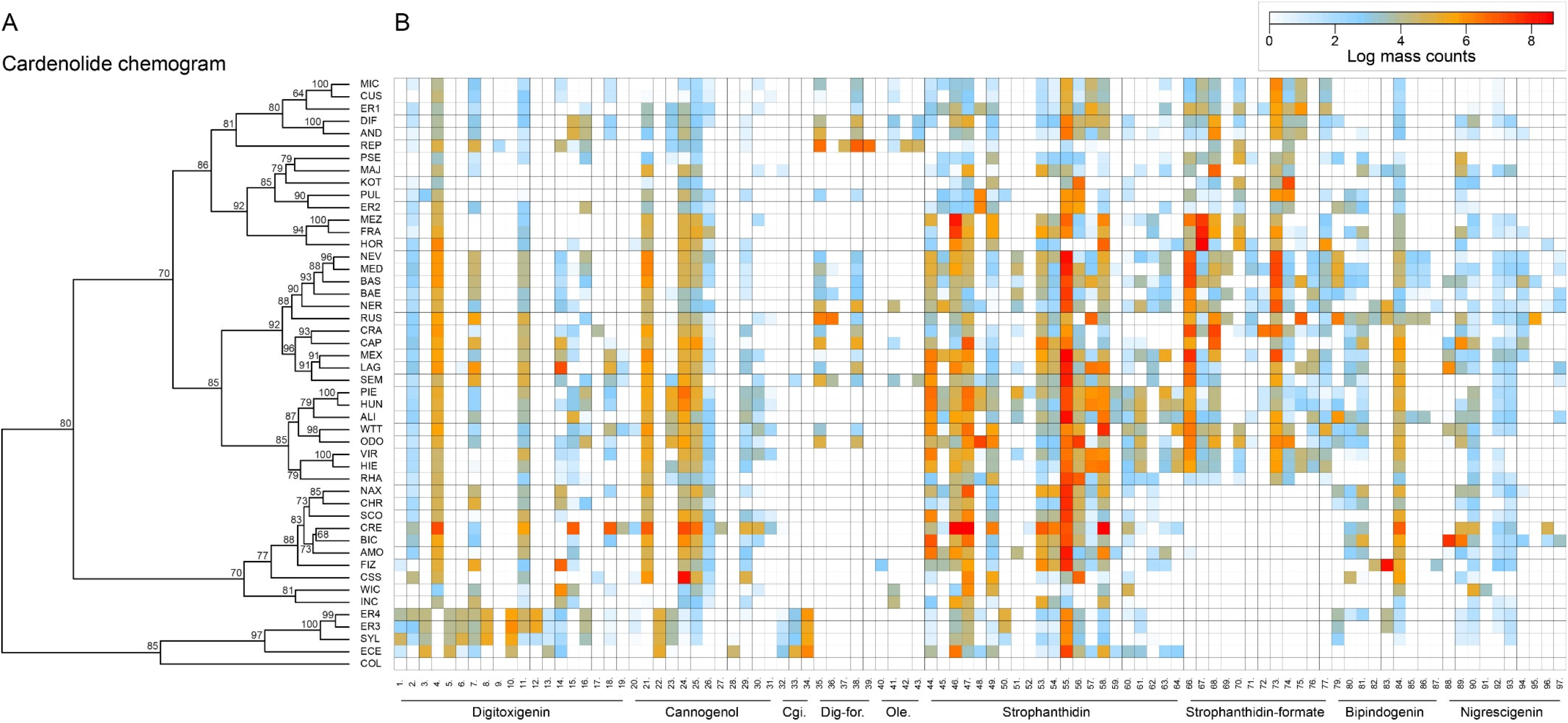
(A) Cardenolide chemogram clustering species according to similarity in cardenolide profiles. Values at nodes are confidence estimates based on 10,000 iterations of multiscale bootstrap resampling. (B) Heatmap of cardenolide profiles expressed by the 48 *Erysimum* species. Color intensity corresponds to log-transformed integrated ion counts recorded at the exact parental mass ([M+H]^+^ or [M+Na]^+^, whichever was more abundant) for each compound, averaged across samples from multiple independent experiments. The species *E. collinum* (COL) only expressed trace amounts of cardenolides, which are not visible on the color scale. Compounds are grouped by shared genin structures. Cgi. = Cannogenin, Dig-for. = Digitoxigenin-formate, Ole. = Oleandrigenin. See Table S5 for additional compound information.

### Macroevolutionary patterns in defense and inducibility

Given the very distinct patterns for glucosinolate and cardenolide diversity among *Erysimum* species, it is unsurprising that concentrations of the two defense traits were not correlated (Pearson’s correlation: r = −0.09, p = 0.534). Foliar application of JA was expected to stimulate defense levels in plant leaves, and among the 30 tested species, glucosinolate levels responded positively to JA, with the majority of species increasing their foliar glucosinolate concentration (Figure 11). However, the glucosinolate inducibility of a species was independent of constitutive glucosinolate levels (r = −0.26, p = 0.169). By contrast, the majority of species exhibited lower cardenolide levels in response to JA, resulting in lack of inducibility across species (Figure 11). The species *E. crepidifolium* (CRE) heavily influenced inducibility patterns, as it not only had three times higher constitutive concentrations of cardenolides than any other *Erysimum* species, but in addition pronouncedly increased both glucosinolate and cardenolide concentrations in response to JA treatment (Figure 11). If this outlier was removed, inducibility (or suppression) of foliar cardenolides was not correlated with constitutive cardenolide levels (r = −0.21, p = 0.284), and inducibilities of glucosinolates and cardenolides were likewise not correlated with each other (r = 0.01, p = 0.995).

**Figure 11.**
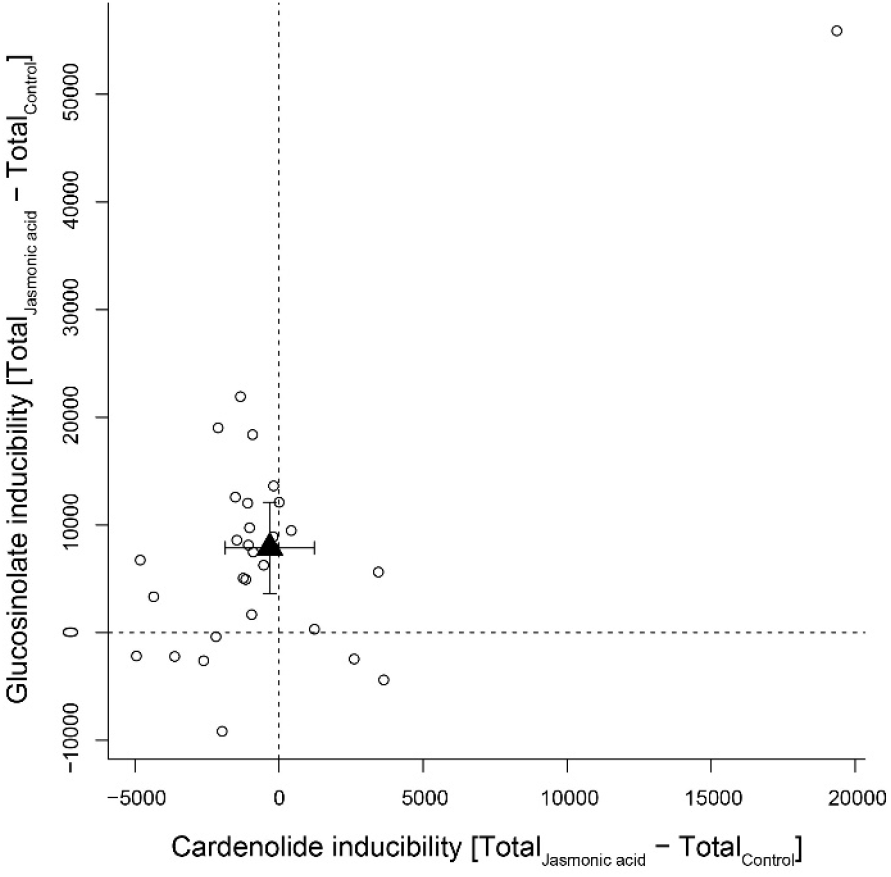
Inducibility of foliar glucosinolates and cardenolides in response to exogenous application of jasmonic acid (JA), expressed as absolute differences in total mass intensity between JA-treated and control plants. Circles are species means, based on single pooled samples of multiple individual plants. The filled triangle is the average inducibility of all measured species with 95% confidence interval. Non-overlap with zero (dashed lines) corresponds to a significant effect. The species in the upper right corner is *E. crepidifolium*, an outlier and strong inducer of both glucosinolates and cardenolides.

## Discussion

The genus *Erysimum* is a fascinating model system of phytochemical diversification that combines two potent classes of chemical defenses in the same plants. The assembled genome of the short-lived annual plant *E. cheiranthoides* allowed us to identify almost the full set of genes involved in *E. cheiranthoides* glucosinolate biosynthesis and myrosinase expression. This genome (www.erysimum.org) will facilitate further identification of glucosinolate genes unique to *Erysimum* and represents a central resource for the identification of cardenolide biosynthesis genes in this emerging model system, as well as for future functional and evolutionary studies in the Brassicaceae.

The extant species diversity in the genus *Erysimum* is the result of an evolutionarily recent, rapid radiation (Moazzeni et al. 2014). All but one species in our study produced evolutionary novel cardenolides, while the likely closest relatives – the genera *Malcolmia*, *Physaria* or Arabidopsis (Moazzeni et al. 2014, Huang et al. 2016) – almost certainly lack these defenses (Jaretzky and Wilcke 1932, Hegnauer 1964). The onset of diversification in *Erysimum* thus appears to coincide with the gain of the cardenolide defense trait. However, even though most species co-expressed two different classes of potentially costly defenses, there was no evidence for a trade-off between glucosinolates and cardenolides. Furthermore, neither defense showed a directional trend from the root to the tip of the phylogenetic tree, and both defenses were similarly diverse in early- and late-diverging species.

Potentially costly, obsolete defenses are expected to be selected against and should disappear over evolutionary time. For example, cardenolides in the genus *Asclepias* and alkaloids across the Apocynaceae decrease in concentration with speciation, consistent with co-evolutionary de-escalation in response to specialized, sequestering herbivores (Agrawal and Fishbein 2008, Livshultz et al. 2018). Due to the relatively recent evolution of cardenolides in *Erysimum*, the system presumably lacks cardenolide-specialized herbivores that could exert negative selection on cardenolides, while the diversity of cardenolides may have evolved too rapidly to detect positive selection. It is therefore likely that the two defenses serve distinct functions: glucosinolates are highly efficient in repelling generalist herbivores (Kerwin et al. 2015), whereas cardenolides may be functionally relevant against glucosinolate-specialized herbivores (Chew 1975, 1977, Wiklund and Åhrberg 1978, Renwick et al. 1989, Dimock et al. 1991).

As further evidence for the distinct roles of glucosinolates and cardenolides, the two defenses responded differently to exogenous JA application. Glucosinolate concentrations were upregulated in response to JA in the majority of species, with an average 52% increase relative to untreated controls. This is similar to inducibility of glucosinolates reported for other Brassicaceae species (Textor and Gershenzon 2009), suggesting that glucosinolate defense signaling remains unaffected by the presence of cardenolides in *Erysimum* plants. In contrast, cardenolide levels were not inducible or were even suppressed in response to exogenous application of JA in almost all tested species, suggesting that inducibility of cardenolides is not a general strategy of *Erysimum*. In the more commonly-studied milkweeds (*Asclepias* spp., Apocynaceae), cardenolides are usually inducible in response to herbivore stimuli (Rasmann et al. 2009, Bingham and Agrawal 2010), but cardenolide suppression is also common, particularly in plants with high constitutive cardenolide concentrations (Bingham and Agrawal 2010, Rasmann and Agrawal 2011). It thus appears that cardenolides accumulate constitutively in *Erysimum*, perhaps due to the presumed lack of cardenolide-specialized herbivores that would use cardenolides as host-finding cues.

### Phylogenetic relationships and phytochemical diversity

The young evolutionary age of *Erysimum* and the potential ongoing gene flow among related species may have limited genetic differentiation. In fact, both previous partial phylogenies of the genus struggled to resolve polytomies among species using conventional approaches (Gómez et al. 2014, Moazzeni et al. 2014). In contrast, we present a highly-resolved species tree for 10 – 30 % of the total species diversity in the genus. Our tree, which was constructed from transcriptome sequences of 9,868 genes with syntenic genome locations, revealed a strong geographic signature in phylogenetic relatedness, an observation shared by the previously published phylogenies (Gómez et al. 2014, Moazzeni et al. 2014). In our species tree, all annual species belonged to early-diverging clades, suggesting that the predominant perennial growth strategy in the genus is a derived state. Three of the annual species, *E. repandum*, *E. incanum*, and *E. wilczekianum*, have widely different distribution ranges but were all collected in Spain or Morocco for this study. Although they co-occur geographically with several perennial *Erysimum* species, they are largely isolated by non-overlapping flowering times.

Perennial species from Spain, Morocco, and the Canary Islands formed a monophyletic clade, with species from southeastern Spain exhibiting closer relatedness to Moroccan species than to species from northeastern or northwestern Spain. Among the species from southeastern Spain, our phylogeny did not agree with a more fine-scale evaluation of species relatedness (Abdelaziz et al. 2014), although this may reflect the limitation of using single accessions of each species in our approach, the possible hybridization occurring between these species (Abdelaziz et al. 2014), or the different sensitivity of phylogenies based on internal transcribed spacer (ITS) sequences to incomplete lineage sorting (Feliner and Rossello 2007). Additional geographic clades were recovered for species from North America, Iran, and Greece/Turkey. Surprisingly, the North American clade was most closely related to the Spanish clade, which does not fit traditional models of dispersal across land bridges. However, the node connecting the two geographic clades had by far the weakest support across the whole phylogeny (local posterior probability = 0.5) and, as our phylogeny did not include East Asian *Erysimum* species, it is possible that the addition of further species would change this grouping. The remaining species belonged to one of three distinct central European clades with no obvious geographic separation. As a prominent exception, the central European *E. crepidifolium* (CRE) was most closely related to Greek *Erysimum* species rather than to other central European species.

Despite vast morphological differences among sampled *Erysimum* species, the diversity in glucosinolate profiles was relatively limited compared to the diversity that is present within Arabidopsis (Kliebenstein et al. 2001). However, broader comparative studies of glucosinolate diversity in other Brassicaceae species would be needed to provide a more natural ‘baseline’ for glucosinolate diversity. The majority of *Erysimum* species produced glucoiberin as their main glucosinolate. Aliphatic glucosinolates such as glucoiberin are derived from methionine in a process that involves elongation and modification of a variable side-chain (Halkier and Gershenzon 2006), and in this context the 3-carbon glucosinolate glucoiberin is one of the least biosynthetically complex glucosinolates. However, the potential to produce additional aliphatic glucosinolates with longer side chains clearly exists in the genus, as 4-, 5-, and 6-carbon glucosinolates with more complex modifications were scattered across the phylogeny. A few species produced glucosinolates that are not found in Arabidopsis, including a sub-class of aliphatic glucosinolates, the methylsulfonyl glucosinolates. The homolog of 3-butenyl glucosinolate 2-hydroxylase (GS-OH), which in Arabidopsis forms 2-hydroxy-but-3-enyl glucosinolate from 3-butenyl glucosinolate, does not have a clear function in *E. cheiranthoides* due to the lack of alkenyl glucosinolates. However, it is possible that the GS-OH homolog in *E. cheiranthoides* may code for the unknown enzyme that hydroxylates 4-methylsulfonylbutyl glucosinolate to form 3-hydroxy-4-methylsulfonylbutyl glucosinolate (Figure 4). Methylsulfonyl glucosinolates are found in several Brassicaceae genera (Fahey et al. 2001), and glucocheirolin, the most abundant methylsulfonyl glucosinolate in *Erysimum* species, is only a weak egg-laying stimulant for the cabbage white butterfly (*Pieris rapae*), compared to other glucosinolates (Huang et al. 1993). Methylsulfonyl glucosinolates may thus represent a plant response to specialist herbivores that use plant defenses as host-finding cues.

The species *E. pulchellum* (PUL) and *E. collinum* (COL) from Turkey and Iran, respectively, accumulated glucoerypestrin as their main glucosinolate compound. This compound was first described in *E. rupestre* [syn. *E. pulchellum*, (Polatschek 2011)] by Kjaer & Gmelin (1957) and to date has been found exclusively in plants of the genus *Erysimum* (Fahey et al. 2001). Radioactive labeling experiments indicated that glucoerypestrin is derived from a dicarboxylic amino acid, possibly 2-amino-5-methoxycarbonyl-pentanoic acid (Chisholm 1973). Modification of the amino acid side chain during methionine-derived aliphatic glucosinolate biosynthesis as a pathway to glucoerypestrin is less likely, due to the lower specific incorporation of ^14^C-labeled methionine compared to ^14^C-labeled dicarboxylic acids into this compound (Chisholm 1973). In any case, the gain of glucoerypestrin represents yet another evolutionary novelty in the *Erysimum* genus, but its relative toxicity and the adaptive benefits of its production have yet to be elucidated.

We found no phylogenetic signal of glucosinolate chemotype, as more closely related species were not more likely to share the same glucosinolate profile. The pattern of more complex glucosinolates scattered across the phylogenetic tree may be generated by horizontal gene transfer, during hybridization, or by repeated gains and losses of biosynthesis genes as species diverge. The latter may also be facilitated by changes in ploidy, as hexaploid species in particular accumulated large numbers of both glucosinolate and cardenolide compounds. Alternatively, a full complement of synthesis genes may be maintained in species’ gene pools at low frequencies until they are favored by a new environment, or they might be maintained in the genome but not expressed in leaves. More extensive sampling within each species will be required to conclusively address this question, although a preliminary screening of multiple *E. cheiranthoides* accessions suggests little to no variation in glucosinolate profiles within this species (T. Züst, unpublished data).

Myrosinase activity levels differed among glucosinolate chemotypes, and activity was positively correlated with glucosinolate abundance in plants when controlling for glucosinolate chemotype. *Erysimum* species that predominantly produced indole glucosinolates or 4-methylsulfinyl glucosinolates had negligible myrosinase activity against the assayed aliphatic glucosinolate sinigrin. Indole glucosinolates can be activated by PEN2 – a thioglucosidase that is more specific for indole glucosinolates (Bednarek et al. 2009, Clay et al. 2009) – or even break down in the absence of plant-derived myrosinase (Kim et al. 2008). The negligible activity in these species could therefore indicate the existence of selective pressures to tailor myrosinase expression to the type and concentrations of glucosinolates that are produced. In contrast to glucosinolate defenses, myrosinase activity was more similar among related species, suggesting that the two defense components are subject to different selective regimes, with the potential for maladaptive combinations between glucosinolate defense and myrosinase activity. In addition, myrosinase activity is highly dependent on the presence of other proteins and cofactors (Halkier and Gershenzon 2006), which may also differ between *Erysimum* species.

We detected considerable amounts of the evolutionarily novel cardenolide defense in 47 out of 48 *Erysimum* species or accessions. Among the 95 likely cardenolide compounds, there were several structures that had not been described previously in *Erysimum*. This metabolic diversity had three main sources: modification of the genin core structure, variation of the glycoside chain, or isomeric variation (e.g., through the incorporation of different isomeric sugars). Structural variation in cardenolides affects the relative inhibition of Na^+^/K^+^-ATPase (Dzimiri et al. 1987, Petschenka et al. 2018) and physiochemical properties such as lipophilicity, which play an important role in uptake and metabolism of plant metabolites by insects (Duffey 1980). Individual *Erysimum* species produced between 15 and 50 different cardenolide compounds, and the comparison of quantification by total mass ion counts *vs.* quantification by inhibition of Na^+^/K^+^-ATPase revealed highly similar results. While both methods of quantification are only approximate, this correspondence at least provides no obvious indication of vast differences in Na^+^/K^+^-ATPase inhibitory activity among *Erysimum* cardenolides.

The metabolic pathways involved in the biosynthesis and modification of cardenolides have yet to be elucidated (Kreis and Müller-Uri 2010, Züst et al. 2018). Here, we propose a pathway for the modification of digitoxigenin, commonly assumed to be the least biosynthetically complex cardenolide (Kreis and Müller-Uri 2010), into the eight structurally more complex genins found within *Erysimum* (Figure 9). Variation in glycoside chains is likely mediated by glycosyltransferases that act on the different genins. In the Brassicaceae genus *Barbarea*, plants produce saponin glycosides as an evolutionary novel defense, and a significant proportion of glycoside diversity in this system has been linked to the action of a small set of UDP glycosyltransferases (Erthmann et al. 2018). Similarly, through the joint action of genin-modifying enzymes and glycosyltransferases, a relatively small set of enzymes and corresponding genes could generate the vast cardenolide diversity found in the *Erysimum* genus. The identification and manipulation of these genes in different *Erysimum* species will make it possible to test the adaptive benefits of this structural diversity.

On average, leaves of *Erysimum* species contained cardenolides equivalent to 6 µg ouabain per mg dry leaf weight (estimated from Na^+^/K^+^-ATPase inhibition), placing them slightly above most species of the well-studied cardenolide-producing genus *Asclepias* (Rasmann and Agrawal 2011). However, two species, *E. collinum* (COL) and *E. crepidifolium* (CRE), were clear outliers in terms of cardenolide content (Figure 6D-E). The almost complete absence of cardenolides in *E. collinum* (COL), which clustered phylogenetically with two other Middle Eastern species producing average concentrations of these compounds (*E. crassipes* [CSS] and *E. crassicaule* [CRA], Figure 5), likely represents a secondary loss of this trait in the course of evolution. This species also accumulated an evolutionary novel glucosinolate, glucoerypestrin (see above), which may have resulted in a shift in selective pressures that led to the loss of potentially costly cardenolide production. Conversely, *E. crepidifolium* (CRE) had cardenolide concentrations more than three times higher than any other tested *Erysimum* species. This is consistent with the highly toxic nature of this species, which has the German vernacular name ‘Gänsesterbe’ (geese death) and has been associated with mortality in geese that consume the plant.

Whereas most species did not induce cardenolide accumulation in response to JA, *E. crepidifolium* (CRE) had a significant 48% increase. While not as extreme, this observation is similar to the results of Munkert et al. (2014), who reported a three-fold increase in cardenolide levels of *E. crepidifolium* in response to methyl jasmonate application. Plants use conserved transcriptional networks to continuously integrate signals from their environment and optimize allocation of resources to growth and defense (Havko et al. 2016). Thus, while these networks commonly govern hardwired responses (e.g., an attenuation of growth upon activation of JA signaling), they may nevertheless be altered by mutations at key nodes of the network (Campos et al. 2016). Given this relative flexibility in signaling networks, it is perhaps not surprising that the evolutionary novel cardenolides have been integrated into the defense signaling of *Erysimum* species to variable degrees. Investigating gene expression changes in the inducible *E. crepidifolium* as a contrast to the non-inducing *E. cheiranthoides* may therefore provide valuable insights into the molecular regulation of this defense.

Cardenolide abundance and compound profiles of *Erysimum* species exhibited clear phylogenetic signals, with closely-related species being more phytochemically similar. However, similarities in cardenolide profiles changed more gradually between species than glucosinolate profiles, and distinct cardenolide chemotypes were less obvious. As the most distinct cardenolide cluster with underlying phylogenetic structure, the annual *E. cheiranthoides* (ECE) grouped together with *E. sylvestre* (SYL) and two accessions of commercial origin. These plants all shared a cardenolide chemotype defined by an unusually high proportion of digitoxigenin glycoside compounds, several of which were uniquely produced by plants of this cluster. This early-diverging species clade, which is defined by a chemotype of potentially lower biosynthetic complexity, could thus be an indication of a stepwise gain of structural complexity over the course of evolution.

### Conclusions

The study of the speciose genus *Erysimum* with two co-expressed chemical defense classes revealed largely independent evolution of the ancestral and the novel defense. With no evidence for trade-offs between the structurally and biosynthetically unrelated defenses, the diversity, abundance, and inducibility of each class of defenses appears to be evolving independently in response to the unique selective environment of each individual species. The evolutionarily recent gain of novel cardenolides has resulted in a system in which no known specific adaptations to cardenolides have evolved in insect herbivores, although general adaptations to toxic food may still allow herbivores to consume the plants. *Erysimum* is thus an ideal model system for phytochemical diversification, as it facilitates the study of coevolutionary adaptations in real time. Our current work provides the foundation for a more mechanistic evaluation of these processes, which promises to greatly improve our understanding of the role of phytochemical diversity for plant-insect interactions.

## Acknowledgments

We thank Hartmut Christier for collecting *E. cheiranthoides* seeds, Erik Poelman for collecting *E. cheiri* seeds, and the botanical gardens listed in Table S1 for providing seeds of additional species. Yvonne Künzi and Christoph Zwahlen assisted with the growing and maintenance of plants in the 48-species experiments, and with the harvesting and extraction of RNA and metabolomics samples. Sabrina Stiehler performed the Na^+^/K^+^-ATPase assay, and Jing Zhang helped with database maintenance. We thank Anurag Agrawal, Hamid Moazzeni, Gaurav Moghe, and Zephyr Züst for helpful advice and comments on the manuscript. This work was supported by Swiss National Science Foundation grant PZ00P3-161472 to TZ, a Triad Foundation grant to SRS and GJ, US National Science Foundation awards 1811965 to CKH and 1645256 to GJ, a fellowship from the Ministry of Science of Iran to MM, a German Research Foundation grant DFG-PE 2059/3-1 to GP, and a grant within the LOEWE program (Insect Biotechnology & Bioresources) of the State of Hesse, Germany to GP.

## Supplementary Figures and Tables

**Supplementary Figure S1.**
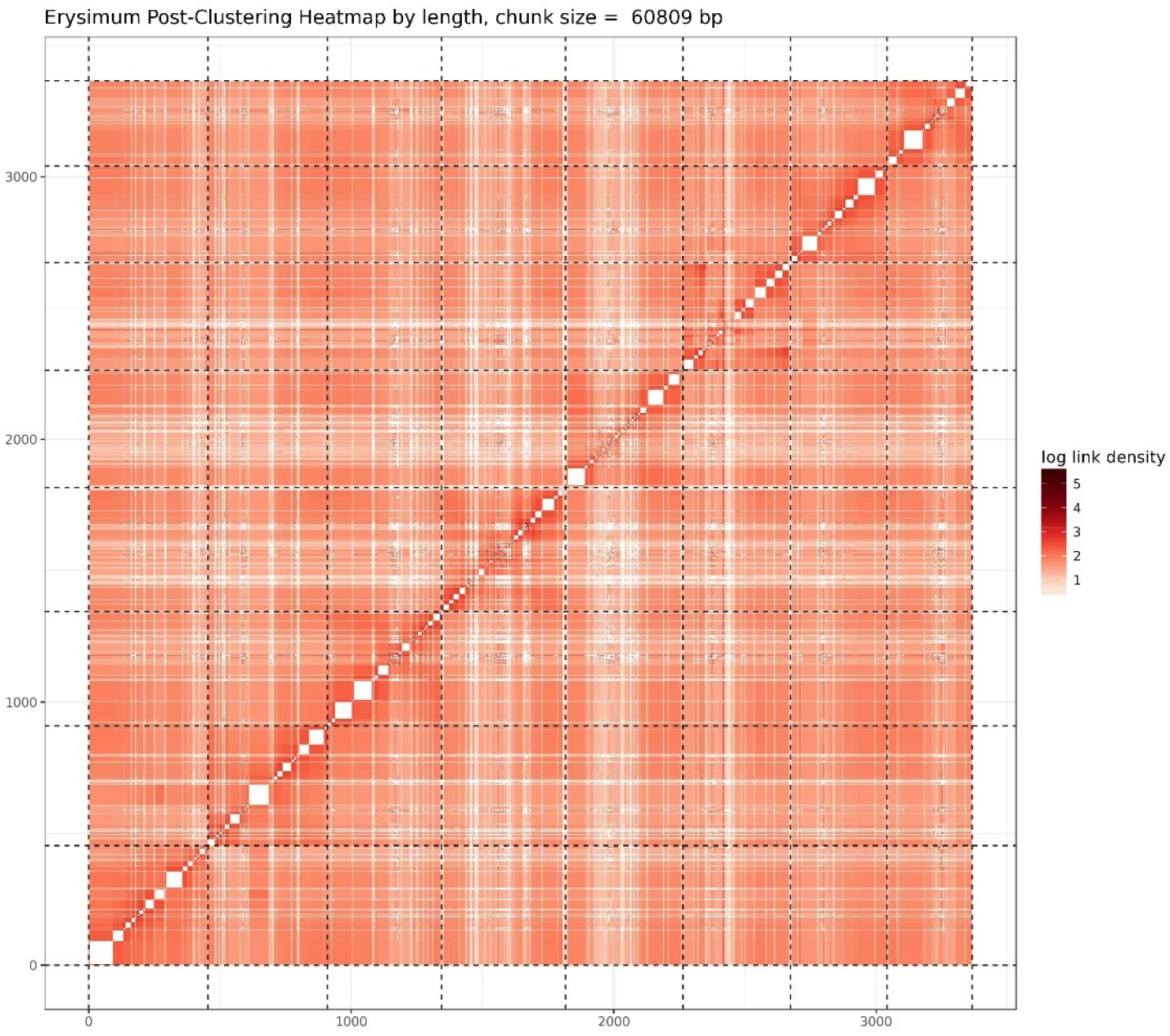
Post-clustering heatmap showing the density of Hi-C interactions between scaffolds used in assembly. Intensity corresponds to the total number of reads per interaction. Dashed lines delimit the eight identified pseudomolecules.

**Supplementary Figure S2.**
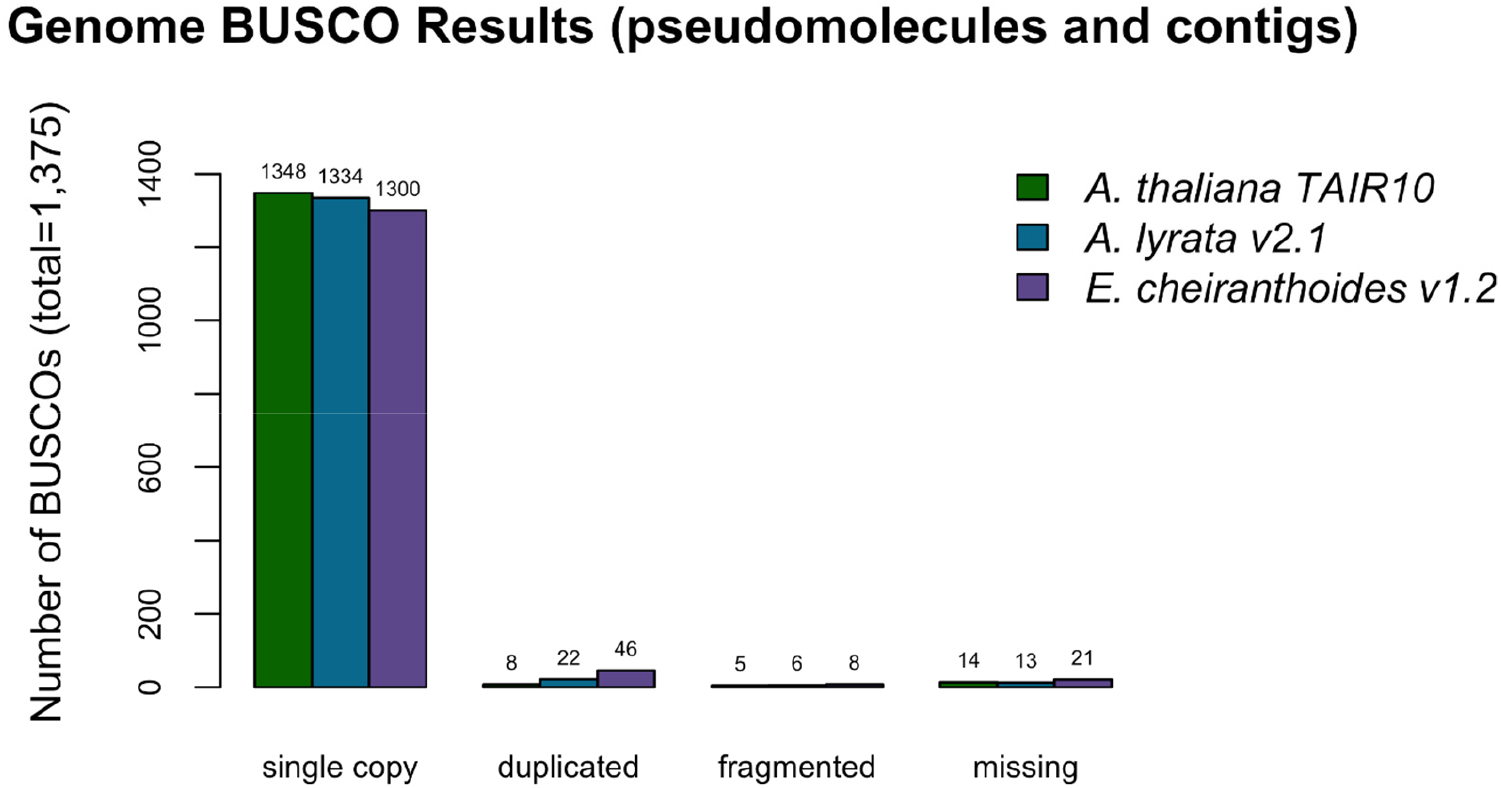
BUSCO completeness assessment for the EC1.2 genome assembly. *A. thaliana* and *A. lyrata* results provided for comparison to EC1.2 genome assembly.

**Supplementary Figure S3.**
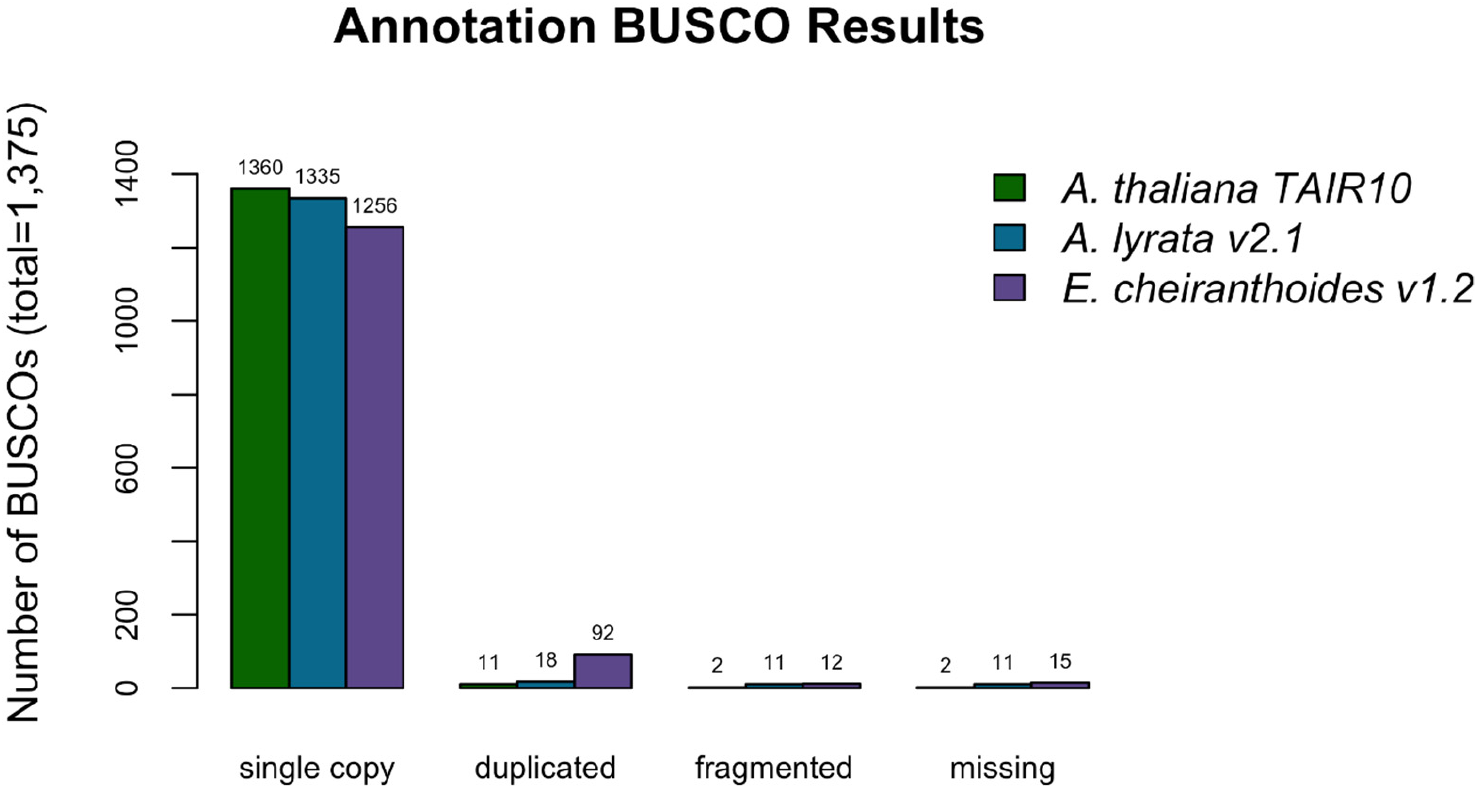
BUSCO completeness assessment for the EC1.2 genome annotation. *A. thaliana* and *A. lyrata* results provided for comparison to the EC1.2 genome annotation.

**Supplementary Figure S4.**
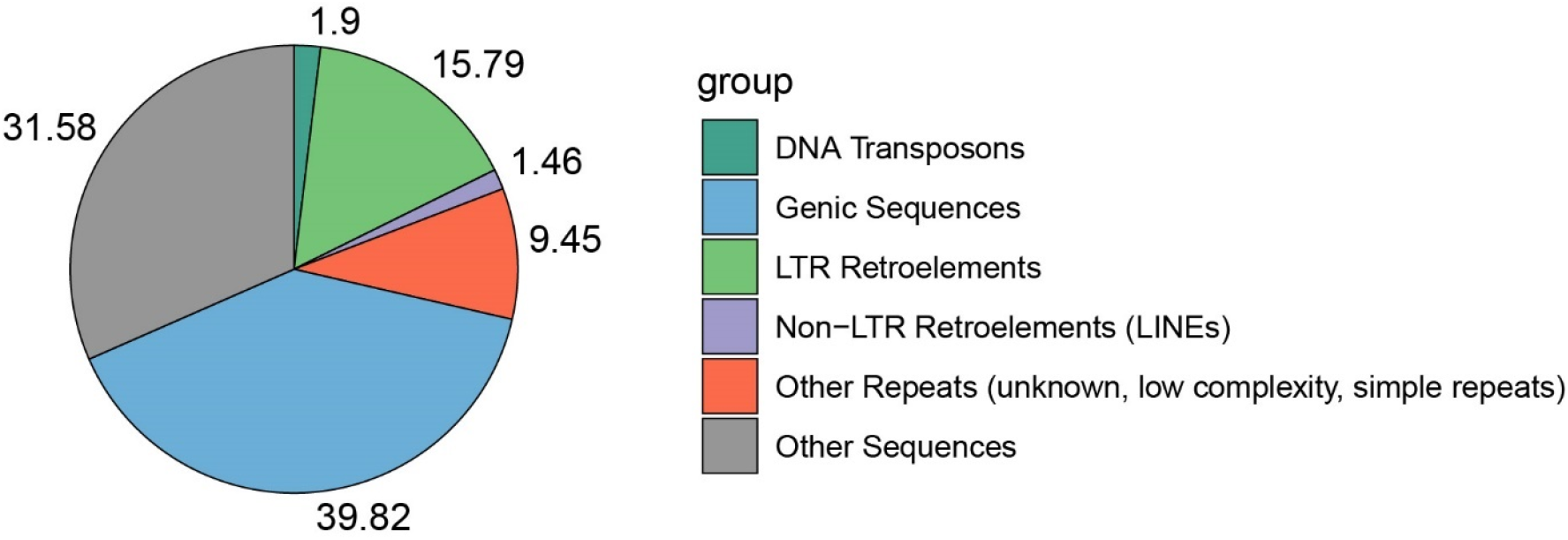
Proportional contributions of sequence classes to the genome sequence of *E. cheiranthoides*.

**Supplementary Figure S5.**
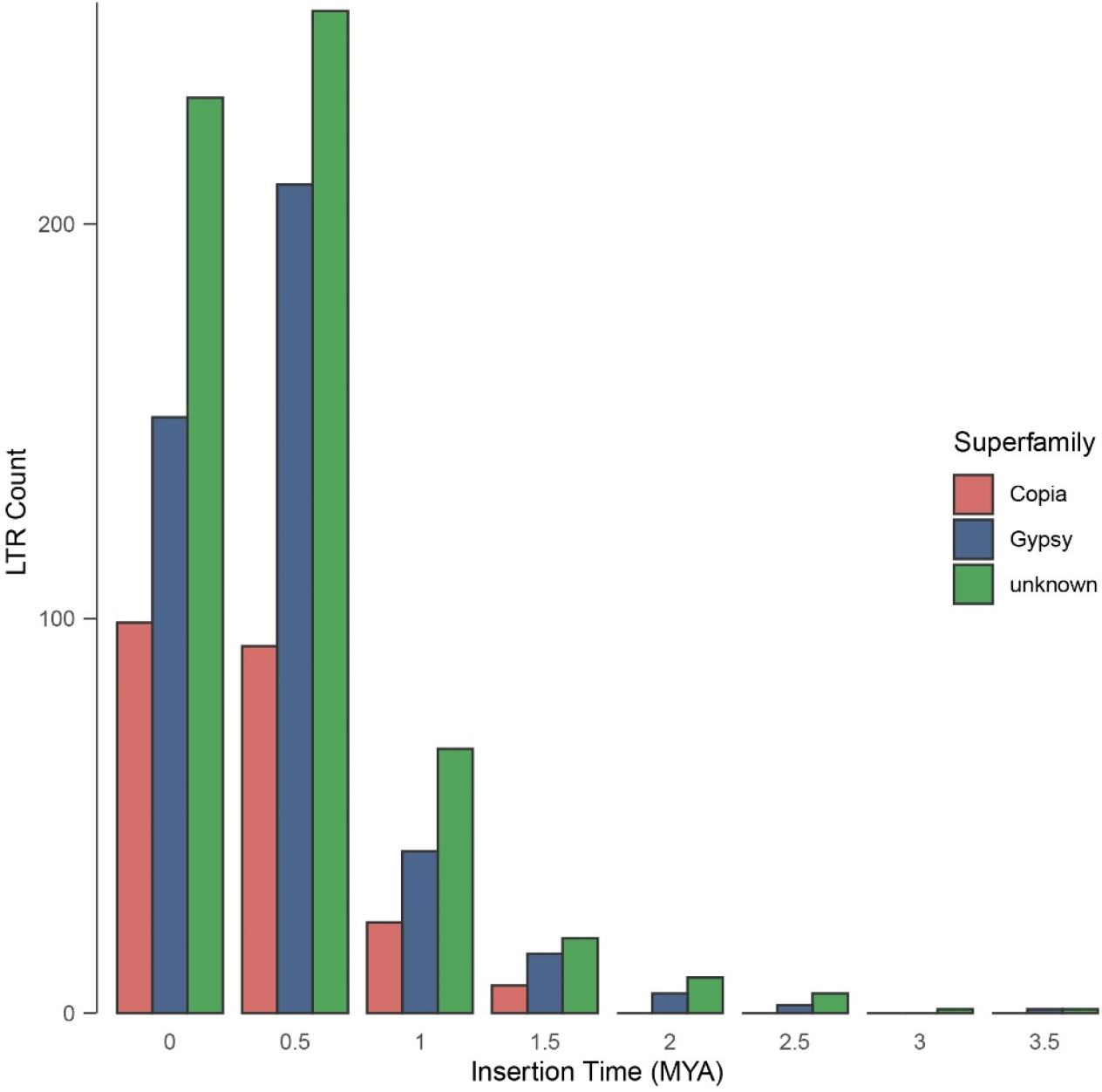
Age distributions of intact LTR-RTs identified by LTR_retriever in the genome of *E. cheiranthoides*.

**Supplementary Figure S6.**
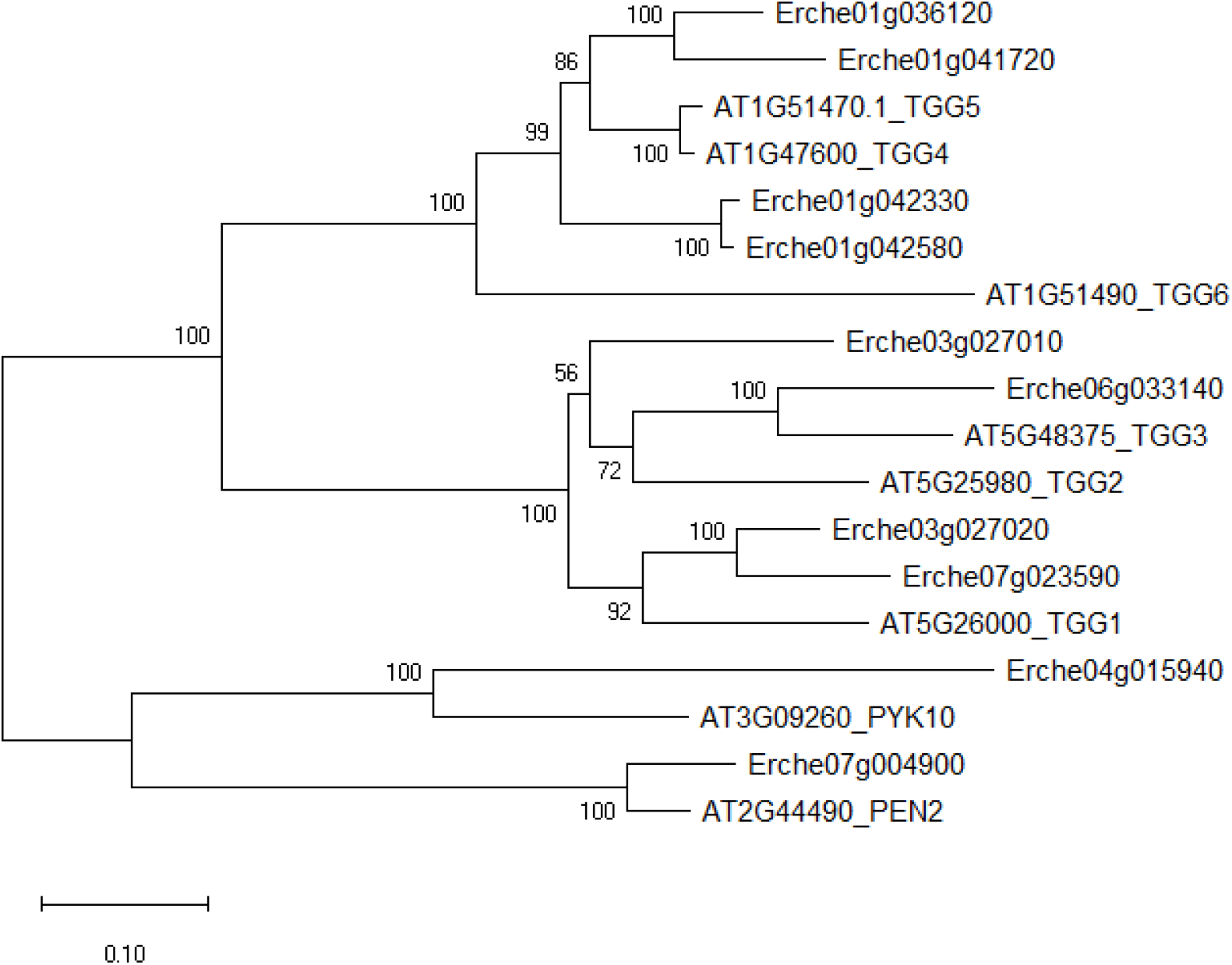
Phylogenetic analysis of myrosinase genes in *E. cheiranthoides* and *A. thaliana* using neighbor-joining methods. Nodes are labelled with bootstrap tests based on 1000 replicates.

**Supplementary Figure S7.**
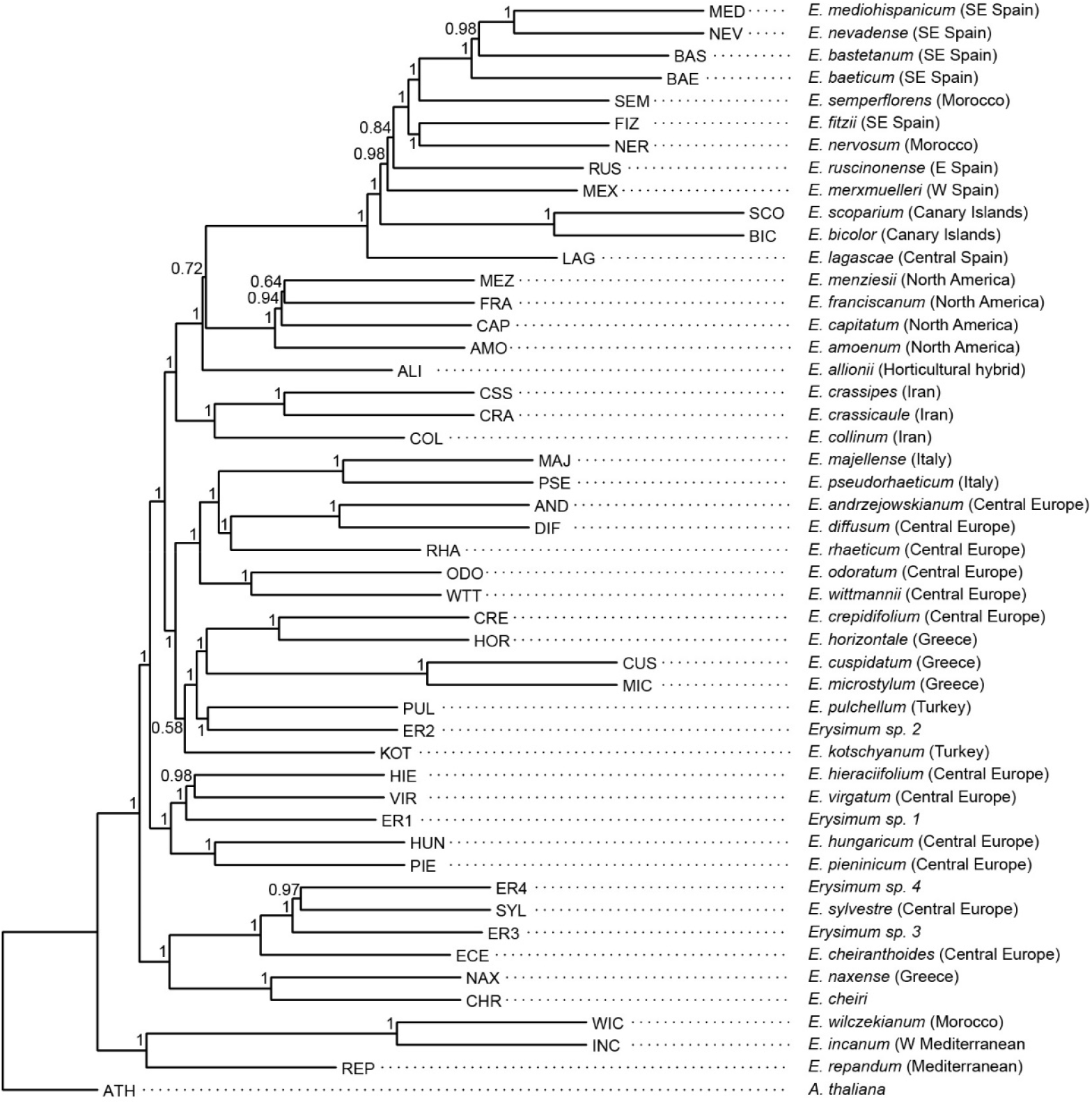
Coalescent species tree inferred from 2,306 orthologous gene sequences. Nodes are labelled with local posterior probability, indicating level of support.

**Supplementary Figure S8.**
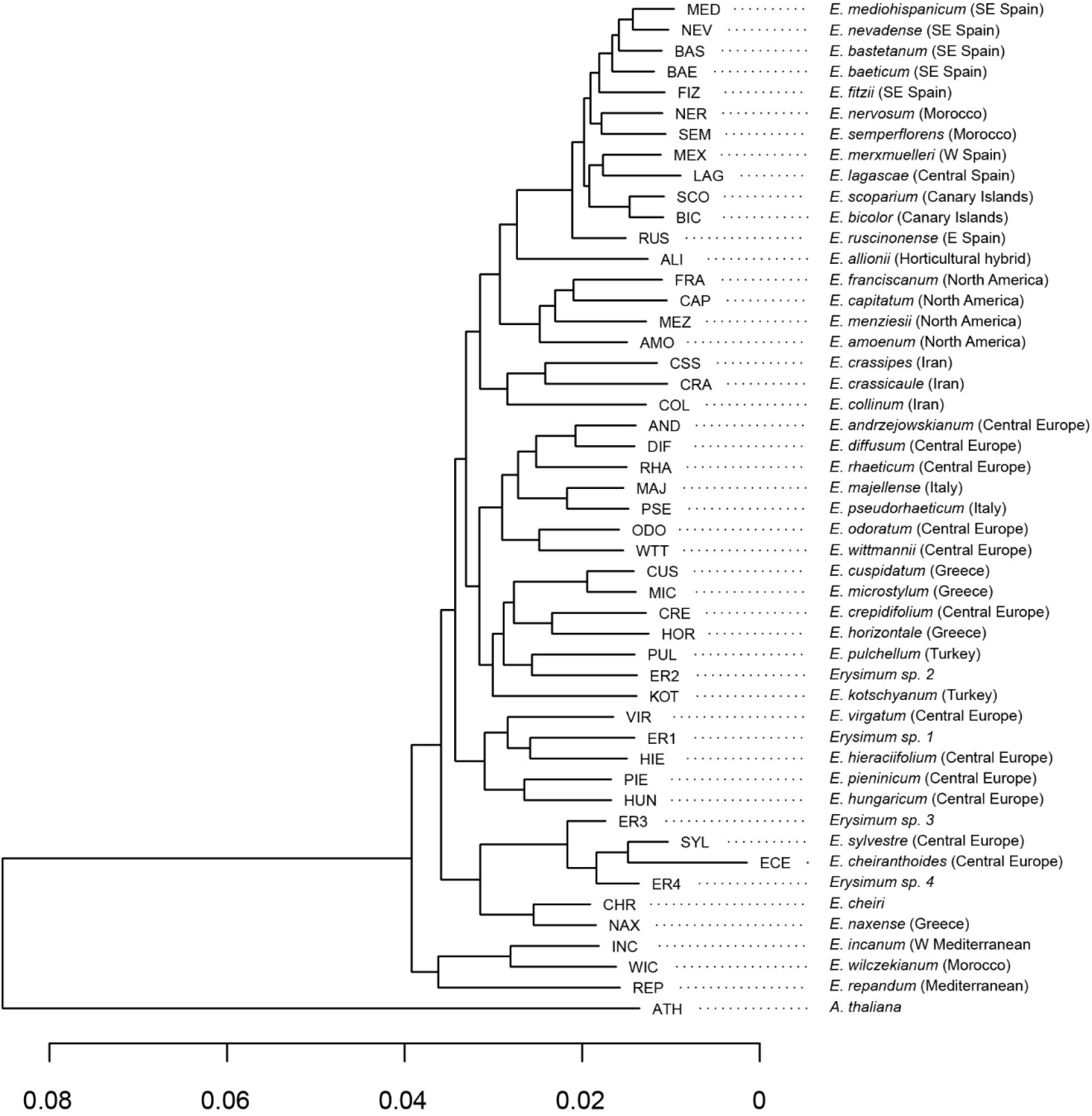
Concatenated species tree inferred from 2,306 orthologous gene sequences. Branch length corresponds to estimated number of substitutions per site.

**Supplementary Figure S9.**
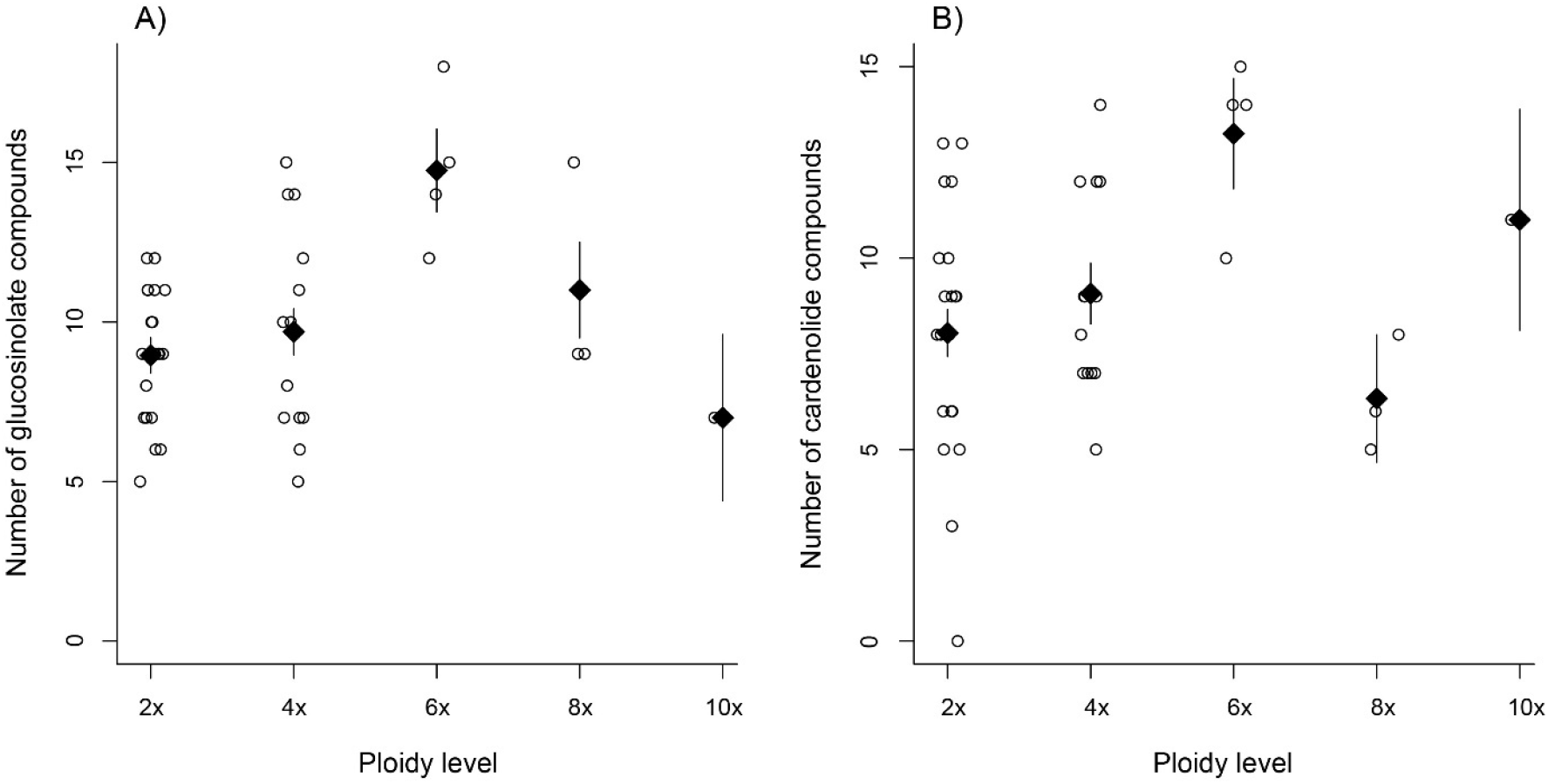
Effect of ploidy on compound diversity. Open circles correspond to species, with ploidy inferred from literature reports. Black triangles are mean values ± 1 SE for each ploidy level. (A) Total number of glucosinolate compounds produced by each *Erysimum* species. (B) Number of cardenolide compounds which together constitute 80% of total cardenolide concentrations. As many cardenolide compounds were produced at very low concentrations, an effect of ploidy was obscured if the total number of cardenolide compounds was considered.

**Supplementary Figure S10.**
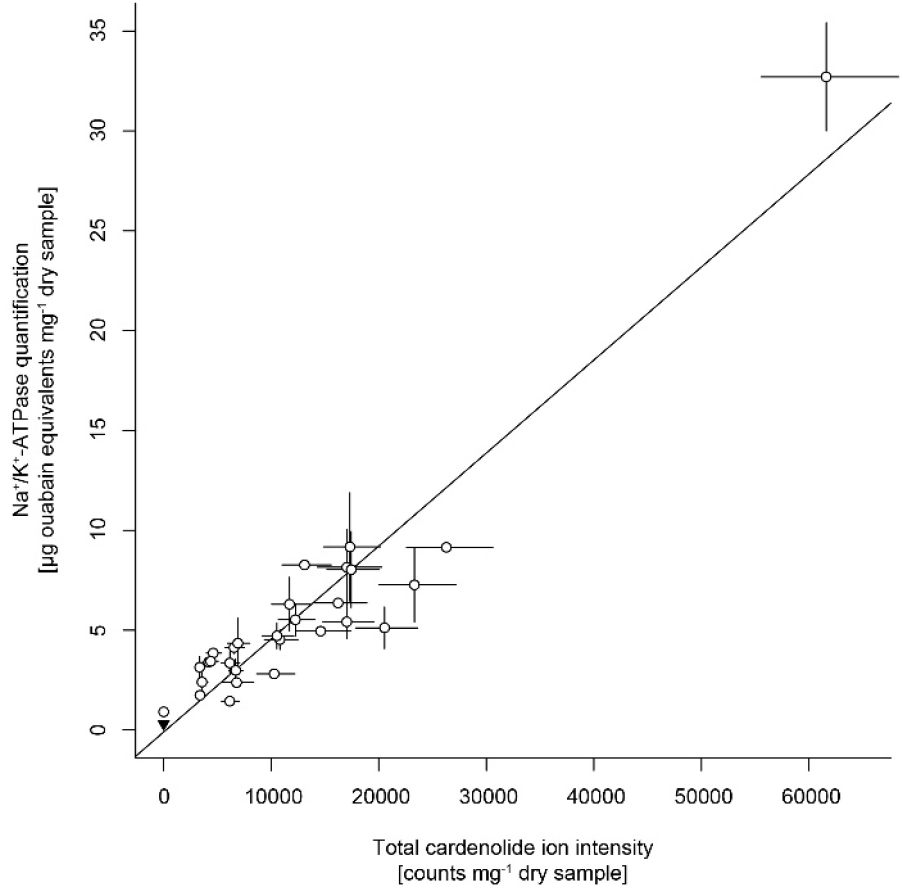
Correlation between cardenolide concentrations approximated by total cardenolide ion intensity and by inhibition of animal Na^+^/K^+^-ATPase. Open circles are species means ± 1 SE. The black triangle in the bottom left corner is the quantification of *Sinapis arvensis* tissue as a negative control. The solid line is the linear regression on species means.

**Supplementary Figure S11.**
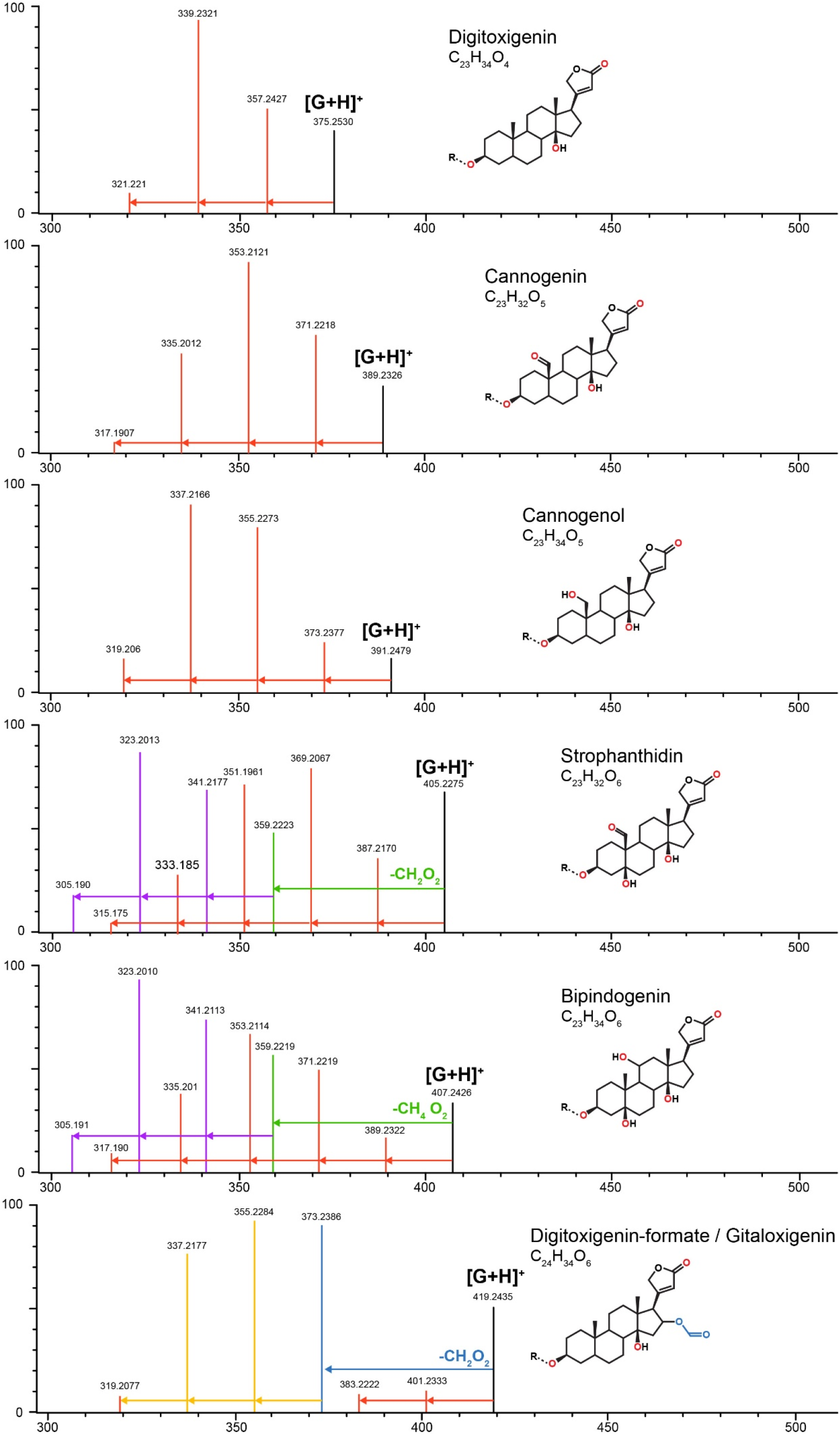
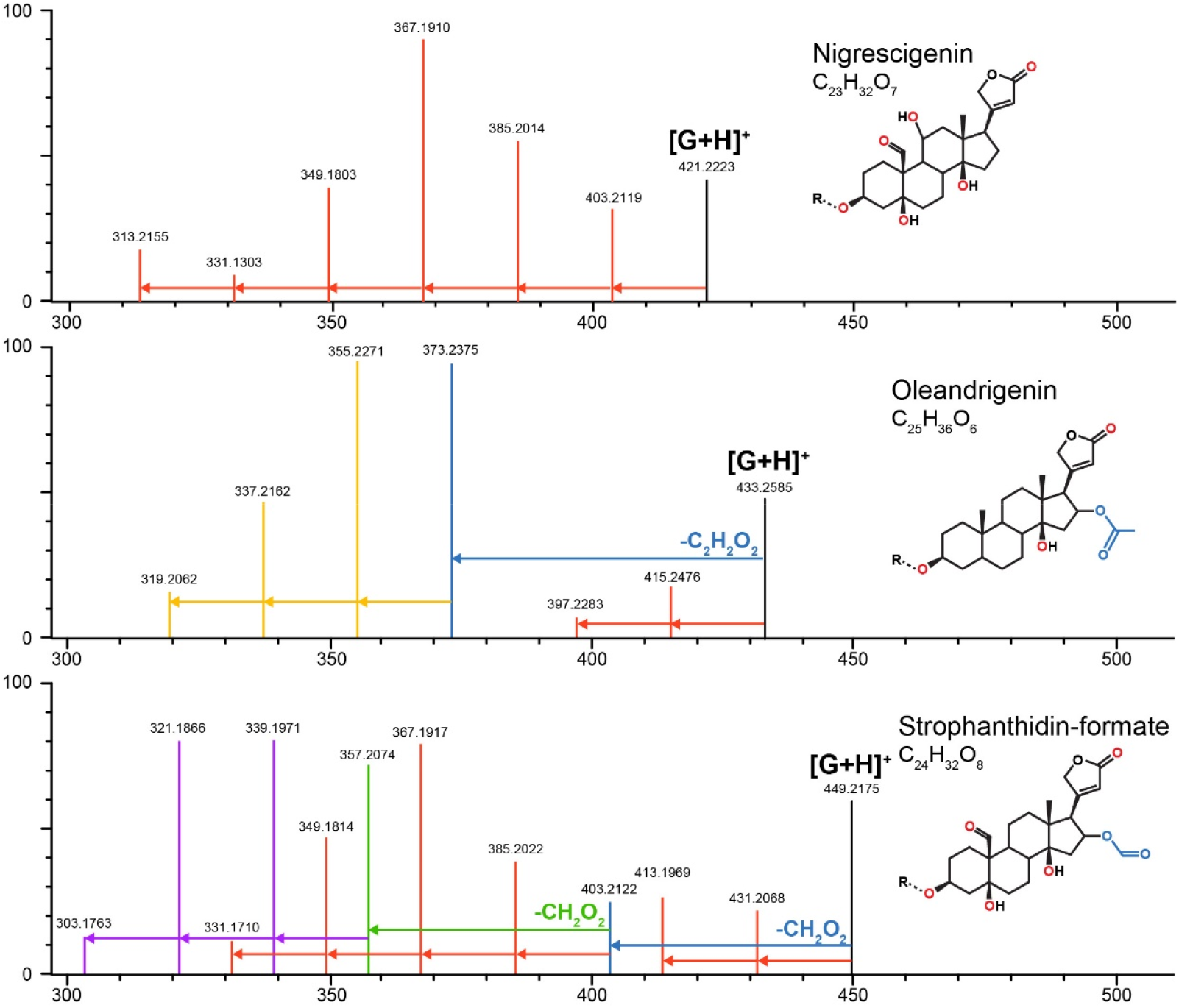
Characteristic MS fragmentation patterns of cardenolide genins used for putative identification of compounds. Patterns are primarily generated by in-source fragmentation of compounds and are visible at standard MS conditions, but fragments are more abundant under MS^E^ (high energy fragmentation) conditions. Height of vertical lines indicates relative ion intensities (in MS^E^) for representative compounds quantified in *Erysimum* spp. The mass and intensity of the intact genin ([G+H]^+^) in each panel is indicated by a vertical black line. Genin fragments are colored to highlight fragmentation series: red vertical lines linked by arrows represent serial losses of water molecules (−18.01 m/z per molecule), corresponding to the number of exposed oxygen groups of that molecule (red symbols). Acetyl and formate groups are lost as intact units (blue lines/arrows and corresponding symbols), after which further loss of water molecules occurs (orange lines/arrows). In strophanthidin and bipindogenin, additional larger fragments are lost, perhaps by reconfiguration of the genin molecule (green lines/arrows), and again, further loss of water molecules occurs (purple lines/arrows). Fragmentation patterns for strophanthidin and digitoxigenin were confirmed by commercial standards. For remaining genins, likely identifications and structures were inferred from literature reports.

**Supplementary Figure S12.**
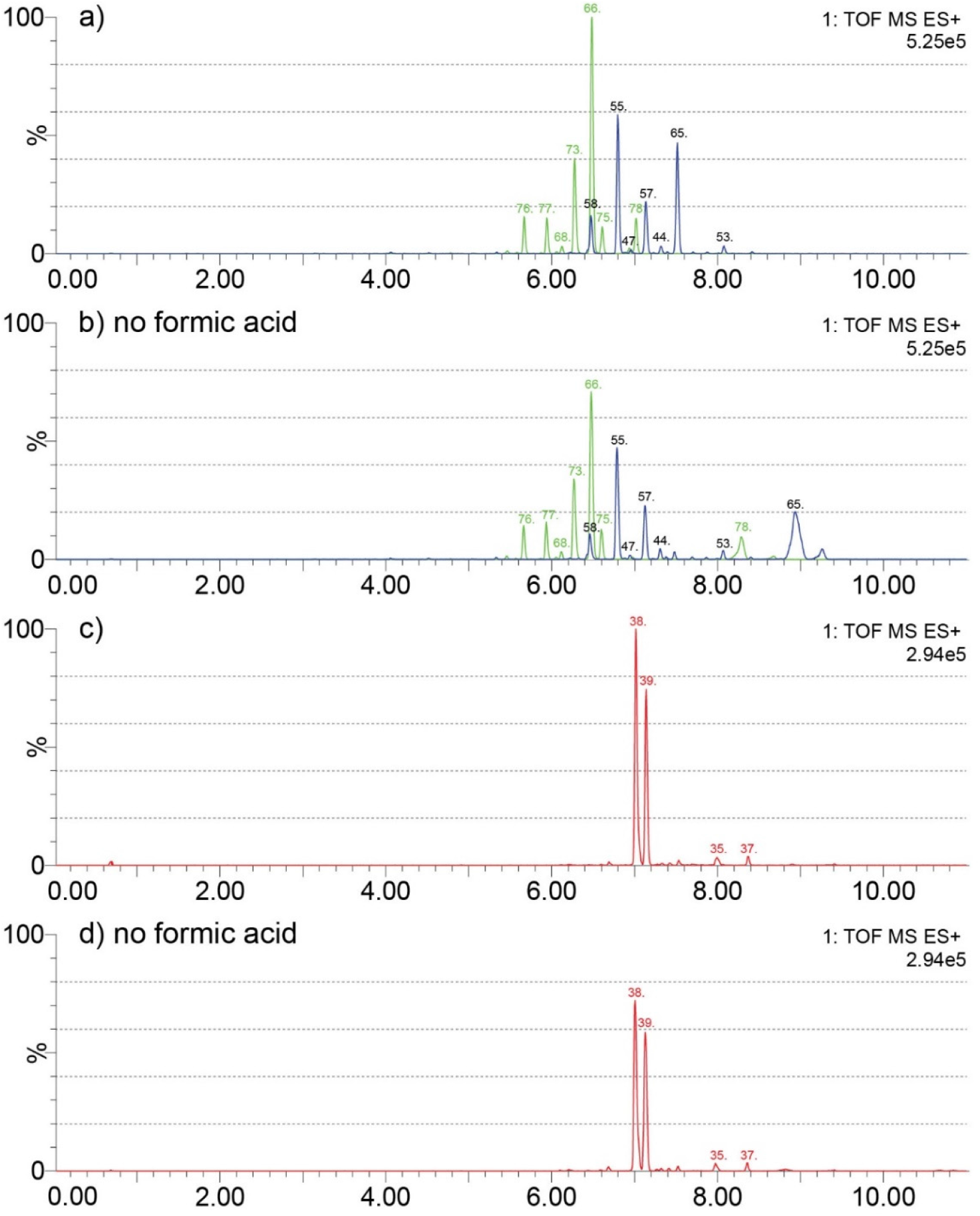
Effects of formic acid as a solvent additive in LC-MS analyses of cardenolide compounds. A-B) Chromatograms for the same leaf extract of *E. bastetanum* (BAS) analyzed with and without formic acid in the mobile phase. Shown are fragment traces for strophanthidin-formate (449.217 m/z, green) and strophanthidin (405.227 m/z, blue). Both panels share the same scale, and compounds are labelled according to the list in Supplementary Table S3. Without formic acid, overall intensity is reduced, but glycosides of the strophanthidin-formate genin are consistently detected. In contrast, two compounds with unusual adducts on their glycoside chain (#65 and #78, Table S5) appear to be changed by formic acid and were removed from the analysis. C-D) Chromatograms for the same leaf extract of *E. repandum* (REP) analyzed with and without formic acid in the mobile phase. Shown are fragment traces for digitoxigenin-formate (419.243 m/z, red). Both panels share the same scale, and compounds are labelled according to the list in Supplementary Table S3.

**Supplementary Table S1.** Origin of species and seed material, and year of original collection where available. Four species from mislabeled seed stocks are referred to as accessions ER1-4, with false species name provided in quotation marks. Ploidy levels can be variable within species and are based on measurements of the sampled populations where available (highlighted by *), or otherwise inferred from literature reports. Leaf material for each species or accession was collected in one of three experiments, and RNA was extracted from pooled tissue of 2-5 individual plants that were sampled at 1-2 time points (TP).

| Species code | Species | Region/country of origin | Seed origin | Original collection | Ploidy | Source experiment | Pooled individuals |
| --- | --- | --- | --- | --- | --- | --- | --- |
| ALI | *E. allionii*<br>(*E. x. marshallii*<sup>1</sup>) | n/a | Botanical Garden Kiel, Germany | 1985 | n/a | 2017-2 | 5 / 2TP |
| AMO | *E. amoenum* | Colorado, USA | Alplains, CO, USA | 2014 | 4x | 2017-1 | 5 / 1 TP |
| AND | *E. andrzejowskianum* | Austria | Collected | 2009 | 10x | 2017-1 | 5 / 2TP |
| BAE | *E. baeticum* | Spain | Collected | 2006 | 8x* | 2016 | 5 / 1TP |
| BAS | *E. bastetanum* | Spain | Collected | 2007 | 6x* | 2017-2 | 5 / 2TP |
| BIC | *E. bicolor* | Canary Islands, Spain | Botanical Garden Konstanz, Germany | unknown | 4x | 2017-2 | 5 / 2TP |
| CAP | *E. capitatum* | USA | B&T World Seeds, France | unknown | 4x | 2017-1 | 4 / 2TP |
| CHR | *E. cheiri* | Netherlands | Collected | 2016 | 2x | 2017-2 | 5 / 2TP |
| COL | *E. collinum*<sup>2</sup> | Iran | Botanical Garden Madrid, Spain | 1993 | 2x | 2017-1 | 3 / 1TP |
| CRA | *E. crassicaule* | Iran | Botanical Garden Madrid, Spain | unknown | 2x | 2017-2 | 2 / 2TP |
| CRE | *E. crepidifolium* | Czechia | Botanical Garden Berlin-Dahlem, Germany | 1996 | 2x | 2017-1 | 5 / 2TP |
| CSS | *E. crassipes* | Iran | Botanical Garden Madrid, Spain | 1993 | 2x | 2017-2 | 5 / 2TP |
| CUS | *E. cuspidatum* | Greece | Botanical Garden Berlin-Dahlem, Germany | 1980 | 2x | 2017-2 | 5 / 2TP |
| DIF | *E. diffusum* | Czechia | Botanical Garden Plzen, Czechia | 2013 | 4x | 2017-2 | 5 / 2TP |
| ECE | *E. cheiranthoides* | Germany | Collected | 2015 | 2x* | 2017-1 | 5 / 1TP |
| ER1 | *Erysimum* sp. 1<br>'*E. crepidifolium*' | n/a | Botanical Garden Berlin-Dahlem, Germany | unknown | n/a | 2017-1 | 4 / 1TP |
| ER2 | *Erysimum* sp. 2<br>'*E. rhaeticum*' | n/a | Botanical Garden Nantes, France | unknown | n/a | 2017-2 | 5 / 2TP |
| ER3 | *Erysimum* sp. 3<br>'*E. asperum*' | n/a | B&T World Seeds, France | unknown | n/a | 2017-1 | 5 / 2TP |
| ER4 | *Erysimum* sp. 4<br>'*E. suffrutescens*' | n/a | B&T World Seeds, France | unknown | n/a | 2017-2 | 5 / 2TP |
| FIZ | *E. fitzii* | Spain | Collected | 2008 | 2x* | 2017-2 | 2 / 2TP |
| FRA | *E. franciscanum* | California, USA | B&T World Seeds, France | unknown | 4x | 2017-1 | 5 / 2TP |
| HIE | *E. hieraciifolium* | Romania | Botanical Garden Jibou, Romania | unknown | 4x | 2017-2 | 5 / 2TP |
| HOR | *E. horizontale* | Greece | Botanical Garden Berlin-Dahlem, Germany | unknown | 4x | 2016 | 4 / 1TP |
| HUN | *E. hungaricum* | Romania | Botanical Garden Jibou, Romania | unknown | 6x | 2017-2 | 5 / 2TP |
| INC | *E. incanum* | Spain | Botanical Garden Madrid, Spain | unknown | 2x | 2017-2 | 4 / 2TP |
| KOT | *E. kotschyianum* | Turkey | Botanical Garden Tübingen, Germany | unknown | 2x | 2017-2 | 5 / 2TP |
| LAG | *E. lagascae* | Spain | Botanical Garden Madrid, Spain | unknown | 2x* | 2017-1 | 5 / 2TP |
| MAJ | *E. majellense* | Italy |  | unknown | 4x | 2016 | 5 / 1TP |
| MED | *E. mediohispanicum* | Spain | Collected | 2007 | 2x* | 2017-2 | 5 / 2TP |
| MEX | *E. merxmuelleri* | Spain | Collected | 2007 | 2x* | 2017-2 | 5 / 2TP |
| MEZ | *E. menziesii* | California, USA | B&T World Seeds, France | unknown | 4x | 2017-1 | 5 / 2TP |
| MIC | *E. microstylum* | Greece | Botanical Garden Berlin-Dahlem, Germany | 1980 | 2x | 2017-1 | 5 / 2TP |
| NAX | *E. naxense* | Greece | Balkan Botanic Garden of Korussia, Greece | unknown | 2x | 2016 | 5 / 1TP |
| NER | *E. nervosum* | Morocco | Collected | 2008 | 4x* | 2017-1 | 5 / 1TP |
| NEV | *E. nevadense* | Spain | Collected | 2007 | 2x* | 2017-2 | 3 / 2TP |
| ODO | *E. odoratum* | Austria | Botanical Garden Bern, Switzerland | 2015 | 4x | 2017-1 | 5 / 2TP |
| PIE | *E. pieninicum* | Romania | Botanical Garden Jibou, Romania | unknown | 6x | 2017-2 | 5 / 2TP |
| PSE | *E. pseudorhaeticum* | Italy | Botanical Garden Nantes, France | unknown | 2x | 2017-2 | 5 / 2TP |
| PUL | *E. pulchellum* | Turkey | Botanical Garden Kursk, Russia | unknown | 8x | 2017-2 | 5 / 2TP |
| REP | *E. repandum* | Spain | Botanical Garden Madrid, Spain | unknown | 2x | 2017-2 | 5 / 2TP |
| RHA | *E. rhaeticum* | Switzerland | Collected | 2016 | 8x | 2017-1 | 5 / 2TP |
| RUS | *E. ruscinonense* | Spain | Collected | 2007 | 2x | 2016 | 5 / 1TP |
| SCO | *E. scoparium* | Canary Islands, Spain | B&T World Seeds, France | unknown | 4x | 2017-1 | 4 / 1TP |
| SEM | *E. semperflorens* | Morocco | Collected | 2008 | 2x | 2017-1 | 5 / 2TP |
| SYL | *E. sylvestre* | Slovenia | B&T World Seeds, France | unknown | 2x | 2017-1 | 5 / 2TP |
| VIR | *E. virgatum* | Switzerland | Botanical Garden St Gallen, Switzerland | 1962 | 6x | 2017-2 | 5 / 2TP |
| WIC | *E. wilczekianum* | Morocco | Collected | 2008 | 4x* | 2017-2 | 5 / 2TP |
| WTT | *E. wittmannii* | Slovenia | Botanical Garden Berlin-Dahlem, Germany | 2010 | 2x | 2017-1 | 5 / 2TP |

<sup>1</sup>horticultural hybrid; <sup>2</sup> previously known as *E. passgalense*

**Supplementary Table S2.**
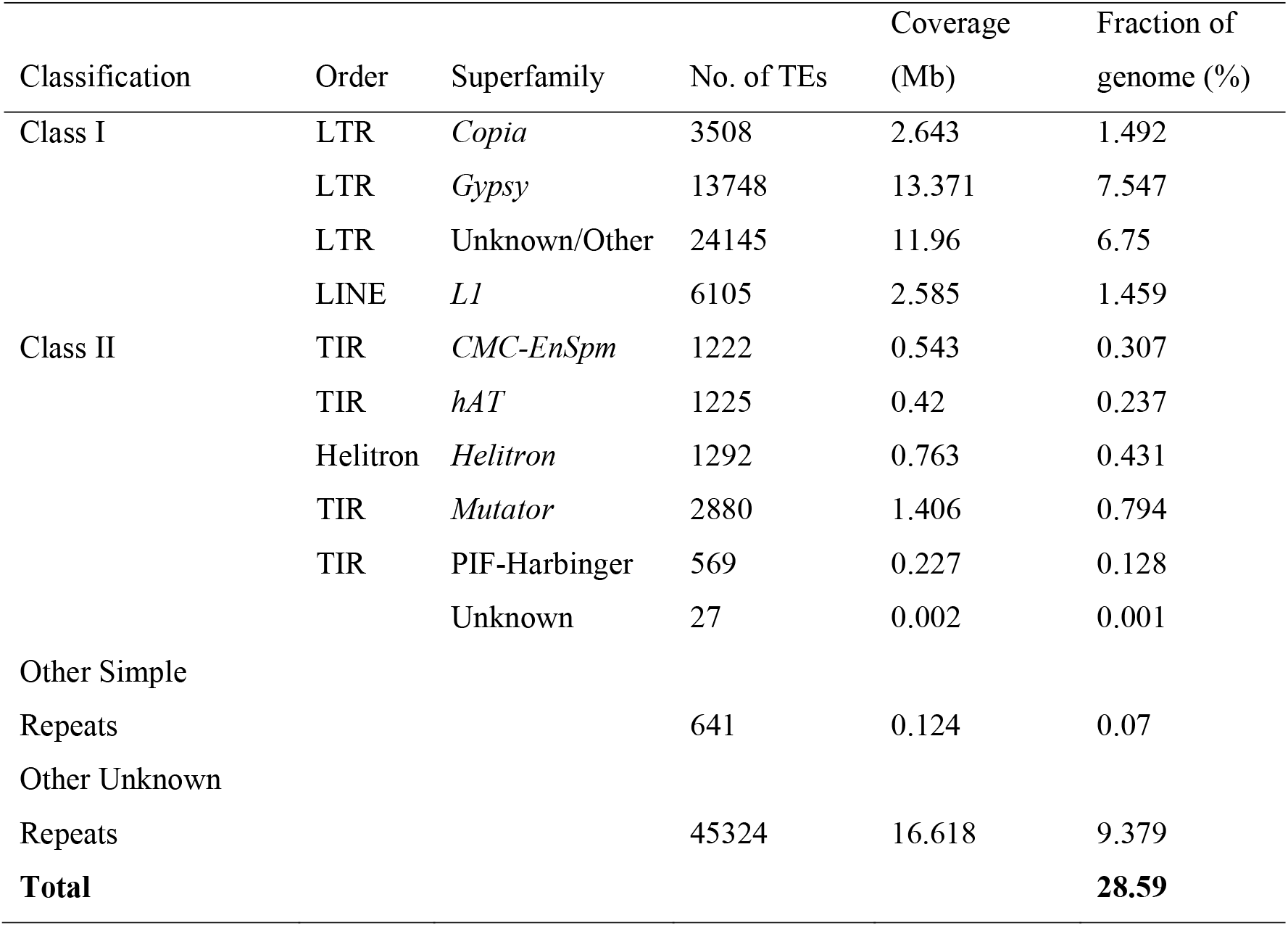
Repetitive sequences and transposable elements in the *E. cheiranthoides* genome.

| Classification | Order | Superfamily | No. of TEs | Coverage<br>(Mb) | Fraction of<br>genome (%) |
| --- | --- | --- | --- | --- | --- |
| Class I | LTR | *Copia* | 3508 | 2.643 | 1.492 |
|  | LTR | *Gypsy* | 13748 | 13.371 | 7.547 |
|  | LTR | Unknown/Other | 24145 | 11.96 | 6.75 |
|  | LINE | *L1* | 6105 | 2.585 | 1.459 |
| Class II | TIR | *CMC-EnSpm* | 1222 | 0.543 | 0.307 |
|  | TIR | *hAT* | 1225 | 0.42 | 0.237 |
|  | Helitron | *Helitron* | 1292 | 0.763 | 0.431 |
|  | TIR | *Mutator* | 2880 | 1.406 | 0.794 |
|  | TIR | PIF-Harbinger | 569 | 0.227 | 0.128 |
|  |  | Unknown | 27 | 0.002 | 0.001 |
| Other Simple |  |  |  |  |  |
| Repeats |  |  | 641 | 0.124 | 0.07 |
| Other Unknown |  |  |  |  |  |
| Repeats |  |  | 45324 | 16.618 | 9.379 |
| **Total** |  |  |  |  | **28.59** |

**Supplementary Table S3.** Transcriptome assembly metrics, including number of sequences, N50 values, and recovered BUSCO gene number. Additionally, transcript lengths were divided by the length of the top BLAST match to the *E. cheiranthoides* v1.1 gene model (EC1.1) to determine fragmentation of the transcriptome assemblies (tophit average trinity_len/EC_len). RNA sequences from each of the 48 *Erysimum* species were mapped to the *E. cheiranthoides* genome, and results are reported as the number of E. cheiranthoides genes represented and the mapping percentage.

| Species | Sequence count | N50 [bp] | BUSCO genes |  |  |  |  | Tophit average trinity_len/EC_len | No. of EC1.1 genes represented | Mapping to EC1.1 (%) |
| --- | --- | --- | --- | --- | --- | --- | --- | --- | --- | --- |
|  |  |  | Complete (percent total) | Complete single-copy | Complete duplicated | Fragmented | Missing |  |  |  |
| ALI | 207422 | 595 | 1010 (70.1%) | 276 | 734 | 250 | 180 | 0.74 | 19150 | 56.6 |
| AMO | 217430 | 888 | 903 (62.7%) | 419 | 484 | 323 | 214 | 0.97 | 21450 | 59.4 |
| AND | 165687 | 1172 | 1043 (72.4%) | 445 | 598 | 252 | 145 | 1.09 | 20995 | 58.5 |
| AUC | 164506 | 1367 | 1163 (80.8%) | 439 | 724 | 183 | 94 | 1.21 | 20711 | 58.6 |
| BAE | 180487 | 1127 | 1041 (72.3%) | 479 | 562 | 253 | 146 | 1.05 | 21187 | 59.1 |
| BAS | 135035 | 1420 | 1183 (82.2%) | 442 | 741 | 147 | 110 | 0.91 | 20279 | 57.8 |
| BIC | 99234 | 1868 | 1314 (91.3%) | 519 | 795 | 67 | 59 | 1.13 | 19925 | 57.8 |
| CAP | 260998 | 842 | 887 (61.6%) | 406 | 481 | 366 | 187 | 0.97 | 22113 | 57.4 |
| CHR | 93525 | 1963 | 1350 (93.8%) | 565 | 785 | 39 | 51 | 1.22 | 20213 | 57.3 |
| CRA | 143666 | 1582 | 1266 (87.9%) | 417 | 849 | 113 | 61 | 1.01 | 20924 | 54.2 |
| CRE | 102472 | 1768 | 1321 (91.7%) | 589 | 732 | 55 | 64 | 1.28 | 19686 | 59 |
| CSS | 118038 | 1588 | 1241 (86.2%) | 469 | 773 | 90 | 108 | 1.01 | 20457 | 56.4 |
| CUS | 122476 | 1644 | 1263 (87.7%) | 457 | 806 | 110 | 67 | 1.02 | 20077 | 58.1 |
| DIF | 139288 | 770 | 1189 (82.6%) | 392 | 797 | 154 | 97 | 0.89 | 20472 | 56.7 |
| ECE | 81984 | 2139 | 1341 (93.1%) | 658 | 683 | 34 | 65 | 1.41 | 19519 | 94.8 |
| ER1 | 223508 | 900 | 951 (66.0%) | 378 | 573 | 287 | 202 | 1.01 | 21335 | 66.5 |
| ER2 | 258578 | 715 | 785 (54.5%) | 307 | 478 | 379 | 276 | 0.57 | 23080 | 55.1 |
| ER3 | 123653 | 1886 | 1338 (92.9%) | 452 | 886 | 40 | 62 | 1.35 | 20130 | 77.5 |
| ER4 | 89871 | 1844 | 1328 (92.2%) | 469 | 859 | 49 | 63 | 1.13 | 23371 | 78.2 |
| FIZ | 94064 | 1956 | 1320 (91.7%) | 551 | 769 | 50 | 70 | 1.21 | 21130 | 59.1 |
| FRA | 220291 | 1004 | 992 (68.9%) | 384 | 608 | 284 | 164 | 1.04 | 21412 | 59.9 |
| HIE | 218565 | 955 | 960 (66.7%) | 296 | 664 | 286 | 194 | 0.71 | 20878 | 66 |
| HOR | 109045 | 1784 | 1306 (90.7%) | 501 | 805 | 77 | 57 | 1.29 | 19758 | 61.9 |
| HUN | 228679 | 881 | 903 (62.7%) | 291 | 612 | 313 | 224 | 0.67 | 21558 | 61.4 |
| INC | 90295 | 2002 | 1337 (92.8%) | 481 | 856 | 40 | 63 | 1.23 | 20671 | 61.4 |
| KOT | 104476 | 1755 | 1322 (91.8%) | 531 | 791 | 56 | 62 | 1.07 | 21558 | 56.8 |
| LAG | 155771 | 1422 | 1196 (83.1%) | 419 | 777 | 153 | 91 | 1.21 | 20777 | 59.1 |
| MAJ | 174858 | 1027 | 958 (66.5%) | 462 | 496 | 286 | 196 | 1.01 | 20830 | 57.9 |
| MED | 106039 | 1869 | 1301 (90.3%) | 506 | 795 | 69 | 70 | 1.41 | 20907 | 57.9 |
| MEX | 102500 | 1115 | 1284 (89.2%) | 501 | 783 | 74 | 82 | 1.13 | 21693 | 57.4 |
| MEZ | 187227 | 1160 | 1074 (74.6%) | 420 | 654 | 203 | 163 | 1.1 | 20811 | 56.9 |
| MIC | 115738 | 1696 | 1271 (88.3%) | 513 | 758 | 94 | 75 | 1.29 | 19993 | 61.2 |
| NAX | 104553 | 2016 | 1351 (93.8%) | 549 | 802 | 32 | 57 | 1.37 | 19589 | 60.5 |
| NER | 161135 | 1485 | 1228 (85.3%) | 471 | 757 | 133 | 79 | 1.23 | 20833 | 59.4 |
| NEV | 115096 | 1710 | 1282 (89.0%) | 450 | 832 | 84 | 74 | 1.08 | 19160 | 59.6 |
| ODO | 193451 | 1087 | 975 (67.7%) | 421 | 554 | 289 | 176 | 1.07 | 21108 | 57.9 |
| PIE | 238425 | 878 | 887 (61.6%) | 275 | 612 | 327 | 226 | 0.69 | 22492 | 59.5 |
| PSE | 129384 | 1513 | 1230 (85.4%) | 428 | 802 | 134 | 76 | 0.95 | 22784 | 57.4 |
| PUL | 251705 | 741 | 773 (53.7%) | 305 | 468 | 391 | 276 | 0.58 | 20633 | 54.6 |
| REP | 64315 | 2160 | 1340 (93.1%) | 727 | 613 | 34 | 66 | 1.33 | 22932 | 57.3 |
| RHA | 220584 | 876 | 887 (61.6%) | 440 | 447 | 335 | 218 | 0.97 | 21715 | 57.4 |
| RUS | 154276 | 1328 | 1125 (78.1%) | 470 | 655 | 188 | 127 | 1.13 | 20616 | 60.7 |
| SCO | 92945 | 1897 | 1331 (92.4%) | 625 | 706 | 48 | 61 | 1.34 | 19314 | 61.2 |
| SEM | 91502 | 1875 | 1317 (91.5%) | 608 | 709 | 51 | 72 | 1.34 | 19479 | 59.9 |
| SYL | 113269 | 1918 | 1331 (92.4%) | 449 | 882 | 44 | 65 | 1.37 | 20013 | 77.8 |
| VIR | 212293 | 574 | 990 (68.8%) | 305 | 685 | 278 | 172 | 0.72 | 22901 | 66.1 |
| WIC | 91007 | 2039 | 1339 (93.0%) | 586 | 753 | 36 | 65 | 1.28 | 23344 | 56.9 |
| WTT | 171465 | 1768 | 1284 (89.2%) | 395 | 889 | 97 | 59 | 1.37 | 21116 | 57.8 |

**Supplementary Table S4.**
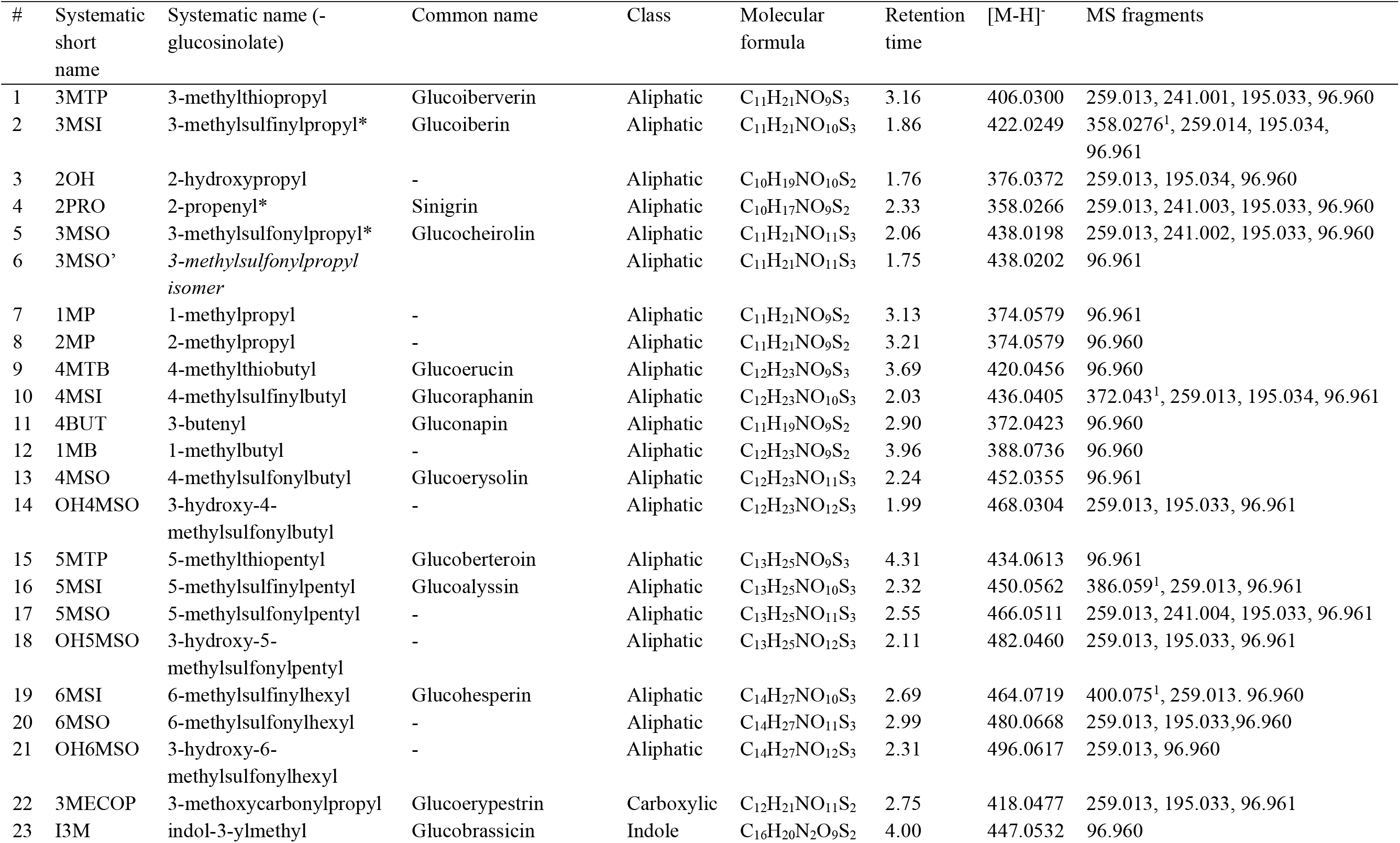
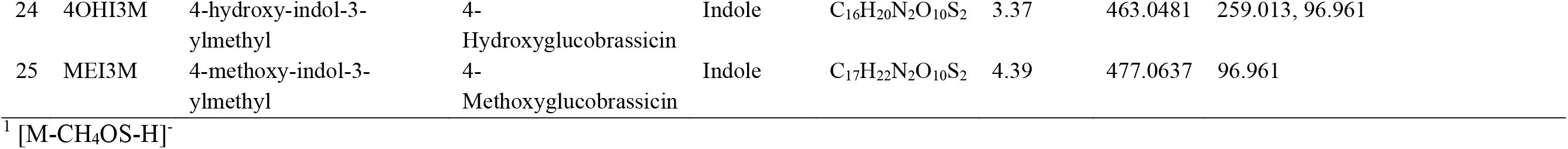
List of glucosinolate compounds, determined by exact mass, fragmentation patterns, and retention time. Asterisks (*) indicate compounds confirmed by commercial standards.

**Supplementary Table S5.**
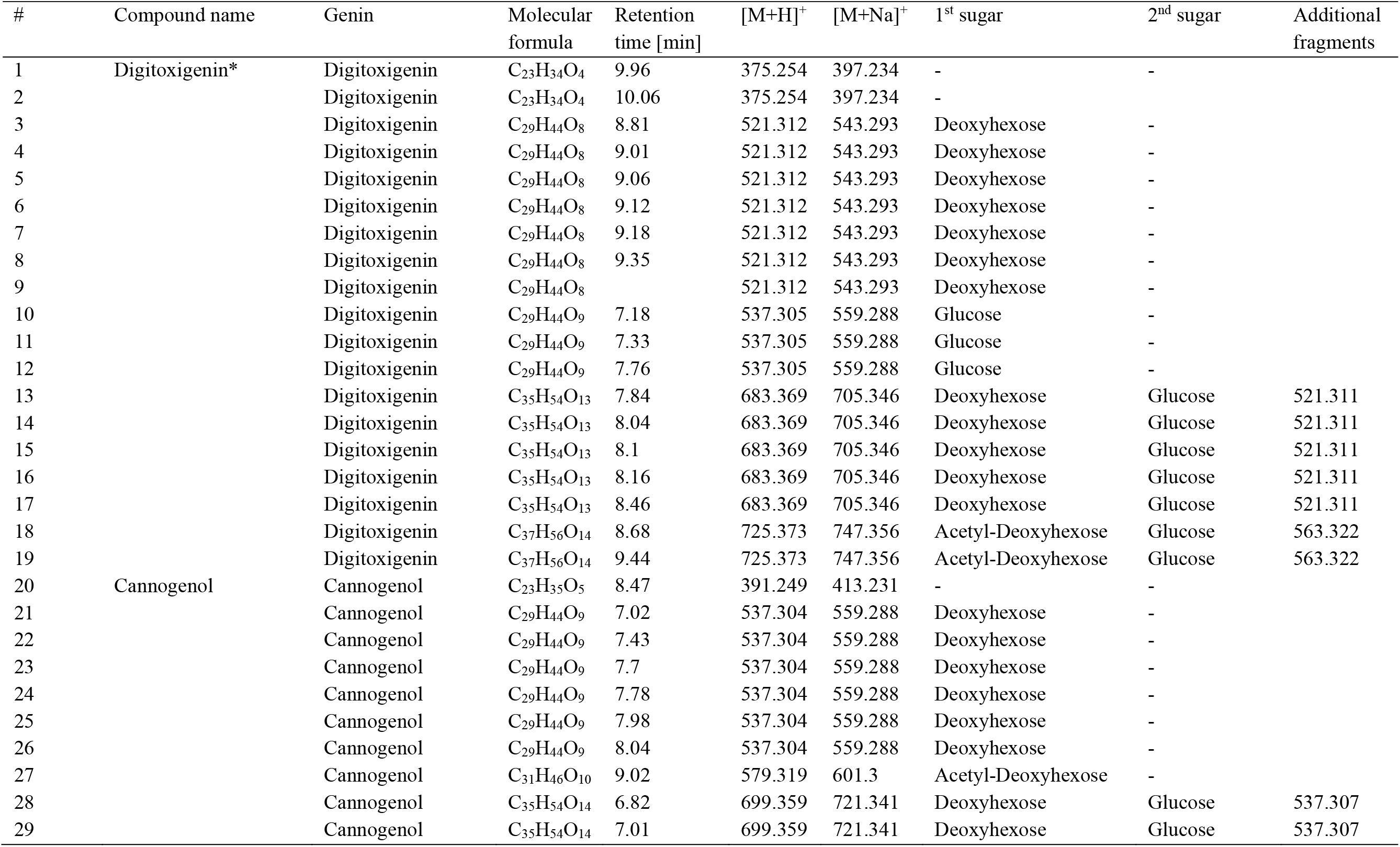
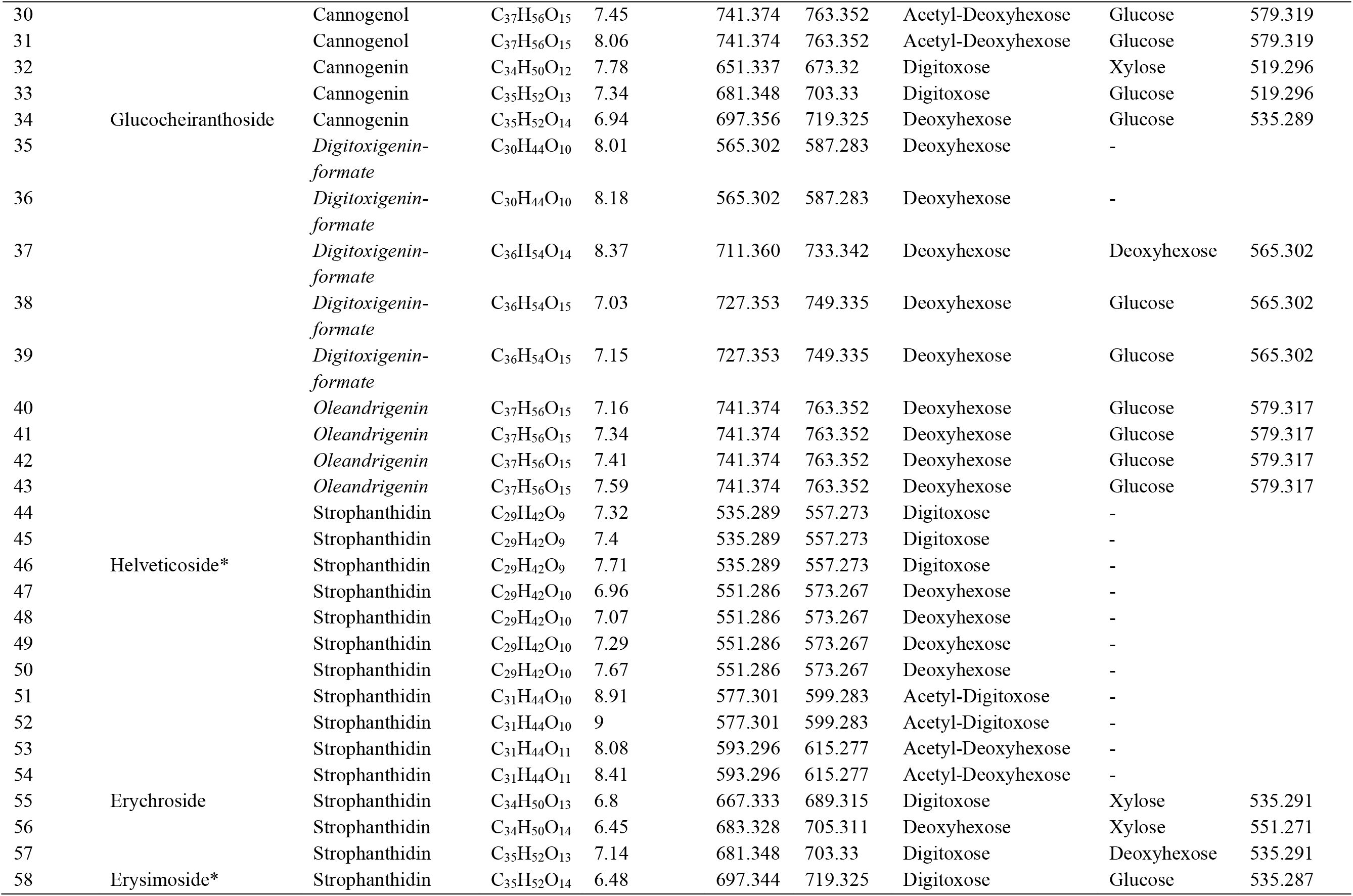
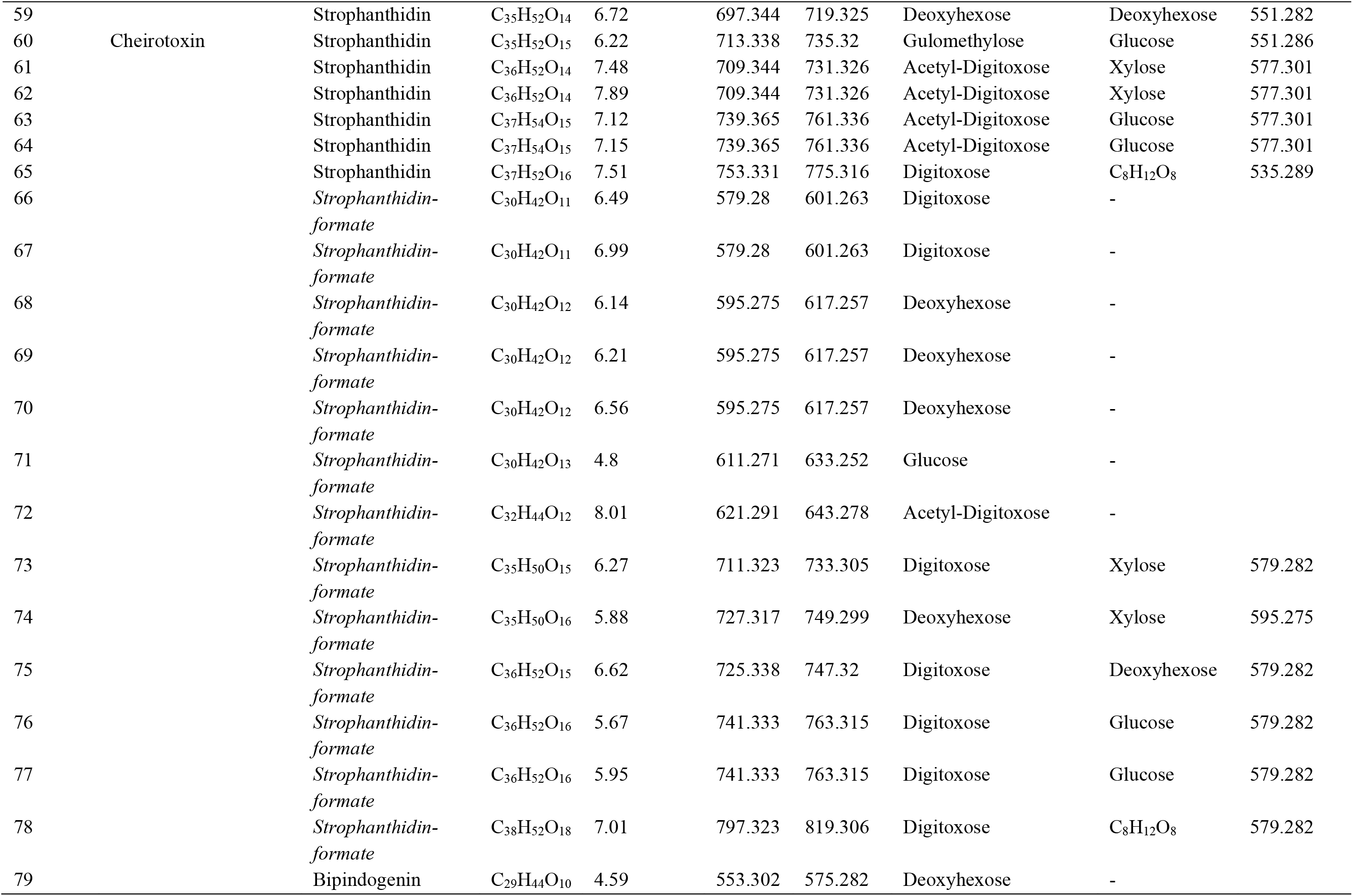
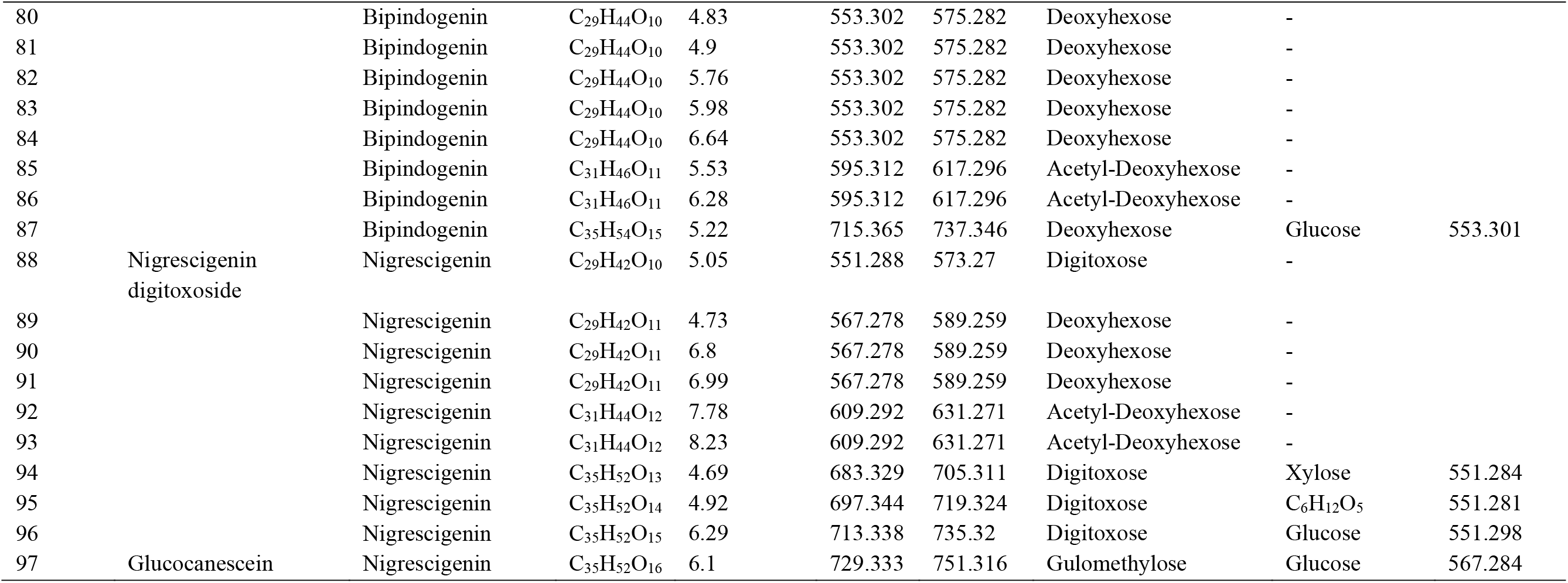
List of candidate cardenolide compounds, determined by exact mass and fragmentation patterns. Asterisks (*) indicate compounds confirmed by commercial standards. Compounds #65 and #78 were excluded due to potential artefact formation with formic acid (see Figure S12).

## Notes

https://www.erysimum.org

## References

Abdelaziz, M., J. Lorite, A. J. Munoz-Pajares, M. B. Herrador, F. Perfectti, and J. M. Gomez. 2011. Using complementary techniques to distinguish cryptic species: a new *Erysimum* (Brassicaceae) species from North Africa. American Journal of Botany 98:1049–1060.

Abdelaziz, M., A. J. Muñoz-Pajares, J. Lorite, M. B. Herrador, F. Perfectti, and J. M. Gómez. 2014. Phylogenetic relationships of Erysimum (Brassicaceae) from the Baetic Mountains (SE Iberian Peninsula). Anales del Jardín Botánico de Madrid 71:e005.

Agrawal, A. A. 2005. Natural selection on common milkweed (*Asclepias syriaca*) by a community of specialized insect herbivores. Evolutionary Ecology Research 7:651–667.

Agrawal, A. A. and M. Fishbein. 2008. Phylogenetic escalation and decline of plant defense strategies. Proceedings of the National Academy of Sciences of the United States of America 105:10057–10060.

Agrawal, A. A., G. Petschenka, R. A. Bingham, M. G. Weber, and S. Rasmann. 2012. Toxic cardenolides: chemical ecology and coevolution of specialized plant-herbivore interactions. New Phytologist 194:28–45.

Al-Shehbaz, I. A. 1988. The genera of *Anchonieae* (Hesperideae) (Cruciferae; Brassicaceae) in the southeastern United States. Journal of the Arnold Arboretum 69:193–212.

Al-Shehbaz, I. A. 2010. Erysimum Linnaeus. Pages 534-545 in N. R. Morin, editor. Flora of North America North of Mexico. Oxford University Press, New York.

Altschul, S. F., W. Gish, W. Miller, E. W. Myers, and D. J. Lipman. 1990. Basic local alignment search tool. Journal of Molecular Biology 215:403–410.

Andersson, D., R. Chakrabarty, S. Bejai, J. M. Zhang, L. Rask, and J. Meijer. 2009. Myrosinases from root and leaves of *Arabidopsis thaliana* have different catalytic properties. Phytochemistry 70:1345–1354.

Barth, C. and G. Jander. 2006. *Arabidopsis* myrosinases TGG1 and TGG2 have redundant function in glucosinolate breakdown and insect defense. Plant Journal 46:549–562.

Bednarek, P., M. Pislewska-Bednarek, A. Svatos, B. Schneider, J. Doubsky, M. Mansurova, M. Humphry, C. Consonni, R. Panstruga, A. Sanchez-Vallet, A. Molina, and P. Schulze-Lefert. 2009. A glucosinolate metabolism pathway in living plant cells mediates broad-spectrum antifungal defense. Science 323:101–106.

Bidart-Bouzat, M. G. and D. J. Kliebenstein. 2008. Differential levels of insect herbivory in the field associated with genotypic variation in glucosinolates in *Arabidopsis thaliana*. Journal of Chemical Ecology 34:1026–1037.

Bingham, R. A. and A. A. Agrawal. 2010. Specificity and trade-offs in the induced plant defence of common milkweed *Asclepias syriaca* to two lepidopteran herbivores. Journal of Ecology 98:1014–1022.

Blomberg, S. P., T. Garland, and A. R. Ives. 2003. Testing for phylogenetic signal in comparative data: behavioral traits are more labile. Evolution 57:717–745.

Boutet, E., D. Lieberherr, M. Tognolli, M. Schneider, and A. Bairoch. 2007. UniProtKB/Swiss-Prot. Pages 89-112 in D. Edwards, editor. Plant Bioinformatics: Methods and Protocols. Humana Press.

Brock, A., T. Herzfeld, R. Paschke, M. Koch, and B. Draeger. 2006. Brassicaceae contain nortropane alkaloids. Phytochemistry 67:2050–2057.

Campbell, M. S., M. Y. Law, C. Holt, J. C. Stein, G. D. Moghe, D. E. Hufnagel, J. K. Lei, R. Achawanantakun, D. Jiao, C. J. Lawrence, D. Ware, S. H. Shiu, K. L. Childs, Y. N. Sun, N. Jiang, and M. Yandell. 2014. MAKER-P: a tool kit for the rapid creation, management, and quality control of plant genome annotations. Plant Physiology 164:513–524.

Campos, M. L., Y. Yoshida, I. T. Major, D. D. Ferreira, S. M. Weraduwage, J. E. Froehlich, B. F. Johnson, D. M. Kramer, G. Jander, T. D. Sharkey, and G. A. Howe. 2016. Rewiring of jasmonate and phytochrome B signalling uncouples plant growth-defense tradeoffs. Nature Communications 7.

Cantarel, B. L., I. Korf, S. M. C. Robb, G. Parra, E. Ross, B. Moore, C. Holt, A. S. Alvarado, and M. Yandell. 2008. MAKER: an easy-to-use annotation pipeline designed for emerging model organism genomes. Genome Research 18:188–196.

Cataldi, T. R. I., F. Lelario, D. Orlando, and S. A. Bufo. 2010. Collision-induced dissociation of the A+2 isotope ion facilitates glucosinolates structure elucidation by electrospray ionization-tandem mass spectrometry with a linear quadrupole ion trap. Analitical Chemistry 82:5686–5696.

Chen, W., S. Shakir, M. Bigham, A. Richter, Z. Feiz, and G. Jander. 2019. Genome sequence of the corn leaf aphid (*Rhopalosiphum maidis* Fitch). GigaScience 8:giz033.

Chew, F. S. 1975. Coevolution of pierid butterflies and their cruciferous foodplants. 1. Relative quality of available resources. Oecologia 20:117-127.

Chew, F. S. 1977. Coevolution of pierid butterflies and their cruciferous foodplants. 2. The distribution of eggs on potential foodplants. Evolution 31:568-579.

Chin, C. S., P. Peluso, F. J. Sedlazeck, M. Nattestad, G. T. Concepcion, A. Clum, C. Dunn, R. O’Malley, R. Figueroa-Balderas, A. Morales-Cruz, G. R. Cramer, M. Delledonne, C. Y. Luo, J. R. Ecker, D. Cantu, D. R. Rank, and M. C. Schatz. 2016. Phased diploid genome assembly with single-molecule real-time sequencing. Nature Methods 13:1050–1054.

Chisholm, M. 1973. Biosynthesis of 3-methoxycarbonylpropyl-glucosinolate in an *Erysimum* species. Phytochemistry 12:605–608.

Clay, N. K., A. M. Adio, C. Denoux, G. Jander, and F. M. Ausubel. 2009. Glucosinolate metabolites required for an Arabidopsis innate immune response. Science 323:95–101.

Cole, R. A. 1976. Isothiocyanates, nitriles and thiocyanates as products of autolysis of glucosinolates in *Cruciferae*. Phytochemistry 15:759–762.

Cornell, H. V. and B. A. Hawkins. 2003. Herbivore responses to plant secondary compounds: A test of phytochemical coevolution theory. American Naturalist 161:507–522.

Dimock, M. B., J. A. A. Renwick, C. D. Radke, and K. Sachdev-Gupta. 1991. Chemical constituents of an unacceptable crucifer, *Erysimum cheiranthoides*, deter feeding by Pieris rapae. Journal of Chemical Ecology 17:525–533.

Duffey, S. S. 1980. Sequestration of plant natural products by insects. Annual Review of Entomology 25:447–477.

Dzimiri, N., U. Fricke, and W. Klaus. 1987. Influence of derivation on the lipophilicity and inhibitory actions of cardiac glycosides on myocardial Na^+^-K^+^-ATPase. British journal of pharmacology 91:31–38.

Ellinghaus, D., S. Kurtz, and U. Willhoeft. 2008. LTRharvest, an efficient and flexible software for de novo detection of LTR retrotransposons. BMC Bioinformatics 9:18.

Emms, D. M. and S. Kelly. 2015. OrthoFinder: solving fundamental biases in whole genome comparisons dramatically improves orthogroup inference accuracy. Genome Biology 16.

English, A. C., S. Richards, Y. Han, M. Wang, V. Vee, J. X. Qu, X. Qin, D. M. Muzny, J. G. Reid, K. C. Worley, and R. A. Gibbs. 2012. Mind the gap: upgrading genomes with Pacific Biosciences RS long-read sequencing technology. Plos One 7:e47768.

Erthmann, P. O., N. Agerbirk, and S. Bak. 2018. A tandem array of UDP-glycosyltransferases from the UGT73C subfamily glycosylate sapogenins, forming a spectrum of mono- and bisdesmosidic saponins. Plant Molecular Biology 97:37–55.

Fahey, J. W., A. T. Zalcmann, and P. Talalay. 2001. The chemical diversity and distribution of glucosinolates and isothiocyanates among plants. Phytochemistry 56:5–51.

Feeny, P. 1977. Defensive ecology of the Cruciferae. Annals of the Missouri Botanical Garden 64:221–234.

Feliner, G. N. and J. A. Rossello. 2007. Better the devil you know? Guidelines for insightful utilization of nrDNA ITS in species-level evolutionary studies in plants. Molecular Phylogenetics and Evolution 44:911–919.

Firn, R. D. and C. G. Jones. 2003. Natural products - a simple model to explain chemical diversity. Natural Product Reports 20:382–391.

Forbey, J. S., M. D. Dearing, E. M. Gross, C. M. Orians, E. E. Sotka, and W. J. Foley. 2013. A pharm-ecological perspective of terrestrial and aquatic plant-herbivore interactions. Journal of Chemical Ecology 39:465–480.

Fraenkel, G. S. 1959. The raison d’être of secondary plant substances. Science 129:1466–1470.

Frisch, T. and B. L. Møller. 2012. Possible evolution of alliarinoside biosynthesis from the glucosinolate pathway in *Alliaria petiolata*. Febs Journal 279:1545–1562.

German, D. A. 2014. Notes on taxonomy of Erysimum (Erysimeae, Cruciferae) of Russia and adjacent states. 1. Erysimum collinum and Erysimum hajastanicum. Turczaninowia 17:10–32.

Gershenzon, J., A. Fontana, M. Burow, U. Wittstock, and J. Degenhardt. 2012. Mixtures of plant secondary metabolites: metabolic origins and ecological benefits.in G. R. Iason, M. Dicke, and S. E. Hartley, editors. The ecology of plant secondary metabolites. Cambridge University Press, Cambridge.

Gómez, J. M. 2005. Non-additive effects of herbivores and pollinators on *Erysimum mediohispanicum* (Cruciferae) fitness. Oecologia 143:412–418.

Gómez, J. M., F. Perfectti, and C. P. Klingenberg. 2014. The role of pollinator diversity in the evolution of corolla-shape integration in a pollination-generalist plant clade. Philosophical Transactions of the Royal Society B 369:20130257.

Gómez, J. M., F. Perfectti, and J. Lorite. 2015. The role of pollinators in floral diversification in a clade of generalist flowers. Evolution 69:863–878.

Goodstein, D. M., S. Shu, R. Howson, R. Neupane, R. D. Hayes, J. Fazo, T. Mitros, W. Dirks, U. Hellsten, N. Putnam, and D. S. Rokhsar. 2011. Phytozome: a comparative platform for green plant genomics. Nucleic Acids Research 40:D1178–D1186.

Gordon, S. P., E. Tseng, A. Salamov, J. W. Zhang, X. D. Meng, Z. Y. Zhao, D. W. Kang, J. Underwood, I. V. Grigoriev, M. Figueroa, J. S. Schilling, F. Chen, and Z. Wang. 2015. Widespread polycistronic transcripts in fungi revealed by single-molecule mRNA sequencing. Plos One 10:e0132628.

Haas, B. J., A. Papanicolaou, M. Yassour, M. Grabherr, P. D. Blood, J. Bowden, M. B. Couger, D. Eccles, B. Li, M. Lieber, M. D. MacManes, M. Ott, J. Orvis, N. Pochet, F. Strozzi, N. Weeks, R. Westerman, T. William, C. N. Dewey, R. Henschel, R. D. Leduc, N. Friedman, and A. Regev. 2013. De novo transcript sequence reconstruction from RNA-seq using the Trinity platform for reference generation and analysis. Nature Protocols 8:1494–1512.

Halkier, B. A. and J. Gershenzon. 2006. Biology and biochemistry of glucosinolates. Annual Review of Plant Biology 57:303–333.

Haribal, M. and J. A. A. Renwick. 2001. Seasonal and population variation in flavonoid and alliarinoside content of *Alliaria petiolata*. Journal of Chemical Ecology 27:1585–1594.

Harmon, L. J., J. T. Weir, C. D. Brock, R. E. Glor, and W. Challenger. 2008. GEIGER: investigating evolutionary radiations. Bioinformatics 24:129–131.

Havko, N. E., I. T. Major, J. B. Jewell, E. Attaran, J. Browse, and G. A. Howe. 2016. Control of carbon assimilation and partitioning by jasmonate: an accounting of growth–defense tradeoffs. Plants 5:7.

Hegnauer, R. 1964. Chemotaxonomie der Pflanzen III. Birkhäuser Verlag, Basel, Switzerland.

Huang, C. H., R. R. Sun, Y. Hu, L. P. Zeng, N. Zhang, L. M. Cai, Q. Zhang, M. A. Koch, I. Al-Shehbaz, P. P. Edger, J. C. Pires, D. Y. Tan, Y. Zhong, and H. Ma. 2016. Resolution of Brassicaceae phylogeny using nuclear genes uncovers nested radiations and supports convergent morphological evolution. Molecular Biology and Evolution 33:394-412.

Huang, X., J. A. A. Renwick, and K. Sachdevgupta. 1993. A chemical basis for differential acceptance of *Erysimum cheiranthoides* by two *Pieris* species. Journal of Chemical Ecology 19:195–210.

Iason, G. R., J. M. O’Reilly-Wapstra, M. J. Brewer, R. W. Summers, and B. Moore. 2011. Do multiple herbivores maintain chemical diversity of Scots pine monoterpenes? Philosophical Transactions of the Royal Society B: Biological Sciences 366:1337–1345.

Illumina. 2017. Effects of index misassignment on multiplexing and downstream analysis. Illumina Whitepapers.

Jaretzky, R. and M. Wilcke. 1932. Die herzwirksamen Glykoside von Cheiranthus cheiri und verwandten Arten. Archiv der Pharmazie 270:81–94.

Jones, P., D. Binns, H. Y. Chang, M. Fraser, W. Z. Li, C. McAnulla, H. McWilliam, J. Maslen, A. Mitchell, G. Nuka, S. Pesseat, A. F. Quinn, A. Sangrador-Vegas, M. Scheremetjew, S. Y. Yong, R. Lopez, and S. Hunter. 2014. InterProScan 5: genome-scale protein function classification. Bioinformatics 30:1236–1240.

Katoh, K., K. Misawa, K. Kuma, and T. Miyata. 2002. MAFFT: a novel method for rapid multiple sequence alignment based on fast Fourier transform. Nucleic Acids Research 30:3059–3066.

Katz, E., S. Nisani, B. S. Yadav, M. G. Woldemariam, B. Shai, U. Obolski, M. Ehrlich, E. Shani, G. Jander, and D. A. Chamovitz. 2015. The glucosinolate breakdown product indole-3-carbinol acts as an auxin antagonist in roots of *Arabidopsis thaliana*. The Plant Journal 82:547–555.

Kerwin, R., J. Feusier, J. Corwin, M. Rubin, C. Lin, A. Muok, B. Larson, B. Li, B. Joseph, M. Francisco, D. Copeland, C. Weinig, and D. J. Kliebenstein. 2015. Natural genetic variation in *Arabidopsis thaliana* defense metabolism genes modulates field fitness. eLife 4:1–28.

Kim, D., B. Landmead, and S. L. Salzberg. 2015. HISAT: a fast spliced aligner with low memory requirements. Nature Methods 12:357–360.

Kim, J. H., B. W. Lee, F. C. Schroeder, and G. Jander. 2008. Identification of indole glucosinolate breakdown products with antifeedant effects on *Myzus persicae* (green peach aphid). Plant Journal 54:1015–1026.

Kjaer, A. and R. Gmelin. 1957. Isothiocyanates XXV: methyl 4-isothiocyanatobutyrate, a new mustard oil present as a glucoside (glucoerypestrin) in *Erysimum* species. Acta Chemica Scandinavica 11:577–578.

Klauck, D. and M. Luckner. 1995. In vitro measurement of digitalis-like compounds by inhibition of Na^+^/K^+^-ATPase: determination of the inhibitory effect. Pharmazie 50:395–399.

Kliebenstein, D. J., J. Kroymann, P. Brown, A. Figuth, D. Pedersen, J. Gershenzon, and T. Mitchell-Olds. 2001. Genetic control of natural variation in *Arabidopsis* glucosinolate accumulation. Plant Physiology 126:811–825.

Korf, I. 2004. Gene finding in novel genomes. BMC Bioinformatics 5:59.

Kreis, W. and F. Müller-Uri. 2010. Biochemistry of sterols, cardiac glycosides, brassinosteroids, phytoecdysteroids and steroid saponins. Pages 304-363 in M. Wink, editor. Biochemistry of plant secondary metabolism. CRC Press, Sheffield.

Krzywinski, M., J. Schein, I. Birol, J. Connors, R. Gascoyne, D. Horsman, S. J. Jones, and M. A. Marra. 2009. Circos: an information aesthetic for comparative genomics. Genome Research 19:1639–1645.

Kumar, S., G. Stecher, M. Li, C. Knyaz, and K. Tamura. 2018. MEGA X: molecular evolutionary genetics analysis across computing platforms. Molecular Biology and Evolution 35:1547–1549.

Kurtz, S., A. Phillippy, A. L. Delcher, M. Smoot, M. Shumway, C. Antonescu, and S. L. Salzberg. 2004. Versatile and open software for comparing large genomes. Genome Biology 5:R12.

Livshultz, T., E. Kaltenegger, S. C. K. Straub, K. Weitemier, E. Hirsch, K. Koval, L. Mema, and A. Liston. 2018. Evolution of pyrrolizidine alkaloid biosynthesis in Apocynaceae: revisiting the defence de-escalation hypothesis. New Phytologist 218:762–773.

Makarevich, I. F., K. V. Zhernoklev, T. V. Slyusarskaya, and G. N. Yarmolenko. 1994. Cardenolide-containing plants of the family Cruciferae. Chemistry of Natural Compounds 30:275–289.

Mapleson, D., L. Venturini, G. Kaithakottil, and D. Swarbreck. 2018. Efficient and accurate detection of splice junctions from RNA-seq with Portcullis. GigaScience 7:giy131.

Mithöfer, A. and W. Boland. 2012. Plant defense against herbivores: chemical aspects. Annual Review of Plant Biology 63:431–450.

Moazzeni, H., S. Zarre, B. E. Pfeil, Y. J. K. Bertrand, D. A. German, I. A. Al-Shehbaz, K. Mummenhoff, and B. Oxelman. 2014. Phylogenetic perspectives on diversification and character evolution in the species-rich genus *Erysimum* (Erysimeae; Brassicaceae) based on a densely sampled ITS approach. Botanical Journal of the Linnean Society 175:497–522.

Moore, B., R. L. Andrew, C. Külheim, and W. J. Foley. 2014. Explaining intraspecific diversity in plant secondary metabolites in an ecological context. New Phytologist 201:733–750.

Müller, C. 2009. Interactions between glucosinolate- and myrosinase-containing plants and the sawfly *Athalia rosae*. Phytochemistry Reviews 8:121–134.

Munkert, J., M. Ernst, F. Müller-Uri, and W. Kreis. 2014. Identification and stress-induced expression of three 3β-hydroxysteroid dehydrogenases from *Erysimum crepidifolium* Rchb. and their putative role in cardenolide biosynthesis. Phytochemistry 100:26–33.

Nagata, W., C. Tamm, and T. Reichstein. 1957. Die Glykoside von Erysimum crepidifolium H. G. L. Reichenbach. Helvetica Chimica Acta 40:41–61.

Nakano, R. T., M. Pislewska-Bednarek, K. Yamada, P. P. Edger, M. Miyahara, M. Kondo, C. Bottcher, M. Mori, M. Nishimura, P. Schulze-Lefert, I. Hara-Nishimura, and P. Bednarek. 2017. PYK10 myrosinase reveals a functional coordination between endoplasmic reticulum bodies and glucosinolates in *Arabidopsis thaliana*. Plant Journal 89:204–220.

Nielsen, J. K. 1978a. Host plant discrimination within Cruciferae: feeding responses of four leaf beetles (Coleoptera: Chrysomelidae) to glucosinolates, cucurbitacins, and cardenolides. Entomologia Experimentalis Et Applicata 24:41–54.

Nielsen, J. K. 1978b. Host plant selection of monophagous and oligophagous flea beetles feeding on crucifers. Entomologia Experimentalis Et Applicata 24:562–569.

Ou, S. J. and N. Jiang. 2018. LTR_retriever: a highly accurate and sensitive program for identification of long terminal repeat retrotransposons. Plant Physiology 176:1410–1422.

Pagel, M. 1999. Inferring the historical patterns of biological evolution. Nature 401:877–884.

Paradis, E. and K. Schliep. 2019. ape 5.0: an environment for modern phylogenetics and evolutionary analyses in R. Bioinformatics 35:526–528.

Petschenka, G., S. Fandrich, N. Sander, V. Wagschal, M. Boppré, and S. Dobler. 2013. Stepwise evolution of resistance to toxic cardenolides via genetic substitutions in the Na^+^/K^+^-ATPase of milkweed butterflies (Lepidoptera: Danaini). Evolution 67:2753–2761.

Petschenka, G., C. S. Fei, J. J. Araya, S. Schröder, B. N. Timmermann, and A. A. Agrawal. 2018. Relative selectivity of plant cardenolides for Na^+^/K^+^-ATPases from the monarch butterfly and non-resistant insects. Frontiers in Plant Science 9:1424.

Polatschek, A. 2011. Revision der Gattung Erysimum (Cruciferae), Teil 2: Georgien, Armenien, Azerbaidzan, Türkei, Syrien, Libanon, Israel, Jordanien, Irak, Iran, Afghanistan. Annalen des Naturhistorischen Museums in Wien – Serie B 112.

Polatschek, A. and S. Snogerup. 2002. Erysimum in A. Strid and K. G. Tan, editors. Flora Hellenica 2. Koeltz Scientific Books, Koenigstein.

Price, M. N., P. S. Dehal, and A. P. Arkin. 2010. FastTree 2 - approximately maximum-likelihood trees for large alignments. Plos One 5:e9490.

Quinlan, A. R. and I. M. Hall. 2010. BEDTools: a flexible suite of utilities for comparing genomic features. Bioinformatics 26:841–842.

Rask, L., E. Andreasson, B. Ekbom, S. Eriksson, B. Pontoppidan, and J. Meijer. 2000. Myrosinase: gene family evolution and herbivore defense in Brassicaceae. Plant Molecular Biology 42:93–113.

Rasmann, S. and A. A. Agrawal. 2011. Latitudinal patterns in plant defense: evolution of cardenolides, their toxicity and induction following herbivory. Ecology Letters 14:476–483.

Rasmann, S., M. D. Johnson, and A. A. Agrawal. 2009. Induced responses to herbivory and jasmonate in three milkweed species. Journal of Chemical Ecology 35:1326–1334.

Renwick, J. A. A., C. D. Radke, and K. Sachdevgupta. 1989. Chemical constituents of *Erysimum cheiranthoides* deterring oviposition by the cabbage butterfly, *Pieris rapae*. Journal of Chemical Ecology 15:2161–2169.

Revell, L. J. 2012. phytools: an R package for phylogenetic comparative biology (and other things). Methods in Ecology and Evolution 3:217–223.

Rhee, S. Y., P. Zhang, H. Foerster, and C. Tissier. 2006. AraCyc: Overview of an Arabidopsis metabolism database and its applications for plant research. Pages 141-154 in K. Saito, R. A. Dixon, and L. Willmitzer, editors. Biotechnology in Agriculture and Forestry. Springer, Berlin.

Richards, L. A., L. A. Dyer, M. L. Forister, A. M. Smilanich, C. D. Dodson, M. D. Leonard, and C. S. Jeffrey. 2015. Phytochemical diversity drives plant-insect community diversity. Proceedings of the National Academy of Sciences of the United States of America 112:10973–10978.

Richards, L. A., A. E. Glassmire, K. M. Ochsenrider, A. M. Smilanich, C. D. Dodson, C. S. Jeffrey, and L. A. Dyer. 2016. Phytochemical diversity and synergistic effects on herbivores. Phytochemistry Reviews 15:1153–1166.

Romeo, J. T., J. A. Saunders, and P. Barbosa, editors. 1996. Phytochemical diversity and redundancy in ecological interactions. Plenum Press, New York.

Sachdev-Gupta, K., C. D. Radke, J. A. A. Renwick, and M. B. Dimock. 1993. Cardenolides from *Erysimum cheiranthoides*: feeding deterrents to *Pieris rapae* larvae. Journal of Chemical Ecology 19:1355–1369.

Sachdev-Gupta, K., J. A. A. Renwick, and C. D. Radke. 1990. Isolation and identification of oviposition deterrents to cabbage butterfly, *Pieris rapae*, from *Erysimum cheiranthoides*. Journal of Chemical Ecology 16:1059–1067.

Salazar, D., J. Lokvam, I. Mesones, M. Vásquez Pilco, J. M. Ayarza Zuñiga, P. de Valpine, and P. V. A. Fine. 2018. Origin and maintenance of chemical diversity in a species-rich tropical tree lineage. Nature Ecology & Evolution 2:983–990.

Sedio, B. E., J. C. Rojas Echeverri, C. A. Boya P., and S. J. Wright. 2017. Sources of variation in foliar secondary chemistry in a tropical forest tree community. Ecology 98:616–623.

Sela, I., H. Ashkenazy, K. Katoh, and T. Pupko. 2015. GUIDANCE2: accurate detection of unreliable alignment regions accounting for the uncertainty of multiple parameters. Nucleic Acids Research 43:W7–W14.

Sherameti, I., Y. Venus, C. Drzewiecki, S. Tripathi, V. M. Dan, I. Nitz, A. Varma, F. M. Grundler, and R. Oelmuller. 2008. PYK10, a beta-glucosidase located in the endoplasmatic reticulum, is crucial for the beneficial interaction between *Arabidopsis thaliana* and the endophytic fungus *Piriformospora indica*. Plant Journal 54:428–439.

Shinoda, T., T. Nagao, M. Nakayama, H. Serizawa, M. Koshioka, H. Okabe, and A. Kawai. 2002. Identification of a triterpenoid saponin from a crucifer, *Barbarea vulgaris*, as a feeding deterrent to the diamondback moth, *Plutella xylostella*. Journal of Chemical Ecology 28:587–599.

Singh, B. and R. P. Rastogi. 1970. Cardenolides - glycosides and genins. Phytochemistry 9:315–331.

Smith, S. A. and C. W. Dunn. 2008. Phyutility: a phyloinformatics tool for trees, alignments and molecular data. Bioinformatics 24:715–716.

Stamatakis, A. 2014. RAxML version 8: a tool for phylogenetic analysis and post-analysis of large phylogenies. Bioinformatics 30:1312–1313.

Stanke, M., M. Diekhans, R. Baertsch, and D. Haussler. 2008. Using native and syntenically mapped cDNA alignments to improve de novo gene finding. Bioinformatics 24:637–644.

Steppuhn, A. and I. T. Baldwin. 2007. Resistance management in a native plant: nicotine prevents herbivores from compensating for plant protease inhibitors. Ecology Letters 10:499–511.

Suzuki, R. and H. Shimodaira. 2014. pvclust: hierarchical clustering with P-values via multiscale bootstrap resampling. Bioinformatics 22:1540–1542.

Taussky, H. H. and E. Shorr. 1953. A microcolorimetric method for the determination of inorganic phosphorus. Journal of Biological Chemistry 202:675–685.

Textor, S. and J. Gershenzon. 2009. Herbivore induction of the glucosinolate-myrosinase defense system: major trends, biochemical bases and ecological significance. Phytochemistry Reviews 8:149–170.

Travers-Martin, N., F. Kuhlmann, and C. Müller. 2008. Revised determination of free and complexed myrosinase activities in plant extracts. Plant Physiology and Biochemistry 46:506–516.

Van Bel, M., T. Diels, E. Vancaester, L. Kreft, A. Botzki, Y. Van de Peer, F. Coppens, and K. Vandepoele. 2017. PLAZA 4.0: an integrative resource for functional, evolutionary and comparative plant genomics. Nucleic Acids Research 46:D1190–D1196.

Venturini, L., S. Caim, G. G. Kaithakottil, D. L. Mapleson, and D. Swarbreck. 2018. Leveraging multiple transcriptome assembly methods for improved gene structure annotation. GigaScience 7:giy093.

Walker, B. J., T. Abeel, T. Shea, M. Priest, A. Abouelliel, S. Sakthikumar, C. A. Cuomo, Q. D. Zeng, J. Wortman, S. K. Young, and A. M. Earl. 2014. Pilon: an integrated tool for comprehensive microbial variant detection and genome assembly improvement. Plos One 9:e112963.

Washburn, J. D., J. C. Schnable, G. C. Conant, T. P. Brutnell, Y. Shao, Y. Zhang, M. Ludwig, G. Davidse, and J. C. Pires. 2017. Genome-guided phylo-transcriptomic methods and the nuclear phylogentic tree of the Paniceae grasses. Scientific Reports 7:13528.

Waterhouse, R. M., M. Seppey, F. A. Simao, M. Manni, P. Ioannidis, G. Klioutchnikov, E. V. Kriventseva, and E. M. Zdobnov. 2018. BUSCO applications from quality assessments to gene prediction and phylogenomics. Molecular Biology and Evolution 35:543–548.

Weber, M. G. and A. A. Agrawal. 2014. Defense mutualisms enhance plant diversification. Proceedings of the National Academy of Sciences of the United States of America 111:16442–16447.

Wiklund, C. and C. Åhrberg. 1978. Host plants, nectar source plants, and habitat selection of males and females of *Anthocharis cardamines* (Lepidoptera). Oikos 31:169–183.

Winde, I. and U. Wittstock. 2011. Insect herbivore counteradaptations to the plant glucosinolate-myrosinase system. Phytochemistry 72:1566–1575.

Wink, M. 2003. Evolution of secondary metabolites from an ecological and molecular phylogenetic perspective. Phytochemistry 64:3–19.

Xu, Z. and H. Wang. 2007. LTR_FINDER: an efficient tool for the prediction of full-length LTR retrotransposons. Nucleic Acids Research 35:W265–W268.

Zhang, C., M. Rabiee, E. Sayyari, and S. Mirarab. 2018. ASTRAL-III: polynomial time species tree reconstruction from partially resolved gene trees. BMC Bioinformatics 19:153.

Zhang, J. M., B. Pontoppidan, J. P. Xue, L. Rask, and J. Meijer. 2002. The third myrosinase gene TGG3 in *Arabidopsis thaliana* is a pseudogene specifically expressed in stamen and petal. Physiologia Plantarum 115:25–34.

Züst, T., M. Mirzaei, and G. Jander. 2018. *Erysimum cheiranthoides*, an ecological research system with potential as a genetic and genomic model for studying cardiac glycoside biosynthesis. Phytochemistry Reviews 17:1239–1251.

Züst, T., G. Petschenka, A. P. Hastings, and A. A. Agrawal. 2019. Toxicity of milkweed leaves and latex: chromatographic quantification versus biological activity of cardenolides in 16 *Asclepias* species. Journal of Chemical Ecology 45:50–60.

